# CCDC186 controls dense-core vesicle cargo sorting by exit

**DOI:** 10.1101/616458

**Authors:** Jérôme Cattin-Ortolá, Irini Topalidou, Ho-Tak Lau, Blake Hummer, Cedric S. Asensio, Shao-En Ong, Michael Ailion

**Affiliations:** Department of Biochemistry, University of Washington, Seattle, United States; Department of Pharmacology, University of Washington, Seattle, United States; Department of Biological Sciences, University of Denver, Denver, United States

## Abstract

The regulated release of peptide hormones depends on their packaging into dense-core vesicles (DCVs). Two models have been proposed for DCV cargo sorting. The “sorting by entry” model proposes that DCV cargos selectively enter nascent DCVs at the trans-Golgi network (TGN). The “sorting by exit” model proposes that sorting occurs by the post-TGN removal of non-DCV cargos and retention of mature DCV cargos. Here we show that the coiled-coil protein CCDC186 controls sorting by exit. *Ccdc186* KO insulinoma cells secrete less insulin, fail to retain insulin and carboxypeptidase E in mature DCVs at the cell periphery, and fail to remove carboxypeptidase D from immature DCVs. A mutation affecting the endosome-associated recycling protein (EARP) complex causes similar defects in DCV cargo retention and removal. CCDC186 and EARP may act together to control the post-Golgi retention of cargos in mature DCVs.

## INTRODUCTION

Neurons and endocrine cells contain specialized secretory vesicles called dense-core vesicles (DCVs) that undergo regulated secretion in response to external stimuli. DCVs contain neuropeptides, growth factors, biogenic amines, and peptide hormones that control numerous biological processes, ranging from growth and metabolism to mental state. DCVs are generated at the trans-Golgi network (TGN) as immature DCVs (iDCVs) and undergo a maturation process that includes vesicle acidification, proteolytic processing of cargos, and the acquisition of proper compartmental identity and release competency (Borgonovo et al., 2006; Gondré-Lewis et al., 2012; Kim et al., 2006; Tooze et al., 2001). Peptide precursors are processed in acidic iDCVs by several distinct processing enzymes. For example, the insulin precursor proinsulin is processed into mature insulin via two proprotein convertase enzymes (PC1/3 and PC2) and carboxypeptidase E (CPE) (Steiner et al., 2011).

Two models of DCV cargo sorting have been proposed. The ‘sorting by entry’ model proposes that cargos are sorted in the trans-Golgi and preferentially enter nascent DCVs as they bud off; the ‘sorting by exit’ model proposes that cargos are sorted in a post-Golgi step in which non-DCV cargos are removed from iDCVs while mature DCV cargos are retained (Arvan and Castle, 1998; Tooze, 1998). Both mechanisms likely contribute to DCV cargo sorting. Cargos removed from iDCVs include the transmembrane proteins carboxypeptidase D (CPD) and the mannose 6-phosphate receptors (CD-MPR and CI-MPR) (Klumperman et al., 1998; Varlamov et al., 1999).

Genetic studies in the nematode *C. elegans* have identified several molecules that function in neuronal DCV biogenesis, including the small G protein RAB-2, the RAB-2 effector CCCP-1 (the *C. elegans* homolog of CCDC186/C10orf118), the endosome-associated recycling protein (EARP) complex (VPS-50, VPS-51, VPS-52, and VPS-53), and the EARP interactor EIPR-1 (Ailion et al., 2014; Edwards et al., 2009; Paquin et al., 2016; Sumakovic et al., 2009; Topalidou et al., 2016). Loss of any of these proteins causes a similar reduction in the levels of cargos in mature DCVs in axons, but it is unclear how RAB-2 and CCCP-1/CCDC186 are connected to EARP. In both *C. elegans* and in the rat insulinoma 832/13 cell line, RAB-2 and CCCP-1/CCDC186 colocalize near the TGN where the early steps of DCV biogenesis take place (Ailion et al., 2014; Cattin-Ortolá et al., 2017). EARP is localized to two distinct compartments in 832/13 cells, a recycling endosome compartment and a compartment near the TGN where it colocalizes with CCDC186 (Topalidou, Cattin-Ortolá, et al., 2018). Additionally, *Eipr1* knockout 832/13 cells have reduced insulin secretion and an altered distribution of insulin in the cell, with relatively more insulin near the TGN and less at the cell periphery (Topalidou, Cattin-Ortolá, et al., 2018). Thus, EARP and EIPR1 may be important for acting near the TGN to ensure the retention of processed cargos in mature DCVs.

Here we investigate the role of CCDC186 in DCV biogenesis, maturation, and cargo sorting in 832/13 cells. Our results suggest that CCDC186 acts together with EARP to control several aspects of DCV cargo sorting, including the post-Golgi removal of some non-DCV cargos and retention of mature DCV cargos.

## RESULTS

### CCDC186 is required for normal insulin secretion and distribution of mature DCV cargos

Our studies in *C. elegans* show that *cccp-1* (the *C. elegans* homolog of *Ccdc186*/C10orf118) and *eipr-1* act in the same genetic pathway to control locomotion and neuronal DCV cargo levels (Figure S1). We recently found that EIPR1 is needed for proper insulin secretion and distribution of mature DCV cargos in rat insulinoma 832/13 cells (Topalidou, Cattin-Ortolá, et al., 2018). CCDC186 is expressed in endocrine cell lines such as 832/13 and in multiple tissues in mice, including brain, heart, liver, kidney, and spleen (Figure S2A). To investigate the function of CCDC186 in 832/13 cells, we generated a *Ccdc186* knock-out (KO) cell line and a KO line rescued by transgenic expression of a wild-type *Ccdc186* cDNA (Figure S2B-D).

We found that *Ccdc186* KO cells share many phenotypes with *Eipr1* KO cells. In *Ccdc186* KO cells, stimulated insulin secretion was significantly reduced (Figure 1A), but the total cellular insulin level was not changed (Figure 1B). Both proinsulin secretion and the cellular level of proinsulin were slightly increased in *Ccdc186* KO cells, but the effects were not statistically significant (Figure S3A,B). Additionally, the ratio of total cellular proinsulin to insulin was unchanged in the *Ccdc186* KO cells (Figure S3C) and there was no change in constitutive secretion (Figure S3D). These results suggest that CCDC186, like EIPR1, is required for normal levels of regulated secretion but is not required for the processing of proinsulin to insulin.

**Figure 1.**
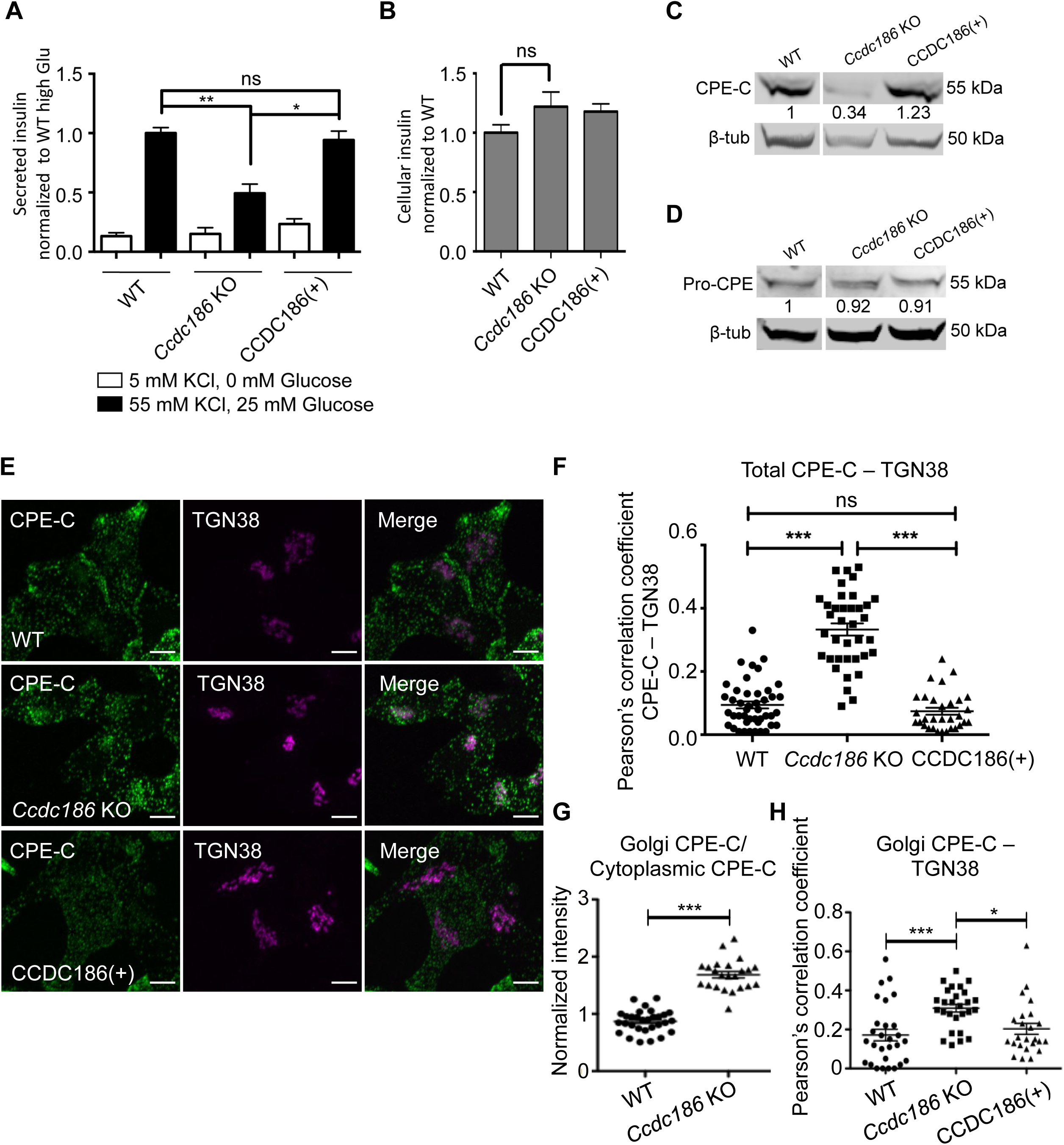
*Ccdc186* knock-out cells have defective insulin secretion and distribution of mature DCV cargos. A. Insulin secretion from WT, *Ccdc186* KO, and *Ccdc186*(+) rescued 832/13 cells under resting conditions (no glucose, 5 mM KCl, white bars) and stimulating conditions (25 mM glucose, 55 mM KCl, black bars). All values were normalized to the WT value in stimulating conditions (WT high Glu). n = 8; *p<0.05, **p<0.01, ns p>0.05, error bars = SEM. B. Total insulin content in WT, *Ccdc186* KO, and *Ccdc186*(+) rescued 832/13 cells. All values were normalized to WT. n = 8; ns p>0.05, error bars = SEM. We performed three biological replicates. For each replicate, the same cells were used to determine the amount of insulin secreted under resting conditions, stimulating conditions, and the amount of total cellular insulin. C. *Ccdc186* KO cells have reduced levels of the processed form of CPE. Detergent lysates from WT, *Ccdc186* KO, and *Ccdc186*(+) rescued 832/13 cells were blotted with an antibody to the C-terminus of CPE (CPE-C). β-tubulin was used as a loading control. The experiment was repeated three times. D. The levels of the unprocessed form of CPE (pro-CPE) are not affected by the loss of CCDC186. Detergent lysates from WT, *Ccdc186* KO, and *Ccdc186*(+) rescued 832/13 cells were blotted with an antibody to pro-CPE. β-tubulin was used as a loading control. The experiment was repeated three times with similar results. The data shown for the WT are the same shown in Figure 3B of (Topalidou, Cattin-Ortolá, et al., 2019, manuscript submitted to *MBoC*) since these experiments were run in parallel with the same WT control. E. The mature processed form of CPE is localized at or near the TGN in *Ccdc186* KO cells. Representative confocal images of WT, *Ccdc186* KO, and *Ccdc186*(+) rescued 832/13 cells costained with the CPE C-terminal antibody (CPE-C) and TGN38. Maximum-intensity projections. Scale bars: 5 µm. The experiment was repeated three times with similar results. F. Quantification of the colocalization between the mature form of CPE and the TGN marker TGN38. Maximum-intensity projection images were obtained and Pearson’s correlation coefficients were determined by drawing a line around each cell. The data shown are from one experiment. n=44 for WT, n=37 for *Ccdc186* KO, n=31 for *Ccdc186*(+); ***p<0.001, ns p>0.05; error bars = SEM. G. *Ccdc186* KO cells have an increased proportion of CPE localized near the TGN. Fluorescence of a region of interest around the TGN divided by the fluorescence of a region of the same size in the cytoplasm. The data shown are from one experiment. n=30 for WT and n=23 for *Ccdc186* KO, ***p<0.001, error bars = SEM. The data shown for the WT are the same shown in Figure 3G of (Topalidou, Cattin-Ortolá, et al., 2019, manuscript submitted to *MBoC*) since these experiments were run in parallel with the same WT control. H. *Ccdc186* KO cells have an increased amount of CPE localized at or near the TGN. Maximum-intensity projection images of z-stacks encompassing the TGN38 signal were obtained and Pearson’s correlation coefficients were determined by drawing a line around the TGN38 positive signal. The data shown are from one experiment. n=28 for WT, n=27 for *Ccdc186* KO, n=24 for *Ccdc186*(+); *p<0.05, ***p<0.001; error bars = SEM.

By measuring exocytotic events using NPY tagged with the pH-sensitive GFP pHluorin, we found that *Eipr1* KO cells have an apparent exocytosis defect that could contribute to their insulin secretion defect (Topalidou, Cattin-Ortolá, et al., 2019, manuscript submitted to *MBoC*). Similarly, we found that exocytotic events were strongly reduced in *Ccdc186* KO cells under stimulating conditions (Figure S4), suggesting that *Ccdc186* cells also have an exocytosis defect. However, a caveat to this experiment is that this assay may have underestimated the true exocytosis rate if it was not possible to detect DCVs carrying reduced amounts of NPY::pHluorin. As *Ccdc186* KO cells appear to carry reduced amounts of cargos in mature DCVs (see below), it is possible that DCV exocytotic events in this mutant line were more difficult to detect.

The processing of proinsulin to insulin is mediated by several processing enzymes, including the proprotein convertases PC1/3 (PCSK1) and carboxypeptidase E (CPE) (Naggert et al., 1995; Smeekens et al., 1992). We recently found that EIPR1 is needed for the normal cellular levels of mature PC1/3 and CPE in 832/13 cells (Topalidou, Cattin-Ortolá, et al., 2019, manuscript submitted to *MBoC*). Similarly, we found that loss of CCDC186 resulted in reduced levels of mature PC1/3 and CPE (Figure 1C and Figure S5A,B). By contrast, the level of the unprocessed pro form of CPE (pro-CPE) was not altered in *Ccdc186* KO cells (Figure 1D), similar to *Eipr1* KO cells (Topalidou, Cattin-Ortolá, et al., 2019, manuscript submitted to *MBoC*). These results suggest that CCDC186, like EIPR1, acts after the processing of pro-CPE.

We found that EIPR1 is required for the proper cellular distribution of mature CPE (CPE-C) and insulin, but not of pro-CPE and proinsulin (Topalidou, Cattin-Ortolá, et al., 2019, manuscript submitted to *MBoC*). In wild-type cells, insulin and mature CPE are found as colocalized puncta spread throughout the cytoplasm (Figure S5C). By contrast, in *Ccdc186* KO cells, mature CPE and insulin are localized primarily to a perinuclear region that partially colocalizes with TGN38 (Figure 1E,F and Figure S6A,B). The redistribution of CPE and insulin in *Ccdc186* KO cells appears to be due to decreased levels of mature CPE and insulin in vesicles in the cell periphery and increased retention of CPE and insulin in or near the TGN (Figure 1G,H and Figure S6C,D). In contrast to mature CPE and insulin, pro-CPE and proinsulin are localized near the TGN in WT and *Ccdc186* KO cells (Figure S7). All of the CPE and insulin localization results of *Ccdc186* KO cells are similar to those of *Eipr1* KO cells (Topalidou, Cattin-Ortolá, et al., 2019, manuscript submitted to *MBoC*). These results suggest that CCDC186, like EIPR1, acts at a post-Golgi level and that it is needed for the proper levels and cellular distribution of several mature DCV cargos, including insulin and the insulin processing enzymes PC1/3 and CPE.

One possible explanation for the increased colocalization of insulin and mature CPE with TGN38 is that there might be increased acidification of the Golgi in the *Ccdc186* KO cells that allows proteolytic processing of DCV cargos. However, there was little change in the pH of the Golgi in *Ccdc186* KO cells and if anything a slight increase (Figure S8). Additionally, pulse-chase experiments show that DCV cargo is not stuck in the Golgi in *Ccdc186* KO cells, but does exit the TGN and move out to the periphery (Figure S9). These results are consistent with CCDC186 acting in a post-Golgi step in the DCV biogenesis pathway.

### CCDC186 localizes to a ring-like compartment around proinsulin

We previously reported that EARP localizes to two distinct compartments: an endosomal recycling compartment and a CCDC186-positive compartment relevant to DCV biogenesis (Topalidou et al., 2018). In *C. elegans* neurons and 832/13 cells, CCCP-1 localizes at or close to the TGN and immature DCVs (Ailion et al., 2014; Cattin-Ortolá et al., 2017). These data were based largely on overexpressed GFP-tagged constructs. To more specifically define the localization of endogenous CCDC186 in 832/13 cells, we performed costaining of CCDC186 with a number of compartmental markers. Endogenous CCDC186 is localized to an area immediately adjacent to but not tightly overlapping with the TGN marker TGN38 (Figure 2A and Figure S10). By contrast, CCDC186 colocalized most closely with the SNARE protein syntaxin 6, and the iDCV cargos chromogranin A (CgA) and proinsulin (Figure 2A and Figure S10). Syntaxin 6 and proinsulin are localized to the TGN and iDCVs, but are excluded from mature DCVs (Klumperman et al., 1998; Wendler et al., 2001). CgA is localized mainly to a perinuclear area, but also to some puncta throughout the cytoplasm and the cell periphery (Figure S10), suggesting that CgA is present at the TGN, iDCVs, and some mature DCVs. CCDC186 does not colocalize tightly with markers of the ERGIC (ERGIC53), cis-Golgi (GM130), early endosome (EEA1), or lysosome (LAMP1) (Figure 2A and Figure S10). We conclude that CCDC186 is localized at or close to the TGN and iDCVs.

**Figure 2.**
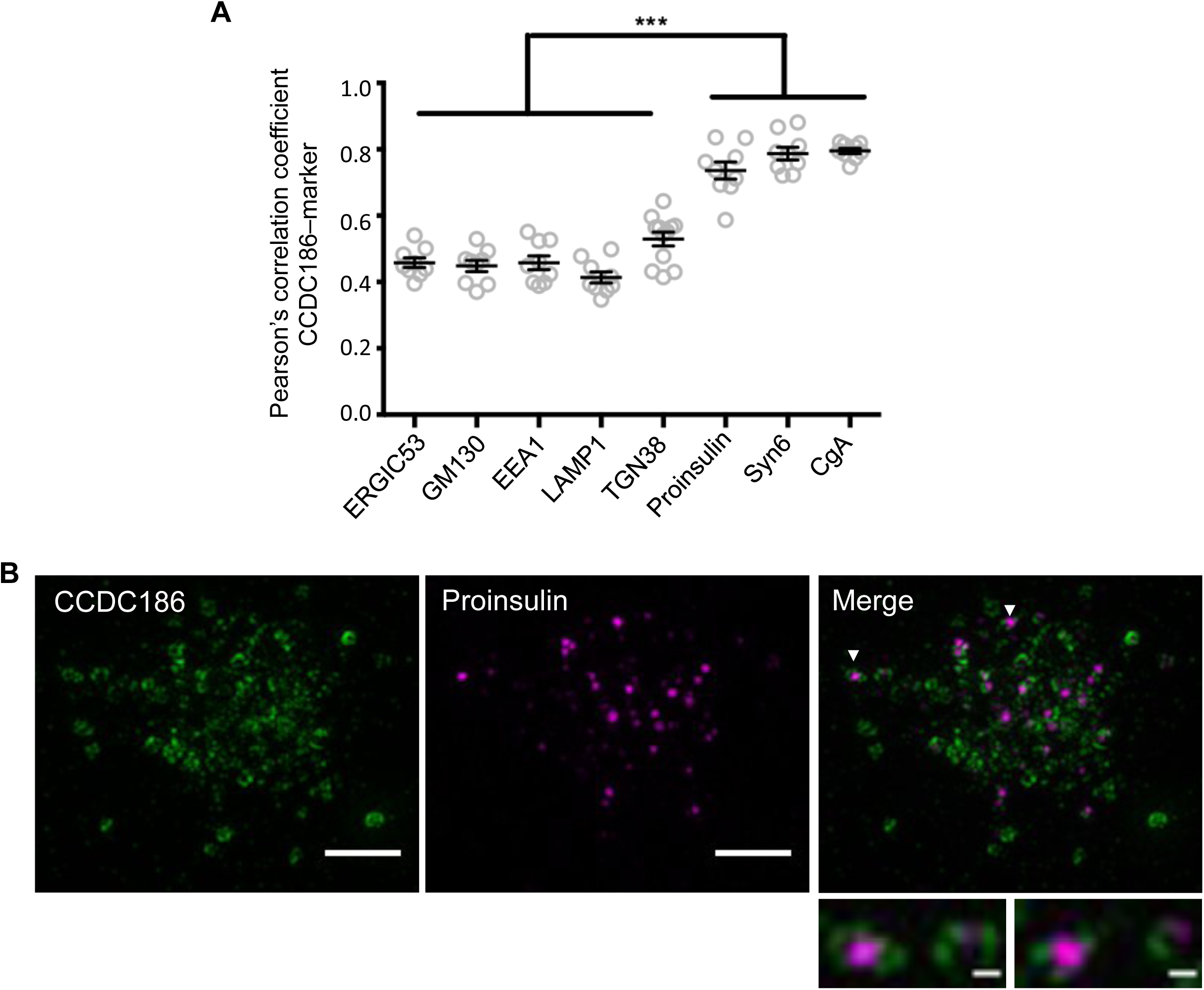
Endogenous CCDC186 localizes to ring-like structures around proinsulin. A. Pearson’s correlation coefficient was measured using confocal images (Figure S10) to quantify the colocalization between endogenous CCDC186 and markers of the different cell compartments (ERGIC53 for the ERGIC; GM130 for the cis-Golgi; EEA1 for early-endosomes; LAMP1 for lysosomes; TGN38 for the TGN; and Syntaxin 6 (Syn6), chromogranin A (CgA) and proinsulin for the TGN and immature DCVs). n=12 cells for TGN38 and n=9 cells for every other marker, ***p<0.001, error bars = SEM. B. Representative structured illumination microscopy (SIM) images of 832/13 cells costained for endogenous CCDC186 and proinsulin. Maximum-intensity projections. Upper panels, scale bars: 2 µm. Lower panels, scale bars: 200 nm.

To determine the localization of CCDC186 at higher resolution, we used structured illumination microscopy (SIM, Figure 2B) and stimulated emission depletion microscopy (STED, Figure S11). With both techniques, we found that endogenous CCDC186 is localized to ring-like structures of variable size (∼ 200-500 nm diameter) rather than the puncta seen by conventional confocal microscopy. Interestingly, the rings of CCDC186 often surrounded proinsulin, suggesting that CCDC186 localizes at or close to iDCV membranes, or to a subdomain of the TGN carrying DCV cargos.

### The CCDC186 C-terminal domain is in close proximity to carboxypeptidase D

To further characterize the compartment where CCDC186 localizes, we used the proximity biotinylation BioID approach to investigate the protein composition of CCDC186-positive membranes (Roux et al., 2012). To increase the specificity of labeling the membranes bound by CCDC186, we used a mitochondrial relocation strategy because it should give a background composed of mitochondrial proteins that we could easily discard (Shin, Gillingham, et al., 2017).

CCDC186 was ectopically expressed and localized to mitochondria through attachment to the mitochondrially-localized transmembrane domain of monoamine oxidase (MAO) (Wong and Munro, 2014) (Figure S12A). We generated constructs with full-length CCDC186 and the CCDC186 C-terminal domain CC3, each fused at their C-terminus to an HA tag and to MAO (Figure S12B). CC3 binds RAB2 and membranes and is both necessary and sufficient for localization of CCDC186 to the compartment near the TGN (Cattin-Ortolá et al., 2017). Experiments using the full-length CCDC186 protein fused to MAO resulted in low expression of the construct and high toxicity in 832/13 cells, so we did not pursue further. We showed by immunofluorescence that CC3::HA::MAO colocalized well with the mitochondrial marker MitoTracker but not with the TGN marker TGN38 (Figure S12C). We also observed that CC3::HA::MAO colocalized with constitutively active forms of both RAB2A and RAB2B (Figure S12D,E). Thus, the mitochondrial relocation experiment recapitulates the known interaction of the CC3 domain with RAB2.

For BioID, we fused the promiscuous biotin ligase BirA* to the C terminus of CC3 (CC3:: BirA*::HA::MAO) (Figure 3A,B). BirA*::HA::MAO was used as a negative control. We generated PC12 cell lines that stably expressed these constructs instead of 832/13 cells for technical reasons (see Material and Methods) and performed mass spectrometry on the biotinylated material pulled down by streptavidin beads. Many peptides for CCDC186 were found in the CC3 sample but few were found in the negative control, demonstrating the efficiency of the labeling (Figure 3C and Table S1). Comparison of CC3::HA::MAO with the negative control also gave carboxypeptidase D (CPD) as a possible specific hit (Figure 3C, Figure S13, and Table S1). Immunoblotting of streptavidin-purified biotinylated proteins from cells expressing the different BirA* constructs using an anti-CPD antibody validated the mass spectrometry result (Figure 3D). We also verified that expression of CC3::HA::MAO resulted in partial relocation of CPD to the mitochondria (Figure 3E). CPD belongs to the family of metallocarboxypeptidases that remove arginine or lysine residues from the C-terminus of peptides, but its targets and cellular function are unclear (Fricker, 2013). CPD localizes mainly to the TGN but also traffics to the plasma membrane and endosomes (Varlamov and Fricker, 1998), and CPD also localizes to iDCVs (Varlamov et al., 1999). Because CPD is a transmembrane protein (Kuroki et al., 1995; Xin, Varlamov, et al., 1997), CCDC186 might interact with a membrane carrying CPD, such as the TGN or vesicles that bud from the TGN.

**Figure 3.**
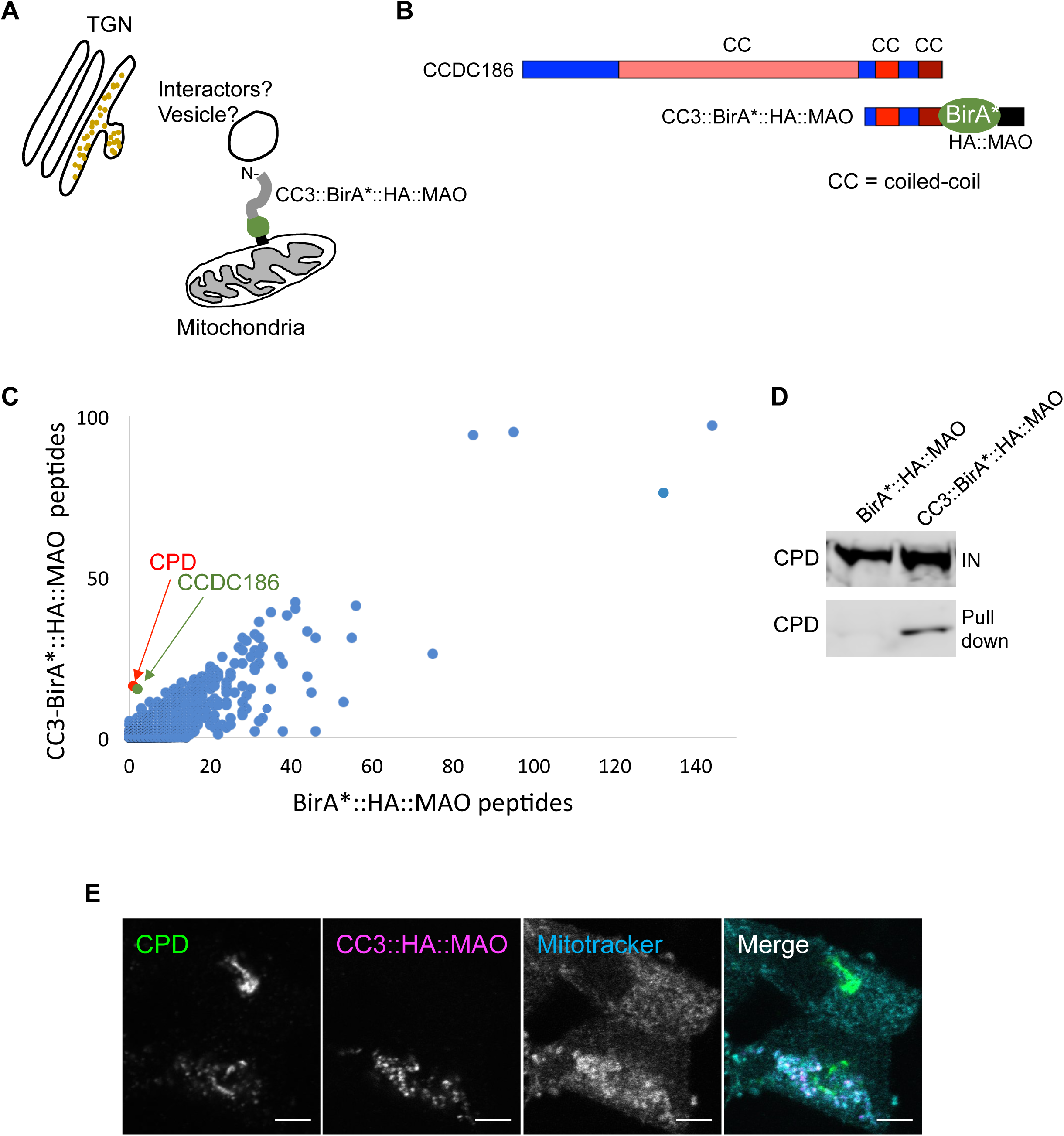
The CCDC186 C-terminal domain CC3 is in close proximity to carboxypeptidase D (CPD). A. Schematic of the BioID experiment. The promiscuous biotin ligase BirA* biotinylates proteins that are in close proximity (approximately < 10 nm). B. Domain structure of CCDC186. The CC3 fragment was fused to the mitochondria-targeting domain MAO (in black) and to the promiscuous biotin ligase BirA* (in green). The predicted coiled-coil domains are marked with different shades of red. CC=coiled-coil. C. Mass-spectrometric (MS) analysis of biotinylated proteins from PC12 cells stably expressing CC3::BirA*::HA::MAO or the negative control BirA*::HA::MAO. The plot compares the number of unique peptides between CC3::BirA*::HA::MAO and the negative control. The data shown are from one out of three independent experiments. D. CPD is biotinylated by CC3::BirA*::HA::MAO. Biotinylated proteins from PC12 cell extracts stably expressing BirA*::HA::MAO or CC3::BirA*::HA::MAO were purified with streptavidin-coated beads and immunoblotted for CPD. IN: input. CPD was found enriched in all three independent pulldowns by MS, but it could only be detected by Western blotting in the sample that showed the highest enrichment in MS. E. CPD is partially relocated to the mitochondria by CC3::HA::MAO. Representative confocal images of 832/13 cells transiently transfected with CC3::HA::MAO, incubated with MitoTracker, and costained for HA and CPD. Single slices. Scale bars: 5 µm.

### CCDC186 is required for the removal of carboxypeptidase D from dense-core vesicles

Studies in AtT-20 cells have shown that although CPD localizes to iDCVs, it is mostly absent from mature DCVs, indicating that CPD is removed from iDCVs during maturation (Varlamov et al., 1999). We found that CCDC186 and CPD partially colocalized in a perinuclear area (Figure 4A). Importantly, although CPD colocalized with the TGN marker TGN38 in WT cells (Figure 4B), CPD was largely mislocalized to puncta distributed throughout the cytoplasm in *Ccdc186* KO cells and this phenotype was rescued by expression of *Ccdc186*(+) (Figure 4B). Because EIPR1 and CCDC186 function in the same pathway in DCV biogenesis (Figure S1), we tested whether CPD localization also depends on EIPR1. We found that CPD was localized to puncta distributed throughout the cytoplasm in *Eipr1* KO cells, but to a lesser extent than in the *Ccdc186* KO cells (Figure 4B). Thus, CCDC186 and EIPR1 are required to restrict CPD localization to the TGN.

**Figure 4.**
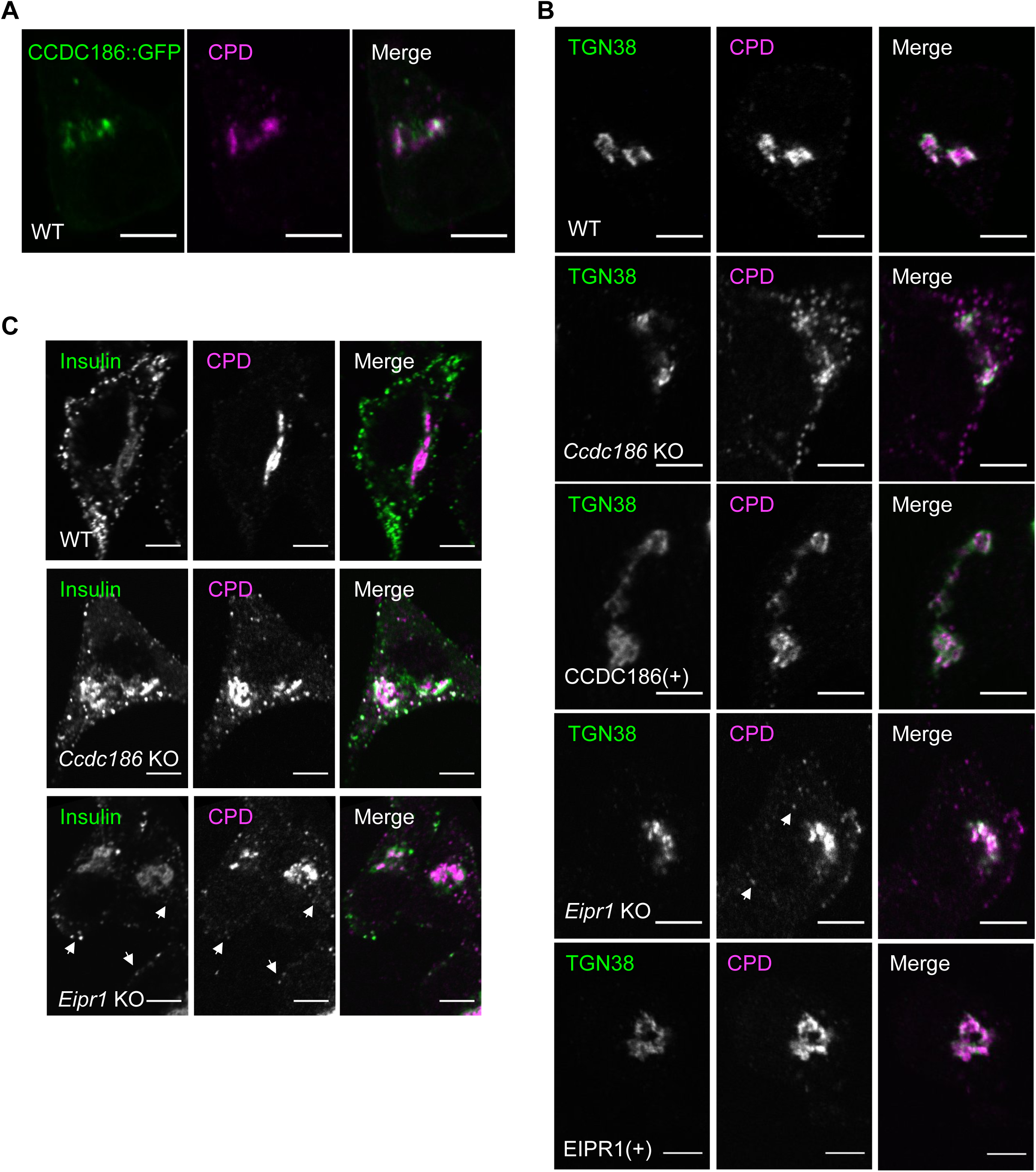
CPD is mislocalized to mature DCVs in *Ccdc186* KO and *Eipr1* KO 832/13 cells. A. CPD partially colocalizes with CCDC186::GFP. Representative confocal images of 832/13 cells transiently transfected with CCDC186::GFP and costained for endogenous CPD and GFP. Single slices. Scale bars: 5 µm. B. CCDC186 and EIPR1 are required to restrict CPD localization to the TGN. Representative confocal images of WT, *Ccdc186* KO, *Ccdc186*(+) rescued, *Eipr1* KO and *Eipr1*(+) rescued 832/13 cells costained for endogenous CPD and the TGN marker TGN38. The white arrows indicate puncta in *Eipr1* KO cells. Single slices. Scale bars: 5 µm. The experiment was repeated three times with similar results. C. In the absence of CCDC186 or EIPR1, CPD is mislocalized to insulin-positive vesicles. Representative confocal images of WT, *Ccdc186* KO, and *Eipr1* KO 832/13 cells costained for endogenous CPD and insulin. The intensity of CPD signal in WT cells was increased to show the occasional presence of low intensity CPD-positive puncta that do not overlap with insulin. The white arrows indicate CPD-positive puncta that colocalize with insulin in *Eipr1* KO cells. Single slices. Scale bars: 5 µm. The experiment was repeated three times.

We reasoned that the punctate cytoplasmic pattern of CPD in *Ccdc186* KO cells and *Eipr1* KO cells could be the result of defective CPD removal from iDCVs. Consistent with this possibility, most CPD-positive cytoplasmic puncta colocalized with insulin in *Ccdc186* KO cells and *Eipr1* KO cells (78 +/- 10 % of CPD cytoplasmic puncta in *Ccdc186* KO cells are insulin positive), suggesting that these CPD-positive puncta represent mature DCVs (Figure 4C).

To further test whether CPD is mislocalized to mature DCVs in the *Ccdc186* and *Eipr1* mutants, we performed cellular fractionation experiments by equilibrium sedimentation through a sucrose gradient to separate the Golgi and DCV compartments (Figure 5). The absence of CCDC186 or EIPR1 did not have a significant effect on the sedimentation profile of PC1/3, indicating that the density of DCVs is not significantly altered in either knock-out cell line. However, we found that some of the CPD shifted to a heavier fraction that comigrated with the DCV marker PC1/3 in both *Ccdc186* and *Eipr1* KO cells (Figure 5), consistent with the model that CPD is retained in mature DCVs in these mutants. Together, these immunofluorescence and biochemical data indicate that CCDC186 and EIPR1 are required for the removal of CPD from iDCVs.

**Figure 5.**
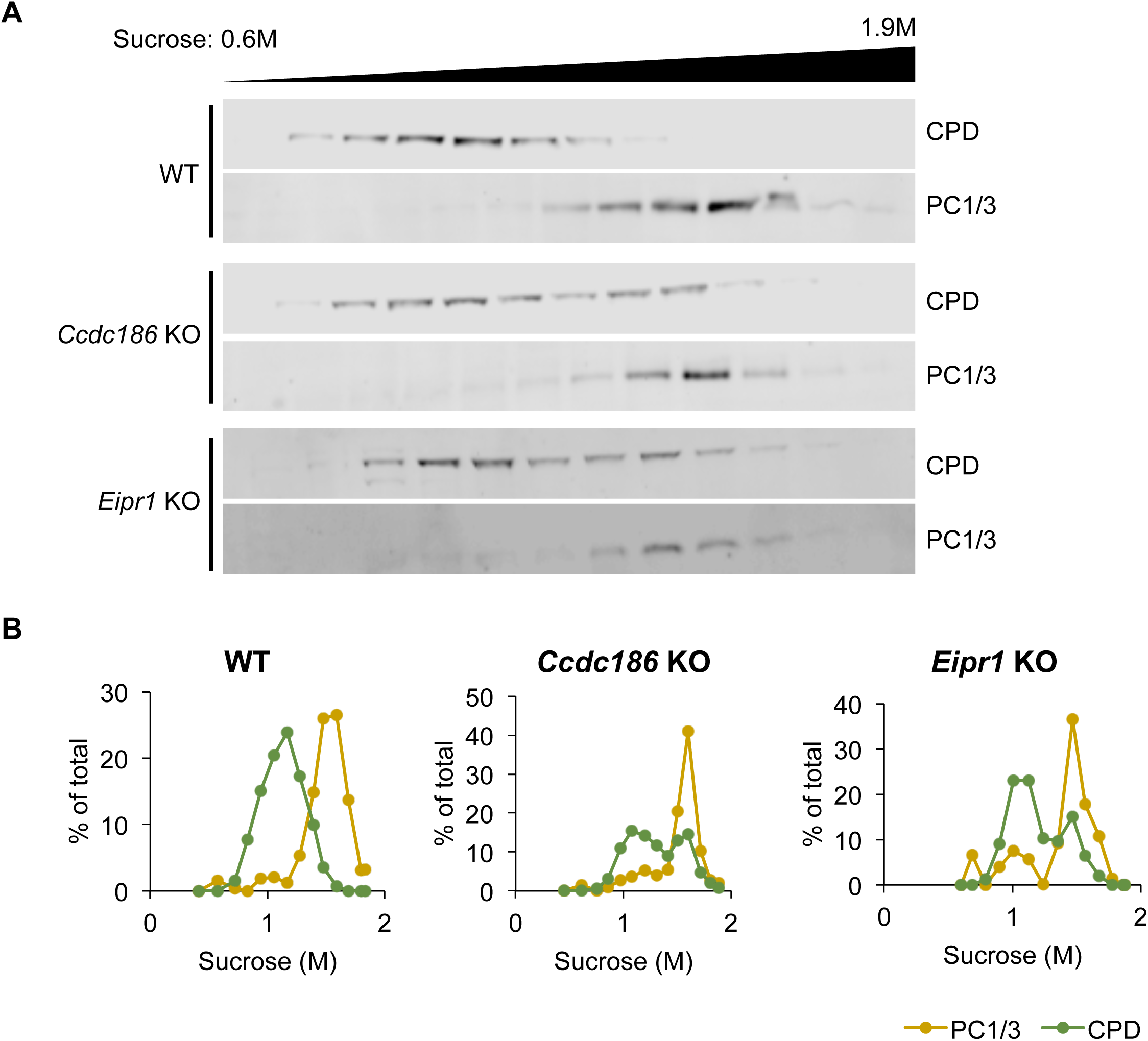
CPD cosediments with mature DCV cargos in *Ccdc186* KO and *Eipr1* KO cells. A. Post-nuclear supernatants from WT, *Ccdc186* KO, and *Eipr1* KO cells were separated by equilibrium sedimentation through 0.6–1.9 M sucrose. Fractions were collected from the top of the gradient and blotted with antibodies against PC1/3 (mature DCV marker) and CPD. The intensity and contrast of each blot was adjusted to show similar band intensities between cell types. The data shown are from one representative experiment of three independent experiments with similar results. B. Band intensity for each fraction was quantified using FIJI, presented as a percentage of total band intensity, and plotted against the sucrose concentration of the fraction. The data shown are from one representative experiment of three independent experiments with similar results.

Similar to CPD, other proteins such as the cation-dependent mannose 6-phosphate receptor (CD-MPR) are removed from mature DCVs (Klumperman et al., 1998). Interestingly, although the CD-MPR is a transmembrane protein with similar cytosolic sorting motifs as CPD (Bonifacino and Traub, 2003; Eng et al., 1999), its removal from iDCVs is not dependent on CCDC186 or EIPR1 (Figure S14). Thus, different mechanisms are needed for the removal of CPD and the CD-MPR from immature DCVs.

## DISCUSSION

In this study we investigated the role of CCDC186 in DCV function in the rat pancreatic beta-cell line 832/13. Using CRISPR knock-outs and rescuing experiments, we found that CCDC186 acts in the post-Golgi maturation of DCVs to control the retention of mature cargos such as insulin and CPE, and the removal of cargos such as CPD. We propose that CCDC186 acts as a regulator of DCV maturation by ensuring that mature DCVs carry the proper type and amount of cargo.

### CCDC186 controls retention of cargos in mature DCVs

*C. elegans* mutants in *rab-2, cccp-1* (the *C. elegans* homolog of *Ccdc186*), *eipr-1,* and the EARP complex subunits have reduced levels of DCV cargo (Ailion et al., 2014; Edwards et al., 2009; Paquin et al., 2016; Sumakovic et al., 2009; Topalidou et al., 2016). This conclusion was based mainly on assays using exogenous DCV cargos overexpressed in neurons. In this study, we used the rat insulin-secreting insulinoma cell line 832/13 to investigate the secretion, localization, and processing of endogenous DCV cargos. We found that cells lacking CCDC186 remain responsive to stimulated secretion, but secrete reduced levels of insulin and have reduced levels of insulin and other luminal cargos in mature DCVs at the cell periphery. Our data are most consistent with a model in which CCDC186 acts in a post-Golgi step to control retention of DCV cargos (Figure 6), rather than acting in the TGN to control sorting of cargos by entry or having a direct effect on peptide processing. First, *Ccdc186* KO cells show no defect in the levels or localization of immature DCV cargos such as proinsulin and pro-CPE, but have reduced levels of mature cargos such as CPE. Second, the exogenous DCV cargo ANF-GFP was not stuck in the TGN in *Ccdc186* KO cells, but moved to the periphery following release of a low-temperature block. Third, there is not increased acidity of the TGN in *Ccdc186* KO cells. Fourth, there is no apparent defect in processing of proinsulin to insulin in *Ccdc186* KO cells. These data suggest that early steps in the sorting and processing of pro-proteins occur normally in the absence of CCDC186, but that mature DCV cargos are lost in a post-Golgi step. *Eipr1* KO cells have similar defects (Topalidou, Cattin-Ortolá, et al., 2018, and this study), supporting the genetic data indicating that CCDC186 and EIPR1/EARP act in the same pathway to regulate DCV maturation.

**Figure 6.**
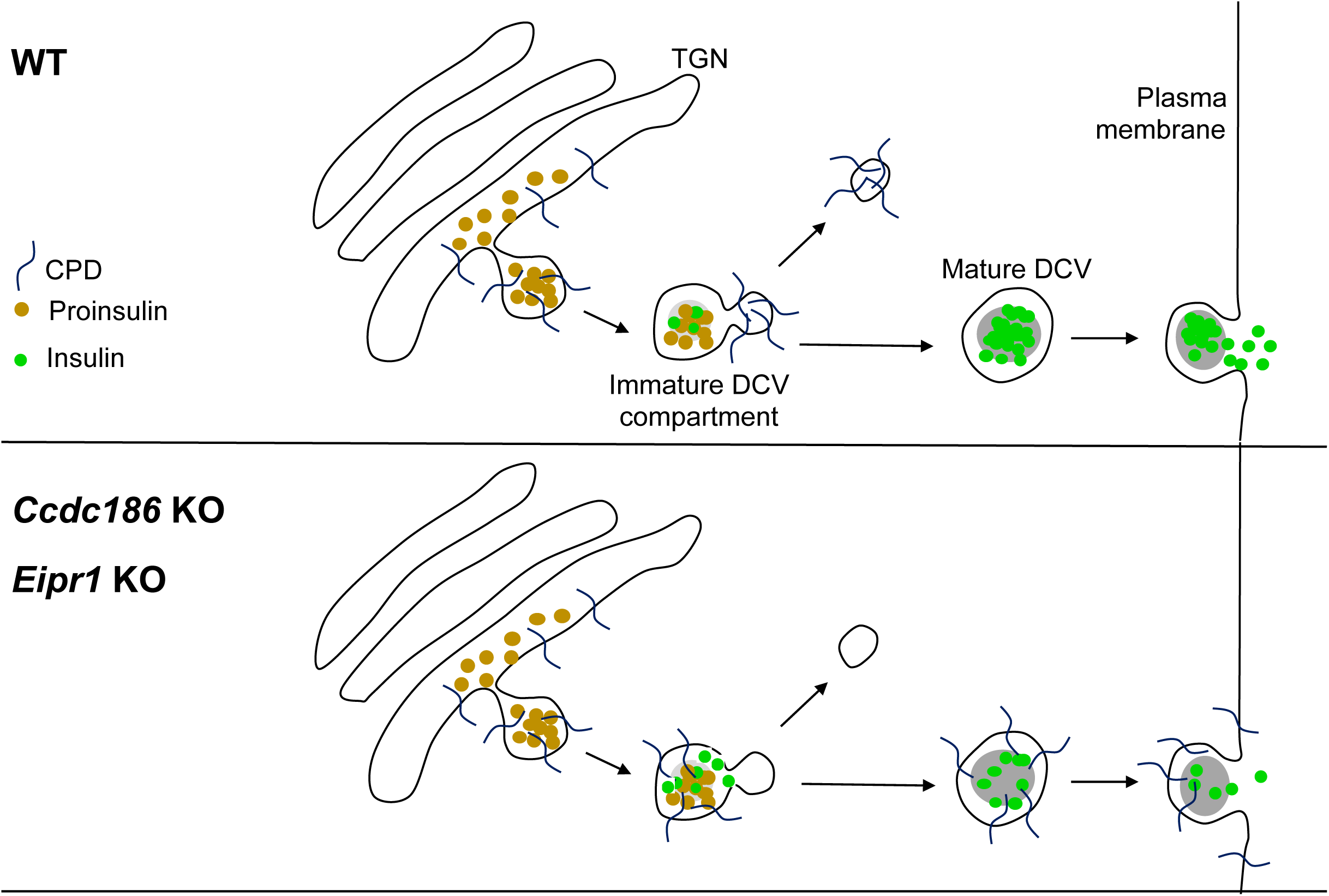
Model for CCDC186 and EIPR1 function in DCV maturation. In *Ccdc186* KO cells, insulin and other luminal DCV cargos are reduced in mature DCVs in the cell periphery and mildly retained near the TGN. Proinsulin and other unprocessed cargos are not affected. Thus, CCDC186 is required for the retention of mature DCV cargos. Additionally, CPD is retained in mature DCVs in *Ccdc186* KO cells, suggesting that CCDC186 is also required for the removal of unwanted cargo from immature DCVs. *Eipr1* KO cells have similar phenotypes to *Ccdc186* KO cells, suggesting that CCDC186 and EIPR1/EARP act together to control the post-Golgi retention and removal of DCV cargos.

### CCDC186 controls the removal of CPD from immature DCVs

In a proximity biotinylation screen, we identified CPD as being in close proximity to the C-terminal CC3 domain of CCDC186. CPD is a transmembrane protein with a short cytosolic tail (∼ 60 amino acids) and a longer N-terminal intraluminal domain (Kuroki et al., 1995; Xin, Varlamov, et al., 1997). CPD is an ubiquitously expressed member of the metallocarboxypeptidase family, but its role in the processing of peptides destined to the regulated secretory pathway remains unclear (Fricker, 2013).

The basis of the interaction between CCDC186 and CPD-positive membranes is not clear. In coimmunoprecipitation experiments from 832/13 detergent lysates, we did not detect an interaction between GFP-tagged CCDC186 and either endogenous CPD or overexpressed mCherry-tagged CPD cytosolic tail (data not shown), indicating that CCDC186 and CPD do not interact directly or their interaction is too weak to detect by this method. CCDC186 might bind to CPD-positive membranes via interaction with another protein or by directly binding to membranes. The CC3 domain of CCDC186 is capable of direct association with synthetic membrane liposomes and is both necessary and sufficient for CCDC186 localization to TGN/iDCV membranes (Cattin-Ortolá et al., 2017).

Studies in AtT-20 cells have shown that CPD localizes to the TGN and to iDCVs but is mostly absent from mature DCVs (Varlamov et al., 1999). We show here that in the absence of CCDC186 or EIPR1, CPD remains in mature, insulin-positive DCVs. Thus, we propose that CCDC186 and EIPR1 ensure the removal of CPD from DCVs during maturation. Similar to CPD, other proteins such as the cation-independent and the cation-dependent mannose 6-phosphate receptors (CI-MPR and CD-MPR) are removed from iDCVs by a pathway that depends on AP-1 and clathrin (Dittie et al., 1999; Klumperman et al., 1998). The fate of these proteins once removed is unknown, but it has been proposed that they follow the endosomal route (Arvan and Halban, 2004; Feng and Arvan, 2003). Strikingly, although the CD-MPR is also a transmembrane protein with similar cytosolic sorting motifs as CPD (Bonifacino and Traub, 2003; Eng et al., 1999), its removal from iDCVs was not affected by the absence of CCDC186 or EIPR1, suggesting that different cargos may be removed from immature DCVs by different mechanisms. Our data suggest that CCDC186 and EIPR1 ensure the removal of CPD from DCVs (Figure 6).

### What is the role of CCDC186 in DCV biogenesis?

Our data suggest that CCDC186 is important for both the retention of mature DCV cargos and the removal of some non-DCV cargos from immature DCVs. These are both post-Golgi processes, but it is unclear whether they are independent or controlled coordinately by a single action of CCDC186. Here we propose that CCDC186 serves a role in ensuring that the proper amount and type of cargo remains in mature DCVs.

Our data suggest that CCDC186 acts after the processing of peptide cargos. This distinguishes CCDC186 from other proteins important for DCV biogenesis such as PICK1, ICA69, and HID1. Like CCDC186, these proteins are also required for normal levels of mature DCV cargos (Cao, Mao, et al., 2013; Du, Zhou, Zhao, Cheng, et al., 2016; Holst, Madsen, et al., 2013; Hummer et al., 2017), but in contrast to CCDC186, loss of PICK1, ICA69, or HID1 leads to a defect in the processing of proinsulin to insulin (Cao, Mao, et al., 2013; Du, Zhou, Zhao, Cheng, et al., 2016). PICK1, ICA69, and HID1 have been proposed to act at an early step of DCV biogenesis during budding of immature DCVs from the TGN; our data suggest that CCDC186 and EIPR1 act at a post-Golgi step during DCV maturation.

The connection between CCDC186 and the EARP complex suggests there may be trafficking between EARP-positive endosomes and CCDC186-positive TGN/iDCV membranes. Such a trafficking step may be involved in the removal of CPD from iDCVs. Trafficking from endosomes to iDCVs may also be involved in the retrieval of DCV membrane proteins following exocytosis. It has been shown that after DCV exocytosis, transmembrane proteins such as phogrin and PAM are recycled and traffic through endosomes before reaching newly-generated DCVs (Bäck et al., 2010; Vo et al., 2004). CCDC186 and EARP could mediate the recycling of DCV proteins from endosomes to iDCVs; disrupting this recycling pathway could also indirectly impair the proper trafficking of sorting receptors required for retention of luminal DCV cargos. Similar to loss of CCDC186 or EIPR1, loss of other proteins such as BAIAP3 and Vti1a/Vti1b that are involved in retrograde trafficking from endosomes to Golgi also leads to reduced levels of DCV cargos (Emperador-Melero et al., 2018; Zhang et al., 2017). However, BAIAP3 and Vti1a/b appear to affect endosome to Golgi trafficking more generally because their loss also affects non-DCV cargos and perturbs Golgi morphology (Emperador-Melero et al., 2018; Zhang et al., 2017), whereas loss of CCDC186 or EIPR1 does not affect non-DCV cargos and has no apparent effect on Golgi morphology as assayed by TGN38 staining. CCDC186 and EARP may be specifically involved in endosomal trafficking of DCV-related factors or cargos.

The idea that CCDC186 and EARP affect DCV cargo levels by regulating trafficking to or from endosomes is supported by experiments in *C. elegans* that found that disrupting trafficking to the endolysosomal system could restore levels of DCV cargo in *rab-2* mutants, suggesting that cargo is lost via an endolysosomal route (Edwards et al., 2009; Sumakovic et al., 2009). Given that CCDC186/CCCP-1 and EARP act in the same genetic pathway as RAB-2 (Ailion et al., 2014; Topalidou et al., 2016), it is likely that they also act to retain cargos in mature DCVs by preventing the loss of cargos to the endolysosomal system. Interestingly, a recent study showed that cholesterol-deficient DCVs are lost through lysosomal degradation, leading to a reduction in insulin secretion (Hussain, Harris, et al., 2018). This observation raises the intriguing possibility that CCDC186 and EARP may be involved not only in protein trafficking, but also in cholesterol transfer to DCVs. Supporting this possibility, the EARP complex associates with Rab11-positive recycling endosomes (Schindler, Chen, et al., 2015), and Rab11 and its effector RELCH act to promote cholesterol transfer from recycling endosomes to the TGN (Sobajima et al., 2018). Perhaps CCDC186 and EARP are also involved in this process.

## MATERIALS & METHODS

### *C. elegans* strains, molecular biology, and plasmids

*C. elegans* strains were cultured using standard methods (Brenner, 1974). A complete list of strains used is provided in the Strain List (Table S2). A complete list of constructs is provided in the plasmid list (Table S3). Vector backbones and PCR-amplified fragments containing 20-30 bp overlapping ends were combined by Gibson cloning (Gibson et al., 2009).

### Locomotion assays

To measure *C. elegans* locomotion, first-day adults were picked to 2 to 3-day-old lawns of OP50 bacteria, stimulated by touching their tail using a worm pick, and body bends were then counted for one minute. A body bend was defined as the movement of the worm from maximum to minimum amplitude of the sine wave.

### *C. elegans* imaging and image analysis

Young adult worms were placed in 8 µl of 50 mM sodium azide on 2% agarose pads and anesthetized for 10 min. Images of animals with dorsal side up were obtained using a Nikon Eclipse 80i wide-field compound microscope. The same region of the dorsal cord around the vulva was imaged in all worms. All strains in an experiment were imaged on the same day and the microscope settings were kept constant. Maximum-intensity projections were quantified using ImageJ. For each animal, the total fluorescence in five regions of interest including the dorsal cord was measured and the background fluorescence of a region of the same size adjacent to the dorsal cord was subtracted from each. The final fluorescence intensity for each animal was calculated as the average total fluorescence of the five regions.

### Cell culture

The 832/13 cell line is an INS-1-derived clone that was isolated by Dr. Christopher Newgard (Duke University School of Medicine) and obtained by Dr. Duk-Su Koh (via Dr. Ian Sweet, both at University of Washington), who provided it to us. The *Eipr1* KO and EIPR1(+) rescued 832/13 cell lines were generated as described (Topalidou, Cattin-Ortolá, et al., 2018). The 832/13 and 832/13-derived cell lines were grown in RPMI 1640-GlutaMAX (GIBCO) medium supplemented with 10% FBS (RMBIO), 1 mM sodium pyruvate (GIBCO), 10 mM HEPES (GIBCO), 1X Pen/Strep (GIBCO), and 0.0005% beta-mercaptoethanol at 5% CO_2_ and 37°C. The PC12 cell line was obtained from Dr. Duk-Su Koh (University of Washington) and was grown in DMEM-GlutaMAX (GIBCO) medium supplemented with 10% horse serum (RMBIO), 5% FBS (RMBIO), and 1X Pen/Strep (GIBCO) at 5% CO_2_ and 37°C. All cell lines were shown to be mycoplasma-free by DAPI staining and PCR test (ABM, G238).

### Protein extraction from mouse tissues

Tissues were dissected from an adult female mouse. Bigger organs such as the liver and brain were placed in 3 ml of lysis buffer, and smaller organs such as the heart and intestine were placed in 1.5 ml of lysis buffer (Tissue-PE LB^TM^, G-Biosciences #786-181). These organs were then frozen in liquid nitrogen and kept at −80°C for 1 hour. Tissues were dissolved with the use of a homogenizer until they became liquid and then were further homogenized with mechanical breakage using a 22-gauge needle, applying 12-15 strokes. The tissues were then spun down at 20,000g in a microcentrifuge for 15 minutes, and the supernatant was collected and spun again for 5 more minutes. The supernatant was finally collected and stored at −80°C.

### Immunoblotting

Protein extracts were loaded on 8, 10, or 12% SDS-PAGE gels and transferred onto PVDF membranes. For the blots shown in Figure S2A,B, membranes were blocked in 3% milk in TBST (50 mM Tris pH 7.4, 150 mM NaCl, 0.1% Tween 20) for 1 hour at room temperature (RT) and stained with the relevant primary and secondary antibodies in 3% milk in TBST for 1 hour at RT or overnight at 4°C, followed by three 5-minute washes in TBST. For the blots shown in Figure 1C,D, Figure 3D, Figure 5A, and Figure S5A,B, membranes were blocked with Odyssey® Blocking Buffer (PBS) (#927-4000), incubated overnight at 4°C with primary antibody diluted in Odyssey® Blocking Buffer (PBS) supplemented with 0.2% Tween 20, washed with PBST (PBS + 0.1% Tween 20), incubated for 1 hour at room temperature in secondary antibody diluted in Odyssey® Blocking Buffer (PBS) supplemented with 0.2% Tween 20 and 0.01% SDS, and washed with PBST. A LI-COR processor was used to image the membranes. The antibodies used are listed in Tables S4 and S5.

### Generation of *Ccdc186* KO by CRISPR

We performed Cas9-mediated genome editing in 832/13 cells as described (Ran, Hsu, et al., 2013). We designed guide RNAs using the online CRISPR design tool (Ran, Hsu, et al., 2013) and selected a guide RNA that recognizes a sequence in the first exon of rat *Ccdc186*: 5’-GCGTGAGTCGTCCTTCAACTCGG-3’.

The guide RNA was cloned into the pSpCas9(BB)-2A-GFP vector as described (Ran, Hsu, et al., 2013). The efficiency of the cloned guide RNA was tested using the SURVEYOR nuclease assay according to the manufacturer’s instructions (Surveyor Mutation Detection kit, Transgenomic). We designed a homology-directed repair (HDR) template using the pPUR vector (Clontech) as a backbone and cloned approximately 1.5 kb *Ccdc186* homology arms upstream and downstream of the puromycin selection cassette. The HDR template was assembled using Gibson cloning.

To cotransfect the CRISPR plasmid and HDR template into 832/13 cells, 832/13 cells were grown in two 10-cm petri dishes to near confluency. Cells were cοtransfected with 7 µg CRISPR plasmid and 7 µg non-linearized HDR template using Lipofectamine 3000 according to the instructions (ThermoFisher). 48 hours after transfection, we removed the culture medium and replaced with new medium containing 1 µg/ml puromycin. The puromycin selection was kept until individual clones could be picked, grown in individual dishes, and tested for *Ccdc186* KO. These clones were screened for *Ccdc186* KO by immunofluorescence and subsequently by RT-PCR and Western blot.

### Lentiviral production, infection of cells, and selection of stable lines

The following method was used to create a *Ccdc186* KO cell line rescued by expression of the *Ccdc186* cDNA. Platinum-E (PlatE) retroviral packaging cells (a gift from Suzanne Hoppins) were grown for a couple of generations in DMEM-GlutaMAX (GIBCO) medium supplemented with 10% FBS (RMBIO), 1X Pen/Strep (GIBCO), 1 µg/ml puromycin, and 10 µg/ml blastocidin at 5% CO_2_ and 37°C. On day one, approximately 3.6 x 10^5^ PlatE cells per well were plated in a six-well dish in DMEM-GlutaMAX medium supplemented with 10% FBS and 1X Pen/Strep. On day two, a mix of 152 µl Opti-MEM (ThermoFisher), 3 µg CCDC186_pBabe-hygro DNA, and 9 µl Fugene HD transfection reagent (Promega) was incubated for 10 minutes at room temperature and transfected into each well. On day three, we removed the medium and replaced with new PlatE medium. On day four, approximately 1.5 x 10^5^ *Ccdc186* KO 832/13 cells per well were plated in a six-well dish in RPMI 1640-GlutaMAX, supplemented with 10% FBS, 1 mM sodium pyruvate, 10 mM HEPES, 1X Pen/Strep, and 0.0005% beta-mercaptoethanol. 3 µl of 8 mg/ml hexadimethrine bromide (Sigma) was added to each well. The supernatant of the PlatE cells (48-hour viral supernatant) was collected with a sterile syringe, passed through a 0.45 micron filter, and added to the *Ccdc186* KO cells. The *Ccdc186* KO cells were incubated for 5-8 hours at 5% CO_2_ and 37°C, then the medium was changed and replaced with new medium and the cells were incubated overnight at 5% CO_2_ and 37°C. On day five, the supernatant was removed from the *Ccdc186* KO cells and replaced with the supernatant from the PlatE cells (72-hour viral supernatant) after passing through a 0.45 micron filter. 3 µl of 8 mg/ml hexadimethrine bromide was added to each well and the cells were incubated for 5-8 hours. The medium was replaced with new 832/13 cell medium. On day six, the *Ccdc186* KO cells were collected, transferred into a 10-cm petri dish, and 200 µg/ml hygromycin was added. The cells were grown under hygromycin selection until individual clones could be picked and tested for CCDC186 expression.

### RT-PCR

WT and *Ccdc186* KO cells were grown in 10-cm tissue-culture dishes, washed twice with ice-cold PBS, and harvested in 1 ml of TRIzol (Invitrogen). Total RNA was isolated following the manufacturer’s protocol and cDNA was synthesized from 1 µg total RNA using the QuantiTekt Reverse Transcription kit (Qiagen). The primers used for detecting *Ccdc186* cDNA were:

F: 5’-ATGAAGATCAGGAGCAGATTTGAAG-3’

R: 5’-GGCCTTCTTCGTCCTCTGCTC-3’

### Immunofluorescence

Cells were seeded onto cover slips (Thomas Scientific #121N79) placed in 24-well cell culture plates. For GFP or epitope-tagged constructs, cells were transfected at least 24 hours after seeding with 500-800 ng of plasmid using Lipofectamine 2000 (ThermoFisher), according to the manufacturer’s instructions, for 24 to 48 hours. When indicated, cells were incubated for 25 minutes with 50 µg/ml Alexa488- or Alexa568-transferrin (Invitrogen #T12242 or # T23365) diluted in serum-free RPMI medium + 25 mM HEPES + 1% BSA before fixation. When indicated, cells were incubated with MitoTracker deep red FM (Thermofisher #M22426) at a final concentration of 300 nM for 15 minutes in complete medium at 37°C.

For antibody staining, cells were rinsed twice with PBS and fixed with 4% paraformaldehyde (made in PBS) for 20 minutes at room temperature, then rinsed twice with PBS and permeabilized with 0.1% Triton X-100 in PBS for 20 minutes or 0.5% Triton X-100 in PBS for 5 minutes at room temperature, and finally rinsed twice with PBS and placed in 5% milk in PBS for 1 hour at room temperature. Cells were stained with primary antibodies in 0.5% milk in PBS at room temperature for 1 hour, washed three times with PBS, and incubated in secondary antibody for 1 hour at room temperature. The cells were washed with PBS three times, mounted onto glass slides using Vectashield (Vector laboratories H1000) or Prolong Diamond (Life Technologies P36965), sealed with transparent nail polish and examined by fluorescence microscopy. Antibodies and their working dilutions are listed in Tables S4 and S5.

Most images were obtained using an Olympus Fluoview FV1200 confocal microscope with a 60X UPlanSApo oil objective (numerical aperture = 1.35). The acquisition software was Olympus Fluoview v4.2. When indicated, a Nikon Eclipse 80i wide-field compound microscope was used. For the quantification of the insulin, proinsulin, CPE, and pro-CPE distributions in *Ccdc186* KO cells (Figure 1F, S6B, S7B and S7D), Pearson’s correlation coefficients were determined from confocal images using Fiji and the coloc-2 plugin by taking maximum-intensity projections of whole cells and drawing a line around each individual cell. For the quantification shown in Figures 1G and S6C, z-stack images were obtained using a Nikon Eclipse 80i wide-field compound microscope. Maximum-intensity projections were quantified using Fiji. The final fluorescence intensity is the total fluorescence in a region of interest around the Golgi divided by the fluorescence of a region of the same size in the cytoplasm of the cell (Golgi/Cytoplasm). For the quantification shown in Figures 1H and S6D, z-stack images were obtained using an Olympus Fluoview FV1200 confocal microscope. Maximum-intensity projections were obtained by only keeping the slices that had TGN38 signal so insulin or CPE signal coming from the top and the bottom of the cell would not impact the quantification. A line was drawn around the TGN38 marker and Pearson’s correlation coefficients were obtained using FIJI and the coloc-2 plugin. For the quantification of colocalization with CCDC186 (Figure 2A), a square was drawn around the perinuclear area where CCDC186 is localized, and Pearson’s correlation coefficients were determined using Fiji and the JACOP plugin.

For super-resolution imaging, cells were prepared as described above. For SIM imaging, coverslips were mounted with DAPI-free Vectashield and imaged with a 3D SIM OMX SR microscope (GE Healthcare). For STED microscopy, secondary antibodies were used at higher concentration (see Table S5). Coverslips were mounted in DAPI-free fresh Prolong Diamond, sealed with twinsil® 22, and imaged with a Leica SP8 STED microscope.

### Insulin and proinsulin secretion assays

Cells were grown in 24-well plates to near confluency. Cells were washed twice with PBS and incubated for 1 hour in 200 µl per well resting buffer (5 mM KCl, 120 mM NaCl, 24 mM NaHCO_3_, 1 mM MgCl_2_, 15 mM HEPES pH 7.4). The medium was collected, cleared by centrifugation, and stored at −80°C. The cells were incubated for 1 hour in 200 µl per well stimulating buffer (55 mM KCl, 25 mM glucose, 70 mM NaCl, 24 mM NaHCO_3_, 1 mM MgCl_2_, 2 mM CaCl_2_, 15 mM HEPES pH 7.4). After stimulation, the medium was cleared by centrifugation and stored at −80°C. The cells were washed once with PBS, harvested in PBS, and extracted in 100 µl per well acid-ethanol solution (absolute ethanol:H_2_O:HCl, 150:47:3). The pH of the acid-ethanol solution was neutralized by addition of 20 µl of 1 M Tris base per 100 µl of acid ethanol and the samples were stored at −80°C.

Samples were assayed for insulin or proinsulin content using ELISA according to the instructions of the manufacturers (Rat/Mouse insulin ELISA, Millipore, #EZRMI-13K; Rat/Mouse proinsulin ELISA, Mercodia, #10-1232-01). Secreted insulin and proinsulin levels were normalized against total cellular protein concentration and were presented as fraction of the wild type under stimulating conditions (Figure 1A and S3A).

### Constitutive secretion assay

WT and *Ccdc186* KO cells were seeded on 12-well plates and grown to subconfluency. Cells were transfected with a plasmid expressing GFP fused to a signal peptide at its N-terminus (ssGFP) (Hummer et al., 2017). 36-48 hours later, cells were incubated for 1 hour at 37°C in the cell culture incubator with resting medium (114 mM NaCl, 4.7 mM KCl, 1.2 mM KH_2_PO_4_, 1.16 mM MgSO_4_, 20 mM HEPES, 2.5 mM CaCl_2_, 25.5 mM NaHCO_3_, 3 mM glucose). The 12-well plate was transferred to ice and the secretion medium was collected and centrifuged at 20,000g for 10 minutes at 4°C to remove the cell debris. The supernatant was collected in a separate tube. The pelleted cells were lysed on ice with 50 mM Tris-HCl, pH 8.0, 150 mM NaCl, 1% Triton X-100, 1 mM EDTA, protease inhibitor cocktail without EDTA (Pierce). The lysate was clarified by centrifugation at 20,000g at 4°C for 10 minutes and the supernatant was collected to a new tube. The fluorescence of the medium (secreted GFP) and the lysate (cellular GFP) were measured using a plate reader (Molecular Devices Spectramax Gemini XPS, excitation = 485 nm, emission = 525 nm, cutoff = 515 nm). Background fluorescence was determined using the medium and lysate from cells that were not transfected. The data are presented as the ratio of fluorescence in the medium to the fluorescence in the cell lysate.

### Exocytosis assay

832/13 cells stably expressing NPY-pHluorin were transfected (FuGene, Promega) with NPY-mCherry. At 1 d after transfection, cells were washed once with PBS, dislodged using media, and transferred onto poly-L-lysine coated 22 mm glass coverslips. After an additional 2 d, cells were reset for 2 hrs in low K+ Krebs-Ringer buffer with 1.5 mM glucose, washed once with low K+ Krebs-Ringer buffer with 1.5 mM glucose, and coverslips were transferred to an open imaging chamber (Life Technologies). Cells were imaged close to the coverslips, focusing on the plasma membrane (determined by the presence of NPY-mCherry-positive plasma membrane docked vesicles), using a custom-built Nikon spinning disk confocal micrscope at a resolution of 512 × 512 pixels. Images were collected for 100 ms at 10 Hz at room temperature with a 63X objective (Oil Plan Apo NA 1.49) and an ImageEM X2 EM-CCD camera (Hamamatsu, Japan). Following baseline data collection (15 s), an equal volume of Krebs-Ringer buffer containing 110 mM KCl and 30.4 mM glucose was added to stimulate secretion and cells were imaged for an additional 80 s. At the end of the experiment, cells were incubated with Krebs-Ringer buffer containing 50 mM NH_4_Cl, pH 7.4, to reveal total fluorescence and to confirm that the imaged cells were indeed transfected. Movies were acquired in MicroManager (UCSF) and exported as tiff files. Movies were analyzed using ImageJ software by counting events and measuring cell area.

### pH measurement of the late-Golgi compartment

WT and *Ccdc186* KO cells stably expressing the 17-residue transmembrane domain of beta-galactoside alpha 2,6-sialyltransferase tagged to pHluorin (St6Gal1::pHluorin) were generated as follows. HEK293T cells were maintained in DMEM with 10% fetal bovine serum under 5% CO_2_ at 37°C. Lentivirus was produced by transfecting HEK293T cells with FUGW, psPAX2, and pVSVG using Fugene HD according to the manufacturer’s instructions. Two days after transfection, the medium was collected and filtered (0.45 µm). Medium containing the lentivirus was then applied to 832/13 cells in suspension at varying ratios. Transduction efficiency was screened using an epifluorescence microscope. The protocol to measure the pH of the late-Golgi compartment (Hummer et al., 2017) was adapted to a plate-reader format. An equal number of WT and *Ccdc186* KO 832/13 cells stably expressing St6Gal1::pHluorin were plated on clear-bottom black-wall 96-well plates and grown to confluence. Cells were washed once with Tyrode buffer (119 mM NaCl, 2.5 mM KCl, 2 mM CaCl_2_, 2 mM MgCl_2_, 25 mM HEPES, 30 mM glucose, pH 7.4) and incubated for 10 minutes either in Tyrode buffer or in KCl-enriched buffer at different pHs (125 mM KCl, 20 mM NaCl, 0.5 mM CaCl_2_, 0.5 mM MgCl_2_, and 25 mM MES (at pH 5.5 or 6) or 25 mM HEPES (at pH 6.5, 7, 7.5, 8, or 8.5). The KCl-enriched buffer was supplemented with 5 µM nigericin (Sigma Aldrich) and 5 nM monensin (Sigma Aldrich). The fluorescence of each well was measured using a Varioskan Lux plate reader (Thermo Scientific) (Excitation=485 nm, Emission=520 nm). For each buffer, cell type, and independent biological replicate, the reading was repeated three times. Calibration curves were generated for each cell type and each independent repetition. The absolute pH values were extrapolated from the calibration curve using the fluorescence of the cells incubated with Tyrode buffer.

### ANF-GFP pulse-chase experiment

To monitor the exit of DCV cargo from the TGN, we used a protocol similar to the one described (Kögel et al., 2013; Kögel, Rudolf, et al., 2010). WT and *Ccdc186* KO 832/13 cells were seeded on glass coverslips in 24-well plates and after at least 24 hours, 100 ng of ANF::GFP plasmid was transfected in each well with Lipofectamine 2000 for 12-16 hours at 37°C in complete medium. Subsequently, cells were incubated at 20°C in PBS for 2 hours in a conventional incubator (pulse) to block protein exit from the TGN. 30 minutes before the end of the low temperature block, 10 µg/ml of cycloheximide was added to the medium to block the synthesis of new ANF::GFP. The PBS was then exchanged for growth medium and the cells were shifted to 37°C for the indicated times, then fixed with 4% PFA, and stained with an anti-GFP antibody as described (see Immunofluorescence section). Cells were scored in three categories: those that had most of the ANF::GFP concentrated at the TGN (“Golgi-like”), those that had ANF::GFP both at the TGN-region and at the cell periphery (“Intermediate”) and those where the ANF::GFP was excluded from the TGN (“Periphery”). 50 to 100 cells per time point and per genotype were imaged and counted using a Nikon Eclipse 80i wide-field compound microscope. The experimenter was blind to the genotypes of the cell lines used and to the time point. The experiment was repeated three times with similar results.

### Generation of PC12 cell lines for the mitochondria relocation assays

For the mitochondria relocation assay, PC12 cells were used instead of 832/13 cells for drug selection compatibility reasons. Moreover, toxicity of the mitochondria relocation constructs appeared to be less in PC12 cells. A 6-well plate was transfected with 4 µg of the appropriate plasmid using Lipofectamine 2000 (ThermoFisher) according to the manufacturer’s instructions. After 48 hours, the cell culture medium was replaced with Pen/Strep-free selection medium (DMEM-GlutaMAX (GIBCO) supplemented with 5% FBS (RMBIO), 10% horse serum (RMBIO), and 700 µg/ml G418). The selection medium was changed every 2-3 days until large single colonies could be seen. Colonies were transferred to a 96-well plate and expanded in selection medium and then to 24-well and 6-cm plates before being frozen in 10% DMSO. Individual clones were screened by immunostaining using an anti-HA antibody. Highly expressing clones were used for the BioID experiment.

### BioID experiment and sample preparation for mass spectrometry

The protocol for isolating proteins biotinylated by BirA* was adapted from (Shin, Gillingham, et al., 2017). PC12 stable lines expressing the BirA* constructs (BirA*::HA::MAO and CC3::BirA*::HA::MAO) were grown to confluence in three 175-cm^2^ flasks in complete PC12 DMEM medium (DMEM-GlutaMAX (GIBCO) supplemented with 5% FBS (RMBIO) and 10% horse serum (RMBIO)). 24 hours before harvest, cells were incubated with 50 µM biotin (10 mg/ml stock in DMSO). Cells were transferred on ice, washed 2x with ice-cold PBS, and harvested by gentle pipetting. Cells were pelleted by centrifugation (10 minutes, 1850g) and resuspended in lysis buffer (50 mM Tris pH 7.4, 0.1M NaCl, 1 mM EDTA, 1% Triton X-100, 1 mM PMSF, protease inhibitor cocktail (Pierce)), vortexed briefly, incubated on ice for 20 min, and vortexed again. Lysates were clarified by centrifugation at 20,000g for 10 minutes at 4°C and supernatants were mixed with an equal volume of 50 mM Tris pH 7.4. The total protein content was about 30 mg as measured by a BCA assay. The supernatant was incubated with 250 µl Dynabeads MyOne Streptavidin C1 beads (Invitrogen # 65002) that had been pre-washed twice in the same buffer. The beads were incubated at 4°C overnight on a Nutator, washed twice for 8 minutes in 2% SDS–PAGE and protease inhibitor cocktail (Pierce), then three times for 8 min in 1% Triton X-100, 0.1% deoxycholate, 500 mM NaCl, 1 mM EDTA, 50 mM HEPES, and protease inhibitor cocktail at pH 7.5, and three times for 8 minutes in 50 mM Tris pH 7.4, 50 mM NaCl, and protease inhibitor cocktail. Finally, the beads were incubated for 5 minutes at 98°C with 50 µl 1X lithium dodecyl sulfate (LDS) sample buffer (Invitrogen NuPAGE® #NP0007) containing 10% beta-mercaptoethanol and 3 mM biotin. The supernatant was collected and the beads were discarded.

The samples were then prepared for mass spectrometry as follows. The eluted proteins in 1X LDS were reduced with 1 mM Tris(2-carboxyethyl)phosphine (TCEP) at 37°C for 20 minutes on a shaker and alkylated with 2 mM chloroacetamide for 20 minutes at 37°C. The reaction was quenched by incubation with 1 mM TCEP for 20 minutes at 37°C and 60% of the eluate was run on an SDS-PAGE gel (Mini-Protean TGX, 4-15%, Bio-Rad #4561084). The gel was stained with colloidal Coomassie (Bio-Rad #1610803) overnight at room temperature and destained with Milli-Q water. Gel slices (4-5 slices per lane) were excised and destained by two 10-minute washes with a 50/50 solution of high-grade ethanol/100 mM triethylammonium bicarbonate (TEAB). The gel was dehydrated with two washes of high-grade ethanol. Gel slices were rehydrated with a solution of 12.5 ng/µl trypsin (Promega Trypsin Gold, Mass Spectrometry Grade, #PRV5280) diluted in 100 mM TEAB. Once rehydrated, 2-3 volumes of 100 mM TEAB were added and the sample was incubated overnight at 37°C in a shaking thermomixer at 1,400 rpm. The reaction was quenched by addition of 1/5 10% trifluoroacetic acid (TFA) and digested peptides were purified by StageTips (Rappsilber et al., 2007). StageTips were washed with 50 µl methanol, 50 µl reaction buffer B (0.1% TFA, 80% acetonitrile (ACN)), and equilibrated with 50 µl of buffer A (5% ACN, 0.1% TFA in water) by centrifugation at 2,000g at room temperature. The samples were loaded onto StageTips that were then washed once with 50 µl buffer A and kept at 4°C until mass spectrometry analysis.

### Liquid chromatography mass spectrometry (LC/MS) analysis

Peptides were eluted from StageTips using elution buffer (50% acetonitrile, 0.1% TFA) and then loaded on a self-pulled 360 µm outer diameter (OD) x 100 µm inner diameter (ID) 20 cm column with a 7-µm tip packed with 3 µm Reprosil C18 resin (Dr. Maisch, Germany). Peptides were analyzed by nanoLC-MS in a 120 minutes, 5% to 35% acetonitrile gradient in 0.1% acetic acid at 300 nl/min (Thermo Dionex RSLCnano) on an Orbitrap Elite. Orbitrap Fourier Transform Mass Spectrometry (FTMS) spectra (R = 30,000 at 400 m/z; m/z 350–1600; 3e6 target; max 500 ms ion injection time) and Top15 data dependent collision-induced dissociation (CID) MS/MS spectra (1e4 target; max 100 ms injection time) were collected with dynamic exclusion for 20 sec and an exclusion list size of 500. The normalized collision energy applied for CID was 35% for 10 msec.

### Data analysis

Mass spectra were searched against the Uniprot rat reference proteome downloaded on Feburary 12, 2017 using MaxQuant v1.5.7.4. Detailed MaxQuant settings used were “LFQ mode” with “Fast LFQ”; and “Match between runs” mode was enabled. Other settings were kept as default.

Data analysis was carried out in R. Briefly, label-free quantification (LFQ, (Cox et al., 2014) intensities calculated by MaxQuant were log_2_ and Z-score transformed; only proteins present in at least two out of three replicates were selected for further analysis; missing values were replaced by random numbers drawn from a distribution with mean of −3.51 and a standard deviation of 0.3 to represent protein abundance below the detection limit; a two-sided Student’s t-test was used to determine the *p*-values and the results are presented in the form of a volcano plot (Figure S13). Proteins with ratios (log_2_ CCDC186 CC3 – log_2_ BirA*) higher than 1.8 are labeled. The data for all mass spectrometry experiments are shown in Tables S6 and S7.

### Equilibrium sedimentation

Wild type, *Ccdc186* KO, and *Eipr1* KO 832/13 cells were grown in 15-cm tissue culture plates. Cells were washed twice with ice-cold PBS, transferred to 15 ml conical tubes, and centrifuged at 300g for 10 minutes at 4°C. Cells were washed once with 5 ml of SH buffer (10 mM HEPES pH 7.2, 0.3 M sucrose, 1 mM PMSF and protease inhibitor cocktail (SIGMAFAST)) and resuspended in 1 ml SH buffer, passed through a 21-gauge needle, and homogenized using a ball bearing device (Isobiotec, 18 µm clearance). The lysate was collected and centrifuged at 1000g for 8 minutes. A 0.6-1.9 M continuous sucrose gradient was prepared in 10 mM HEPES pH 7.2 in ultracentrifuge tubes (Beckman Coulter) that were coated with Sigmacote (Sigma #SL2). The post-nuclear supernatant was loaded onto the gradient and centrifuged at 30,000 rpm in an SW41 rotor for 14-16 hours at 4°C. Fractions (750 µl each) were collected from top to bottom. The sucrose concentration for each fraction was determined by measuring the Brix using a Misco palm abbe digital refractometer. Fractions were analyzed by immunoblotting using antibodies to PC1/3 and CPD (see above and Tables S4 and S5). Band intensity was quantified using FIJI and plotted against the sucrose concentration of the fraction.

### Statistics

Data were tested for normality by a Shapiro-Wilk test. When data did not pass the normality test, we tested for statistical significance between groups by the Kruskal-Wallis test followed by Dunn’s test when making comparisons among groups of three or more, and the Mann-Whitney test for comparisons between groups of two. When data passed the normality test, we tested for statistical significance by a one-way ANOVA test with Bonferroni correction when making comparisons among groups of three or more, and an unpaired t-test for comparisons between groups of two.

## Abbreviations

DCV: dense-core vesicle
TGN: trans-Golgi network
EARP: endosome-associated recycling protein
iDCV: immature dense-core vesicle
CPE: carboxypeptidase E
PC1/3: proprotein convertase 1/3
CgA: chromogranin A
MAO: monoamine oxidase
CPD: carboxypeptidase D
CD-MPR: cation-dependent mannose 6-phosphate receptor

## ACKNOWLEDGEMENTS

We thank Alex Merz for mentoring and sharing equipment; Christopher Newgard, Ian Sweet, and Duk-Su Koh for the 832/13 cell line; Suzanne Hoppins for the pBabe vector, platE cells, protocol for lentiviral production and infection, help with cell culture, and sharing equipment; Richard Palmiter for the pPUR vector; Sean Munro and John Shin for the mitochondrial relocation and BirA* plasmids, advice, and protocols for BioID; Lloyd Fricker for the anti-CPD and anti-CPE antibodies; Joshua Vaughan and Aaron Halpern for secondary antibodies used in STED microscopy; Patrina Pellett and David Castaneda-Castellanos for assistance with SIM and STED microscopy, respectively; Ngoc-Han Thi Nguyen and Donna Prunkard from the UW Pathology Flow Cytometry Core Facility; Ken Miller and Joshua Kaplan for worm strains with integrated transgenes; and Kelly Duong for the mouse tissues and protocol for protein extraction. This work used an EASY-nLC1200 UHPLC and Thermo Scientific Orbitrap Fusion Lumos Tribrid mass spectrometer purchased with funding from a National Institutes of Health SIG grant S10OD021502 (S-E.O.). Some worm strains were provided by the CGC, which is funded by NIH Office of Research Infrastructure Programs (P40 OD010440). This work was supported by American Diabetes Association grant #1-17-JDF-064 and by NIH grant R01 GM124035 to CSA, and by a University of Washington Diabetes Research Center Pilot and Feasibility Award (NIH grant P30 DK017047) and by NIH grant R01 GM121481 to MA.

## SUPPLEMENTARY FIGURE LEGENDS

**Figure S1.**
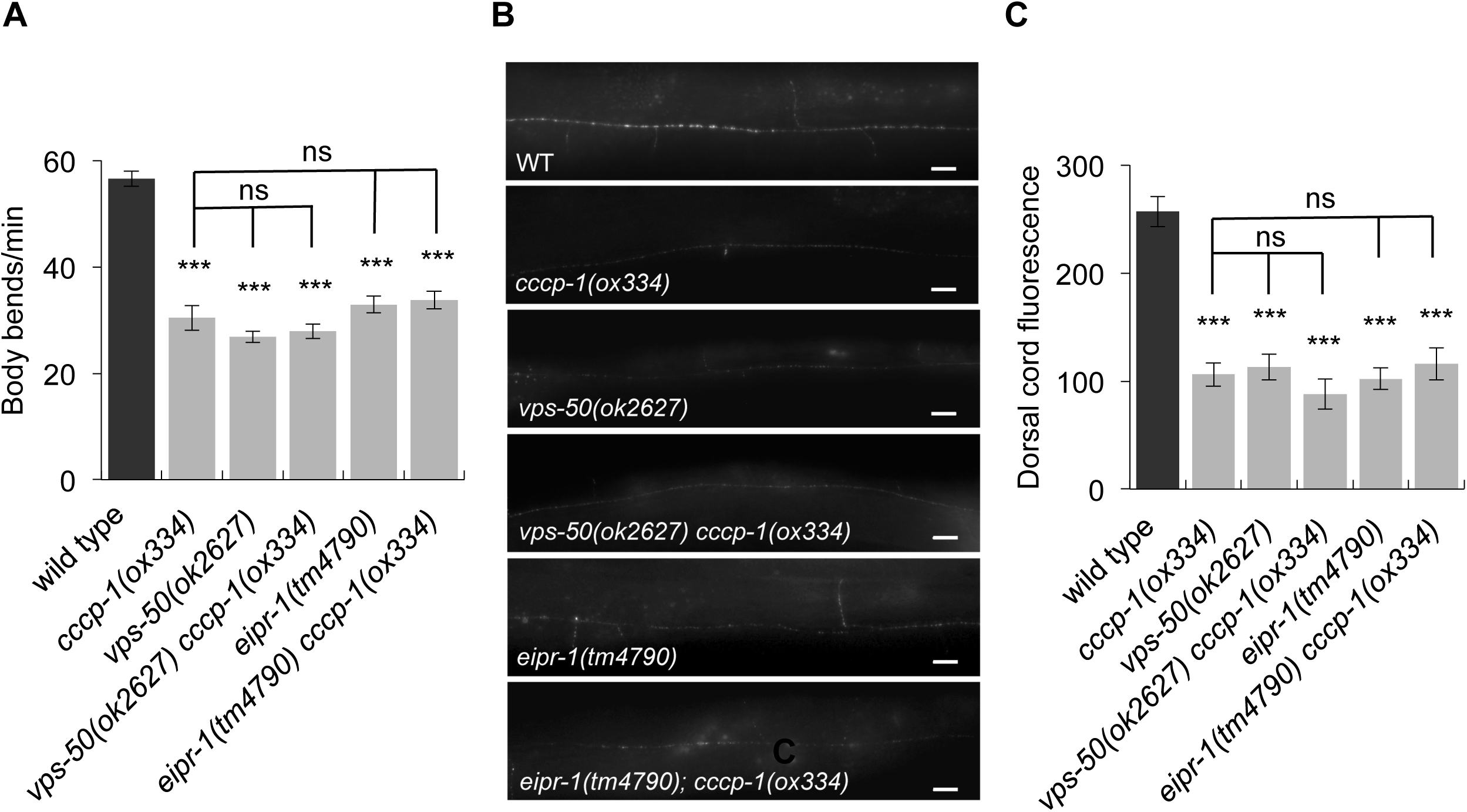
*cccp-1* functions in the same pathway as *eipr-1* and *vps-50* to control locomotion and dense-core vesicle cargo sorting in *C. elegans*. A. *cccp-1* acts in the same pathway as *eipr-1* and *vps-50* to control locomotion. The *cccp-1*(*ox334*) mutation does not enhance the slow locomotion phenotype of either *eipr-1*(*tm4790*) or *vps-50*(*ok2627*) mutants. Error bars = SEM; n = 12-22. B. *cccp-1* acts in the same genetic pathway as *eipr-1* and *vps-50* to control DCV cargo trafficking. Representative images of NLP-21::Venus fluorescence in motor neuron axons of the dorsal nerve cord. Maximum-intensity projections. Scale bar: 10 µm. C. Quantification of NLP-21::Venus fluorescence levels in the dorsal nerve cord. Double mutants of *cccp-1*(*ox334*) with *eipr-1*(*tm4790*) or *vps-50*(*ok2627*) are not significantly different from the single mutants (***, p<0.001; ns, not significant, p>0.05). Error bars = SEM; n = 10-15.

**Figure S2.**
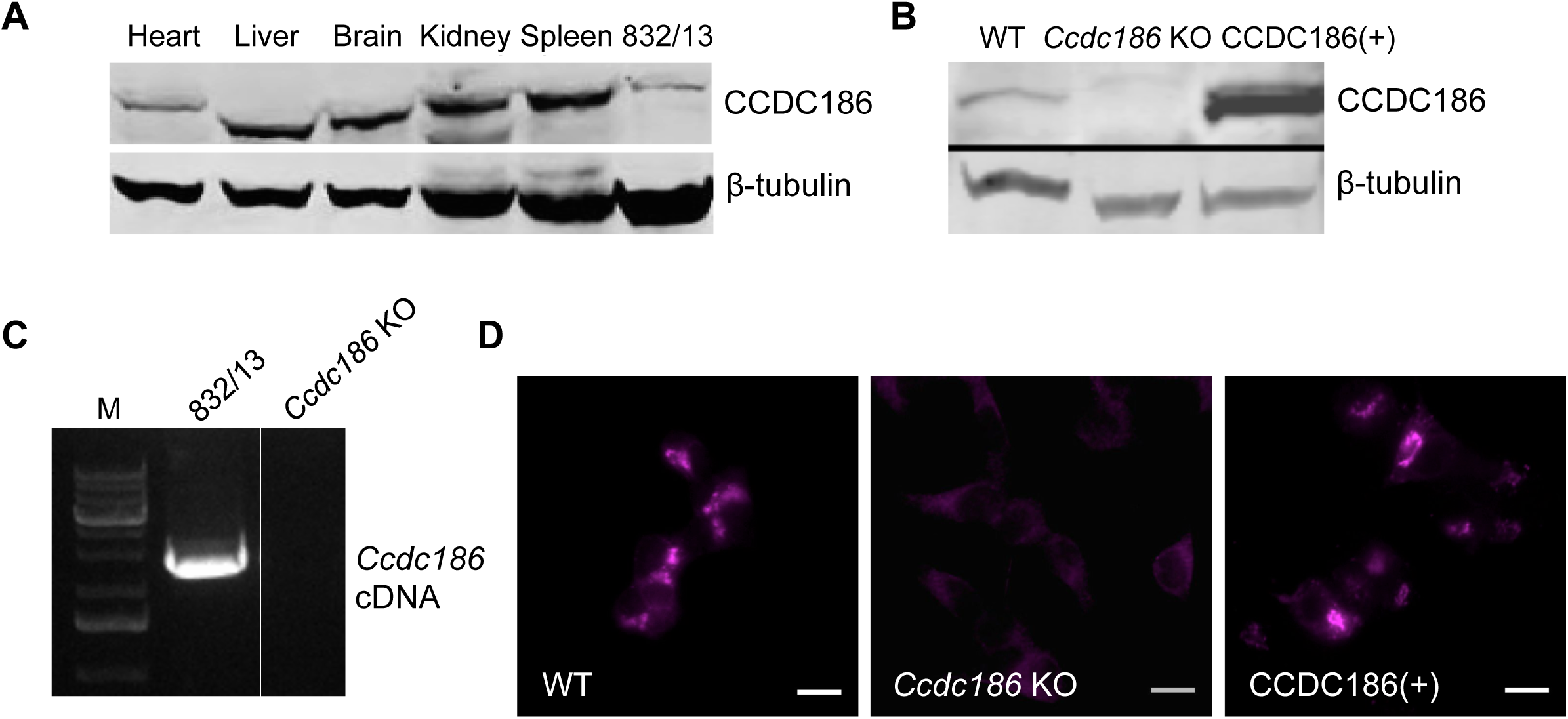
Generation of a *Ccdc186* KO 832/13 beta-cell line. A. CCDC186 is expressed in multiple mouse tissues. Protein extracts from different mouse tissues were blotted for CCDC186 using a commercial antibody. β-tubulin was used as a loading control. In two independent blots run from the same protein samples, we saw the same small differences in the size of the CCDC186 band in different tissues. B. *Ccdc186* KO cells do not express CCDC186. Protein extracts from WT, *Ccdc186* KO, and *Ccdc186* KO 832/13 cells stably expressing wild type CCDC186 (*Ccdc186*(+)) were blotted with anti-CCDC186 antibody. The faint band of slightly larger size in the *Ccdc186* KO sample was not seen in other repetitions of this experiment and is presumed to be a nonspecific band. β-tubulin was used as a loading control. C. Total cDNA from WT 832/13 and *Ccdc186* KO cells was PCR amplified using primers that detect the *Ccdc186* cDNA. D. WT, *Ccdc186* KO, and *Ccdc186*(+) rescued cells were stained with anti-CCDC186 antibody. Single slices. Wide-field fluorescence compound microscopy. Scale bars: 10 µm.

**Figure S3.**
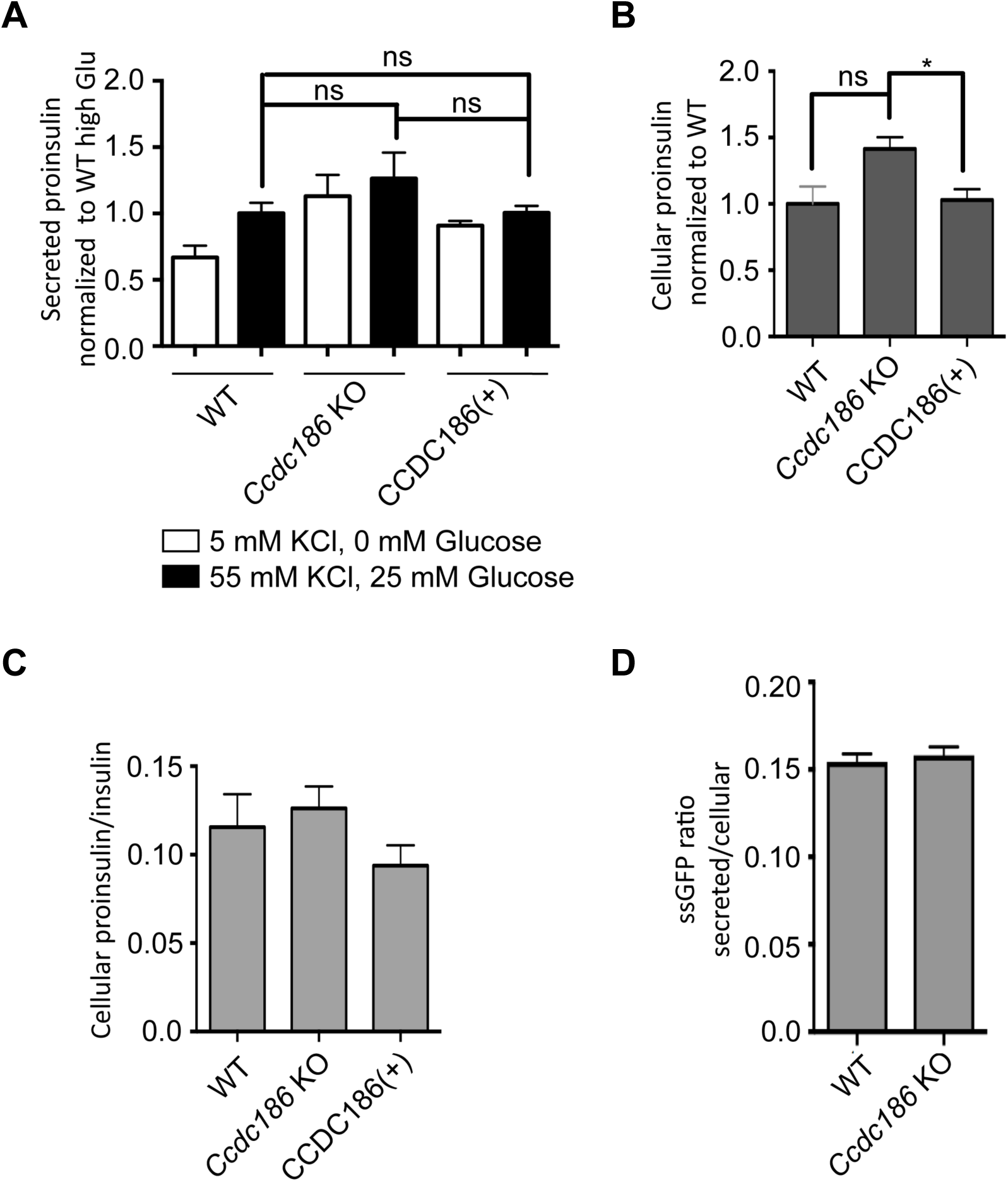
Proinsulin secretion and constitutive secretion are not affected in *Ccdc186* knock-out cells. A. Proinsulin secretion from WT, *Ccdc186* KO and *Ccdc186*(+) rescued 832/13 cells under resting and stimulating conditions. All values were normalized to the WT value in stimulating conditions (WT high Glu). n = 6; ns p>0.05, error bars = SEM. B. Total proinsulin content in WT, *Ccdc186* KO, and *Ccdc186*(+) rescued 832/13 cells. All values were normalized to WT. n = 6; *p<0.05, ns p>0.05, error bars = SEM. We performed three biological replicates. For each replicate, the same cells were used to determine the amount of proinsulin secreted under resting conditions, stimulating conditions, and the amount of total cellular proinsulin. C. CCDC186 is not required for proinsulin processing. Ratio of the cellular proinsulin content to the total cellular insulin content in WT, *Ccdc186* KO, and *Ccdc186*(+) rescued 832/13 cells. We performed two biological replicates. For each replicate, the amounts of total cellular proinsulin and total cellular insulin were determined from the same cells. n = 6, error bars = SEM. D. *Ccdc186* KO cells have normal constitutive secretion. Secretion of ssGFP (GFP fused at its N-terminus to the signal peptide of rat ANF). The data shown are combined from two independent experiments with similar results. n=6, error bars = SEM. The data shown for the WT are the same shown in Figure 1F of (Topalidou, Cattin-Ortolá, et al., 2019, manuscript submitted to *MBoC*) since these experiments were run in parallel with the same WT control.

**Figure S4.**
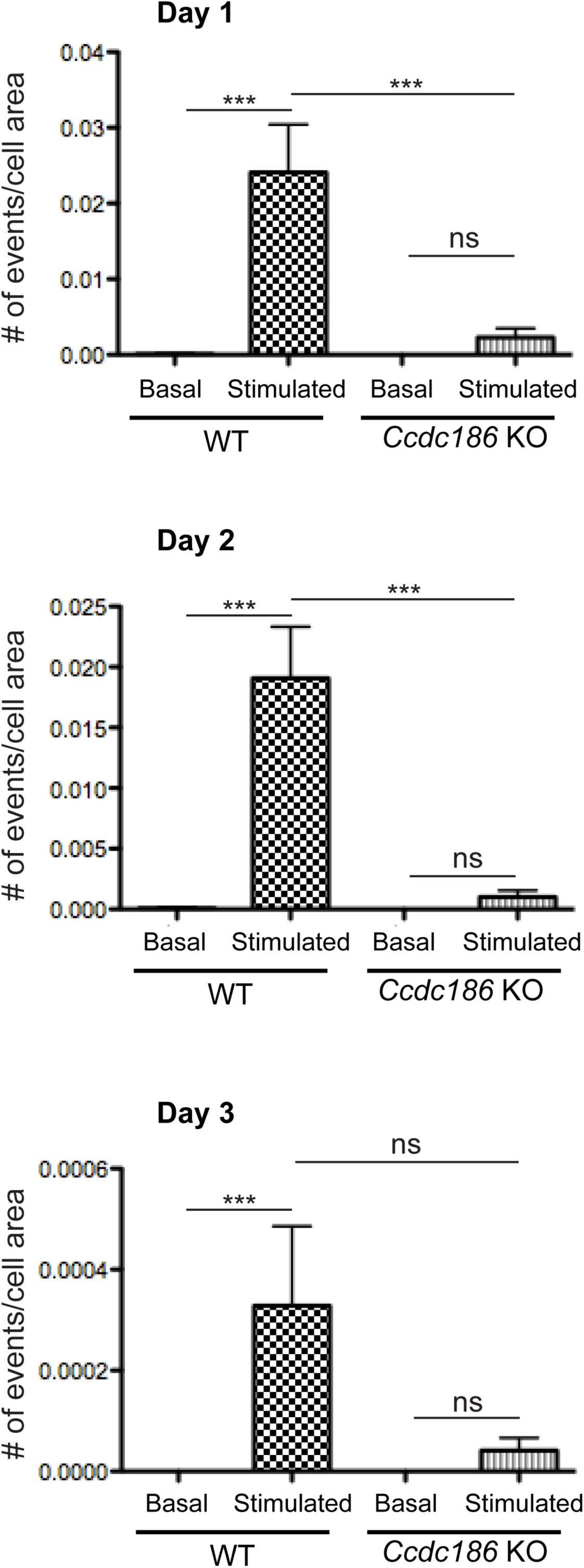
*Ccdc186* knock-out cells have exocytosis defects. WT and *Ccdc186* KO 832/13 cells stably expressing NPY-pHluorin were reset for 2 hrs in low K+ Krebs-Ringer buffer with 1.5 mM glucose. Cells were then imaged in low K+ Krebs-Ringer buffer with 1.5 mM glucose for 15 s using spinning-disk confocal microscopy. Cells were then stimulated with 60 mM K+ and 16.7 mM glucose for 80 s and imaged again. Images (100 ms exposure) were collected at 10 Hz. Exocytotic events were hand-counted. Bar graphs show the number of exocytotic events per second normalized to cell surface area. *ns*<>0.05, \**p* < 0.05. Three experiments were performed on different days and plotted separately. The *Ccdc186* KO showed reduced stimulated exocytosis on days 1 and 2, but no significant difference from WT on day 3. However, very few events were observed on day 3, even for WT (note the scale of the y-axis), suggesting problems on that day with the stimulation protocol or ability to detect events. On day 1, n=47 cells for WT (four coverslips imaged separately with 20, 12, 7, and 8 cells); n=42 cells for *Ccdc186* KO (four coverslips with 10, 15, 7, and 10 cells). On day 2, n=79 cells for WT (four coverslips with 26, 13, 22, and 18 cells); n=28 cells for *Ccdc186* KO (two coverslips with 10 and 18 cells). On day 3, n=30 cells for WT (two coverslips with 12 and 18 cells); n=21 cells for *Ccdc186* KO (two coverslips with 15 and 6 cells). The data shown for the WT are the same shown in Figure S3 of (Topalidou, Cattin-Ortolá, et al., 2019, manuscript submitted to *MBoC*) since these experiments were run in parallel with the same WT control.

**Figure S5.**
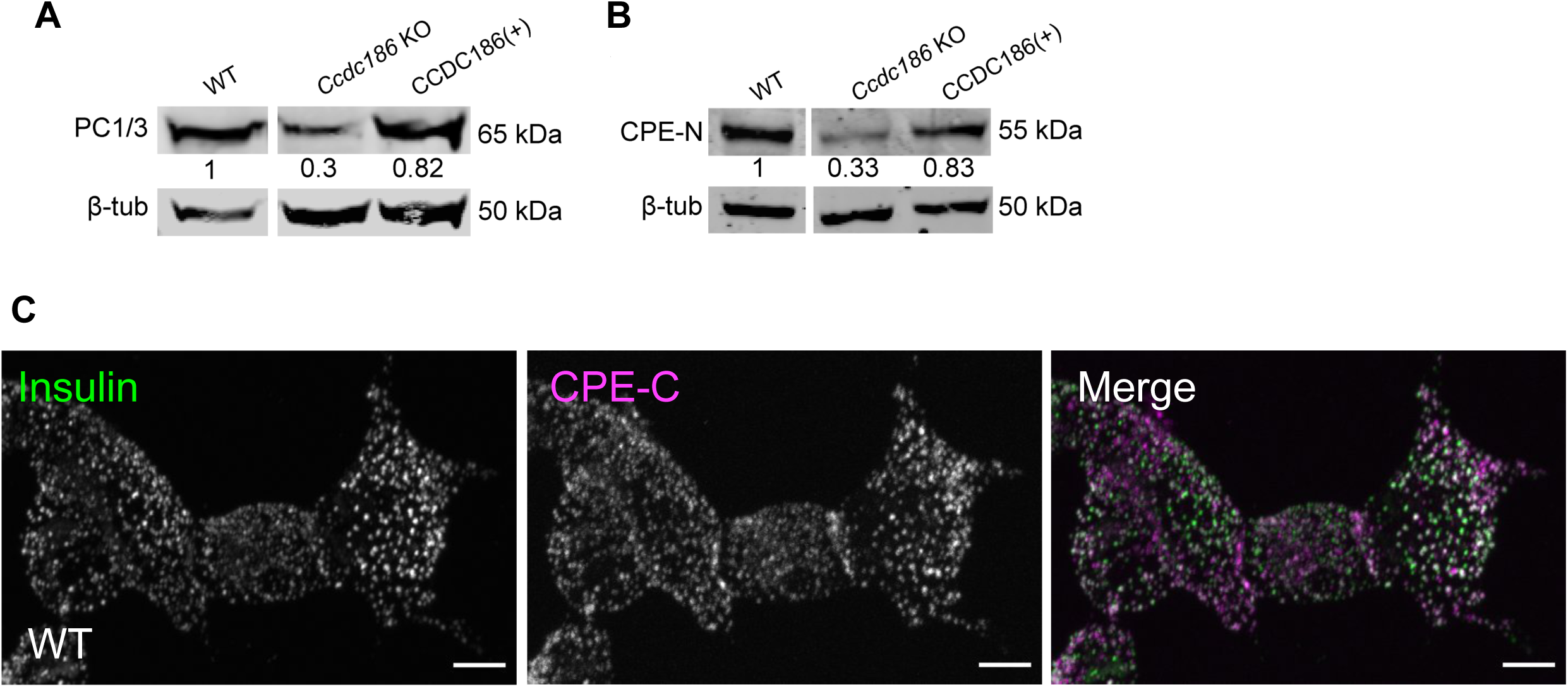
The levels of mature PC1/3 and CPE are decreased in *Ccdc186* KO cells and mature CPE colocalizes with insulin in WT cells. A. *Ccdc186* KO cells have reduced levels of the processed form of the proprotein convertase PC1/3. Detergent lysates from WT, *Ccdc186* KO, and *Ccdc186*(+) rescued 832/13 cells were blotted with an antibody to PC1/3. β-tubulin was used as a loading control. The experiment was repeated four times with similar results. B. *Ccdc186* KO cells have reduced levels of the processed form of CPE. Detergent lysates from WT, *Ccdc186* KO, and *Ccdc186*(+) rescued 832/13 cells were blotted with an antibody to the N-terminus of CPE (CPE-N). β-tubulin was used as a loading control. The experiment was repeated three times. The data shown for the WT are the same shown in Figure 3C of (Topalidou, Cattin-Ortolá, et al., 2019, manuscript submitted to *MBoC*) since these experiments were run in parallel with the same WT control. C. CPE and insulin almost fully colocalize. Representative confocal images of 832/13 cells costained for endogenous insulin and endogenous CPE using the CPE-C antibody. Maximum-intensity projections. Scale bars: 5 µm.

**Figure S6.**
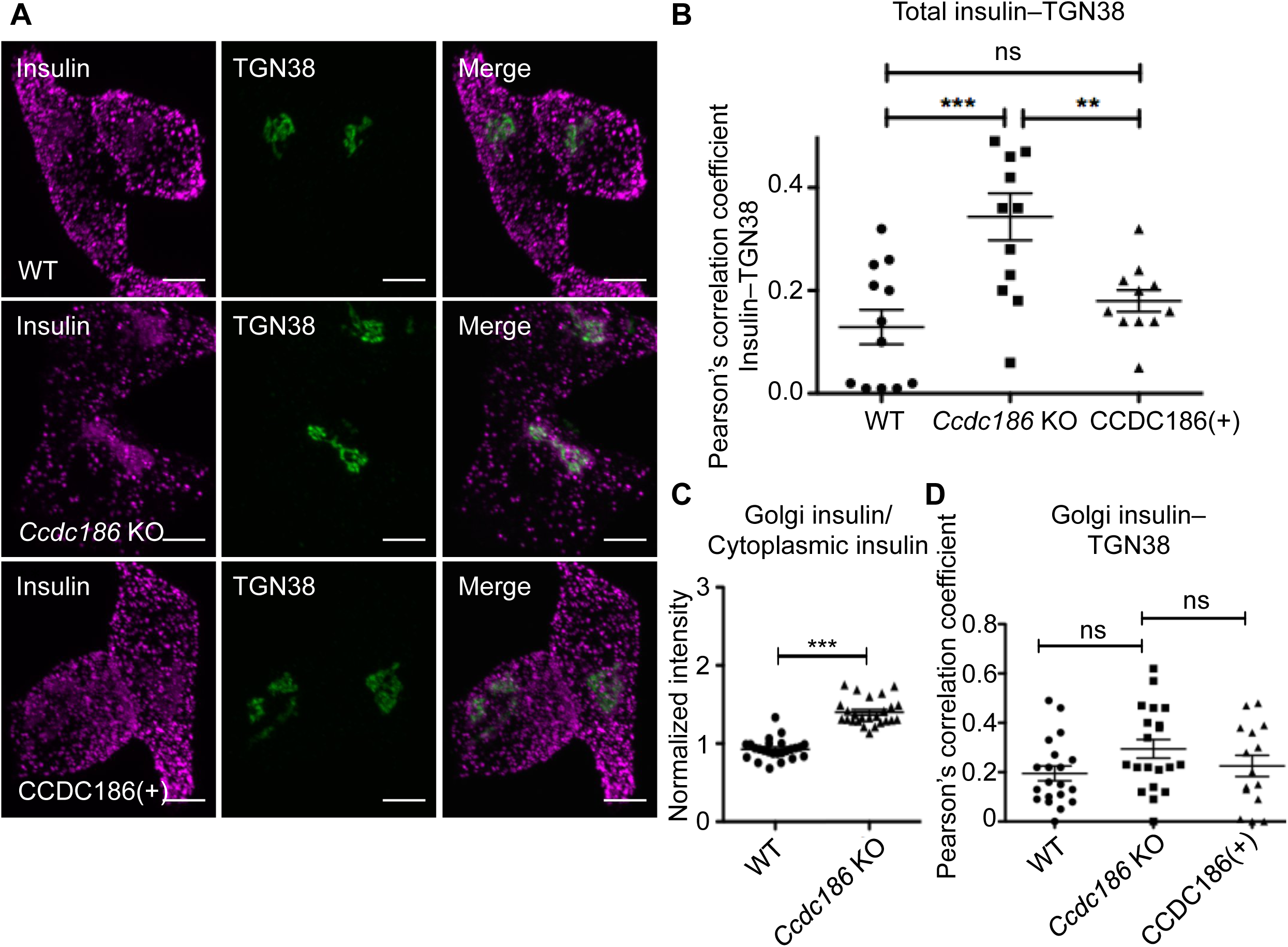
*Ccdc186* knock-out cells have defective distribution of insulin. A. Insulin is localized less in the periphery and more at or near the TGN in *Ccdc186* KO cells. Representative confocal images of WT, *Ccdc186* KO, and *Ccdc186*(+) rescued 832/13 cells costained for insulin and TGN38. Maximum-intensity projections. Scale bars: 5 µm. We performed three biological replicates. B. Quantification of the colocalization between insulin and the TGN marker TGN38. Maximum-intensity projection images were obtained and Pearson’s correlation coefficients were determined by drawing a line around each cell. The data shown are from one representative experiment of three biological replicates with similar results. n=12 for WT and *Ccdc186* KO; n=11 for *Ccdc186*(+), **p<0.01, ***p<0.001, ns p>0.05, error bars = SEM. C. *Ccdc186* KO cells have an increased proportion of insulin localized near the TGN. Fluorescence of a region of interest around the TGN divided by the fluorescence of a region of the same size in the cytoplasm. The data shown are from one experiment. n=25 for WT and *Ccdc186* KO, ***p<0.001, error bars = SEM. The data shown for the WT are the same shown in Figure 2C of (Topalidou, Cattin-Ortolá, et al., 2019, manuscript submitted to *MBoC*) since these experiments were run in parallel with the same WT control. D. Quantification of insulin localized at or near the TGN. Maximum-intensity projection images of z-stacks encompassing the TGN38 signal were obtained and Pearson’s correlation coefficients were determined by drawing a line around the TGN38 positive signal. The data shown are from one representative experiment of three biological replicates with similar results. n=20 for WT and *Ccdc186* KO, n=15 for *Ccdc186*(+); ns p>0.05; error bars = SEM.

**Figure S7.**
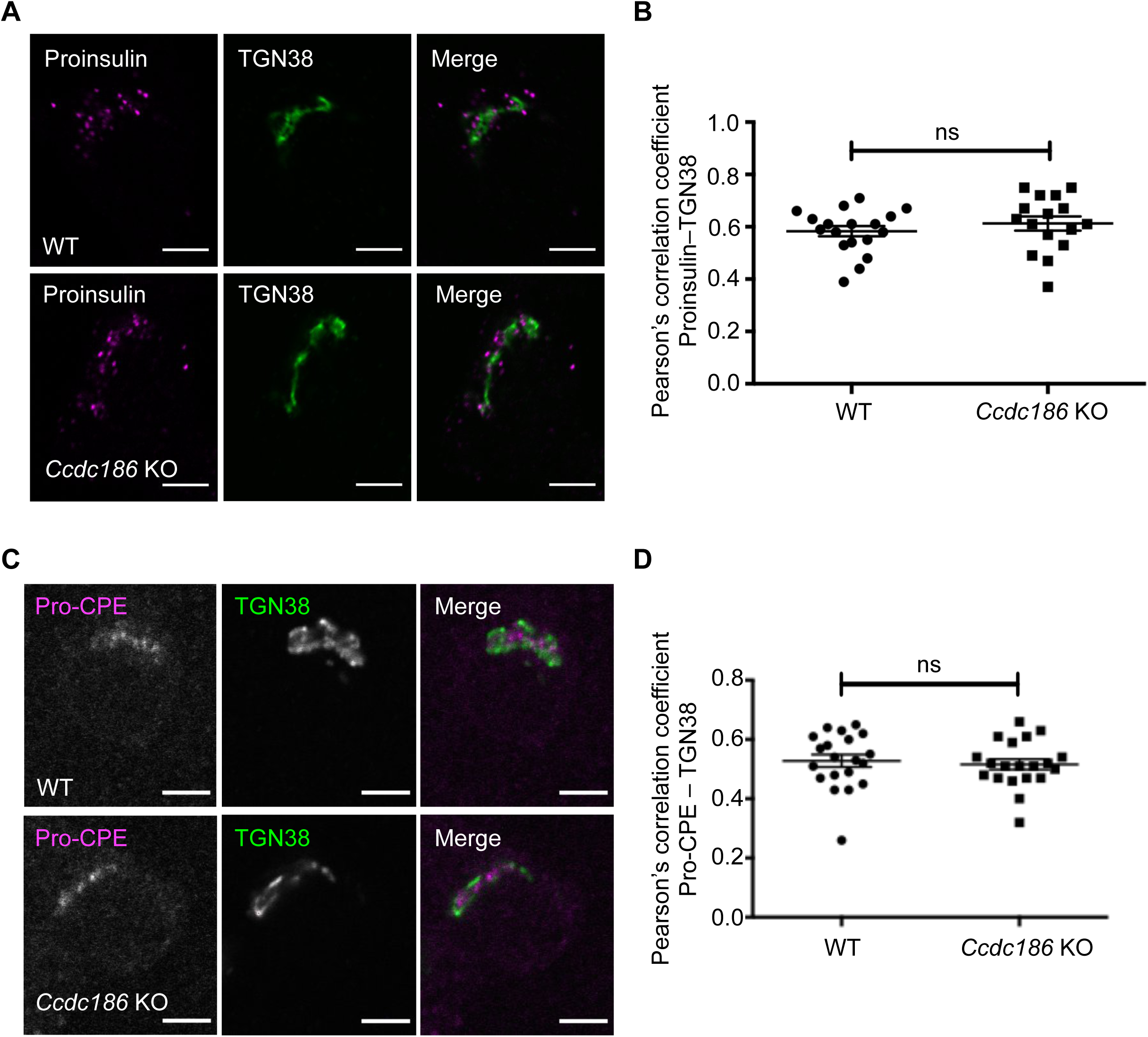
*Ccdc186* knock-out cells have normal distributions of proinsulin and unprocessed CPE. A. The localization of proinsulin is not affected by the absence of CCDC186. Representative images of WT and *Ccdc186* KO 832/13 cells costained for endogenous proinsulin and TGN38. Single slices. Scale bars: 5 µm. We performed three biological replicates. B. Quantification of the colocalization between proinsulin and the TGN marker TGN38. The data shown are from one representative experiment of three biological replicates with similar results. n=18 for WT and n=16 for *Ccdc186* KO, ns p>0.05, error bars = SEM. The data shown for the WT are the same shown in Figure 2G of (Topalidou, Cattin-Ortolá, et al., 2019, manuscript submitted to *MBoC*) since these experiments were run in parallel with the same WT control. C. The unprocessed form of CPE (pro-CPE) is localized at or near the TGN in both WT and *Ccdc186* KO cells. Representative confocal images of WT and *Ccdc186* KO cells costained for pro-CPE antibody and TGN38. Single slices. Scale bars: 5 µm. The experiment was repeated twice. D. Quantification of the colocalization between pro-CPE and the TGN marker TGN38. The data shown are from one experiment. n=20 for both WT and *Ccdc186* KO, ns p>0.05, error bars = SEM.

**Figure S8.**
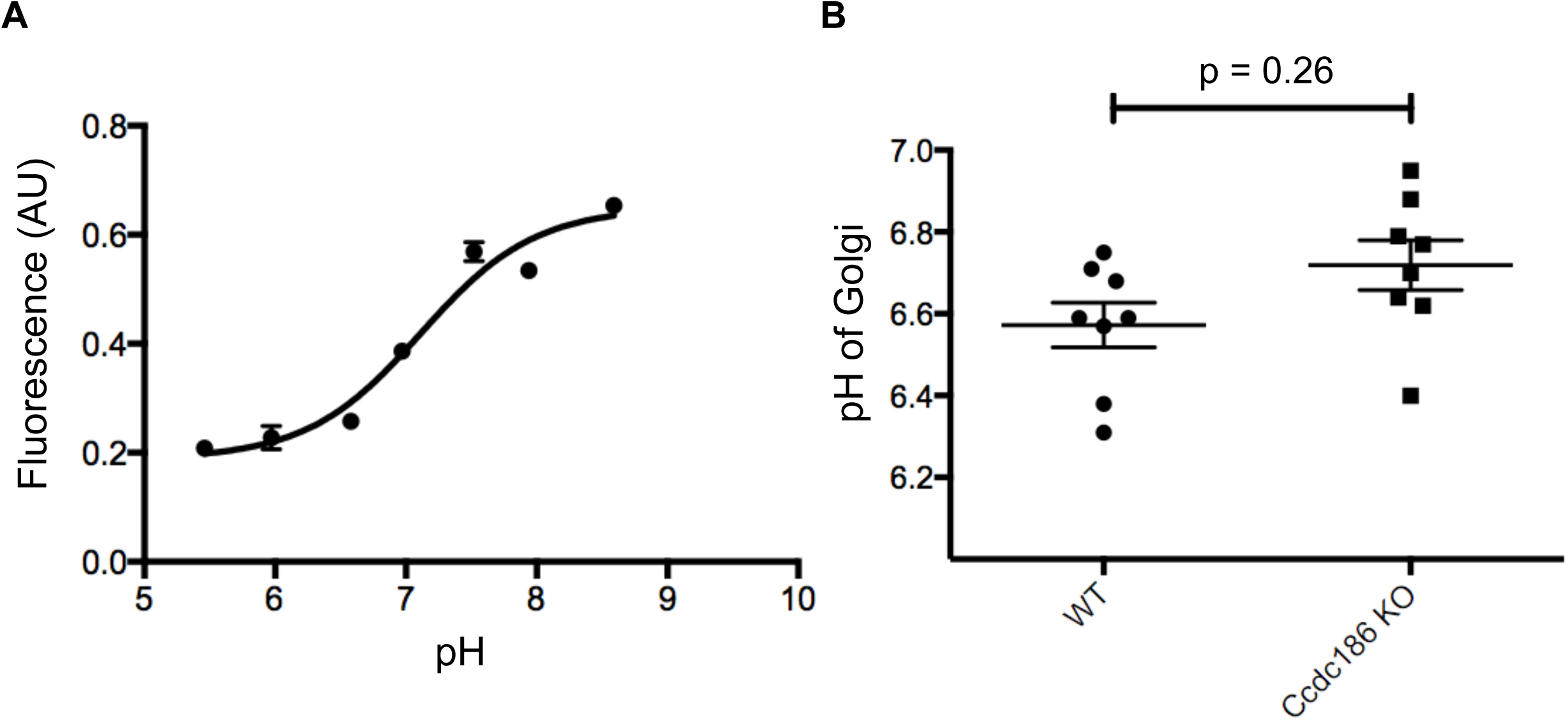
The acidity of the Golgi does not significantly change in *Ccdc186* KO cells. A. Example calibration curve for Golgi pH measurements based on measuring the fluorescence of TGN-targeted pHluorin in solutions of defined pH. For each biological replicate of WT and *Ccdc186* KO 832/13 cells tested for pH, an individual calibration curve such as the one shown was obtained from WT and *Ccdc186* KO cells grown in the same 96-well plate as the test samples and exposed to buffers of decreasing pH (8.5–5.5) in the presence of nigericin and monensin (see Materials and Methods). Each data point shows the mean of the fluorescent measurements (in arbitrary units (AU)) from cells in three different wells. Error bars=SEM. B. The late-Golgi compartment is not more acidic in *Ccdc186* KO 832/13 cells. The fluorescence of TGN-targeted pHluorin was measured in WT and *Ccdc186* KO cells. The absolute pH value of each sample was extrapolated from a paired calibration curve (as in A). The data show mean ± SEM; n=8 for WT and *Ccdc186* KO. The data shown for the WT are the same shown in Figure S4B of (Topalidou, Cattin-Ortolá, et al., 2019, manuscript submitted to *MBoC*) since these experiments were run in parallel with the same WT control.

**Figure S9.**
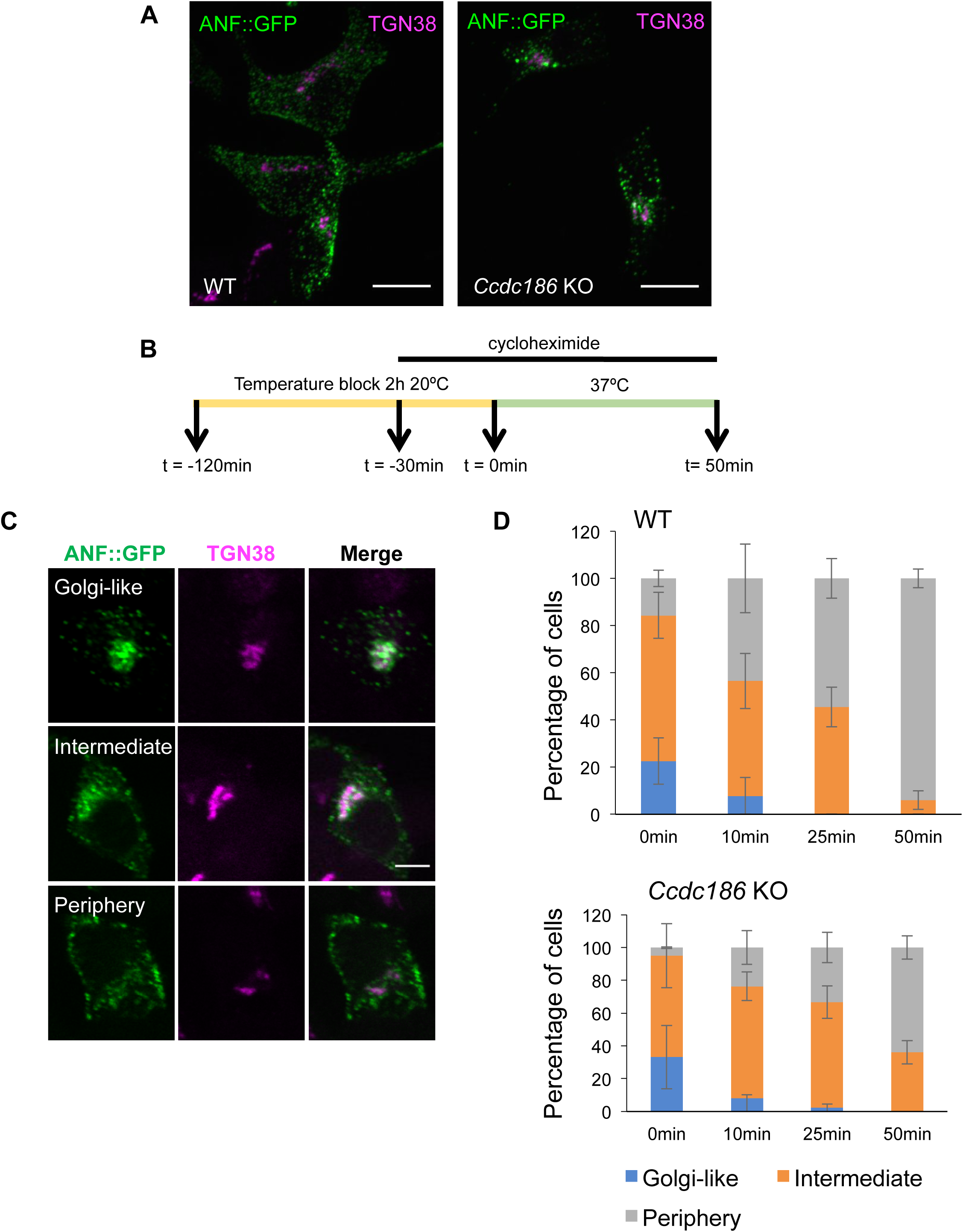
DCV cargo exit the TGN in *Ccdc186* KO cells. A. The exogenous DCV cargo ANF::GFP is mostly localized near the TGN in *Ccdc186* KO cells at steady state. Representative images of WT and *Ccdc186* KO 832/13 cells transfected with ANF::GFP and costained for GFP and TGN38. Maximum-intensity projections. Scale bars: 10 µm. B. Experimental set up. Cells were transiently transfected with ANF::GFP. Incubation at 20°C for 2 hours resulted in blocking ANF::GFP from exiting the TGN (pulse). 30 minutes before the end of the low temperature incubation, cycloheximide was added to inhibit synthesis of new ANF::GFP. Following the low temperature block, cells were incubated at 37°C (chase) before fixation and imaging. C. Representative images of the three cell categories used for qualitative assessment of TGN exit: those with most of the fluorescence concentrated at the TGN (Golgi-like), those where the TGN was still apparent but a large portion of the fluorescence was at the cell periphery (Intermediate), and those where the TGN was no longer apparent (Periphery). Single slices. Scale bar: 5 µm. D. ∼70 cells for each time point and genotype were imaged blindly. The data are plotted as percentage of each phenotype at each time point. The experiment was repeated three times with similar results. Error bars = SEM. The data shown for the WT are the same shown in Figure 4D of (Topalidou, Cattin-Ortolá, et al., 2019, manuscript submitted to *MBoC*) since these experiments were run in parallel with the same WT control.

**Figure S10.**
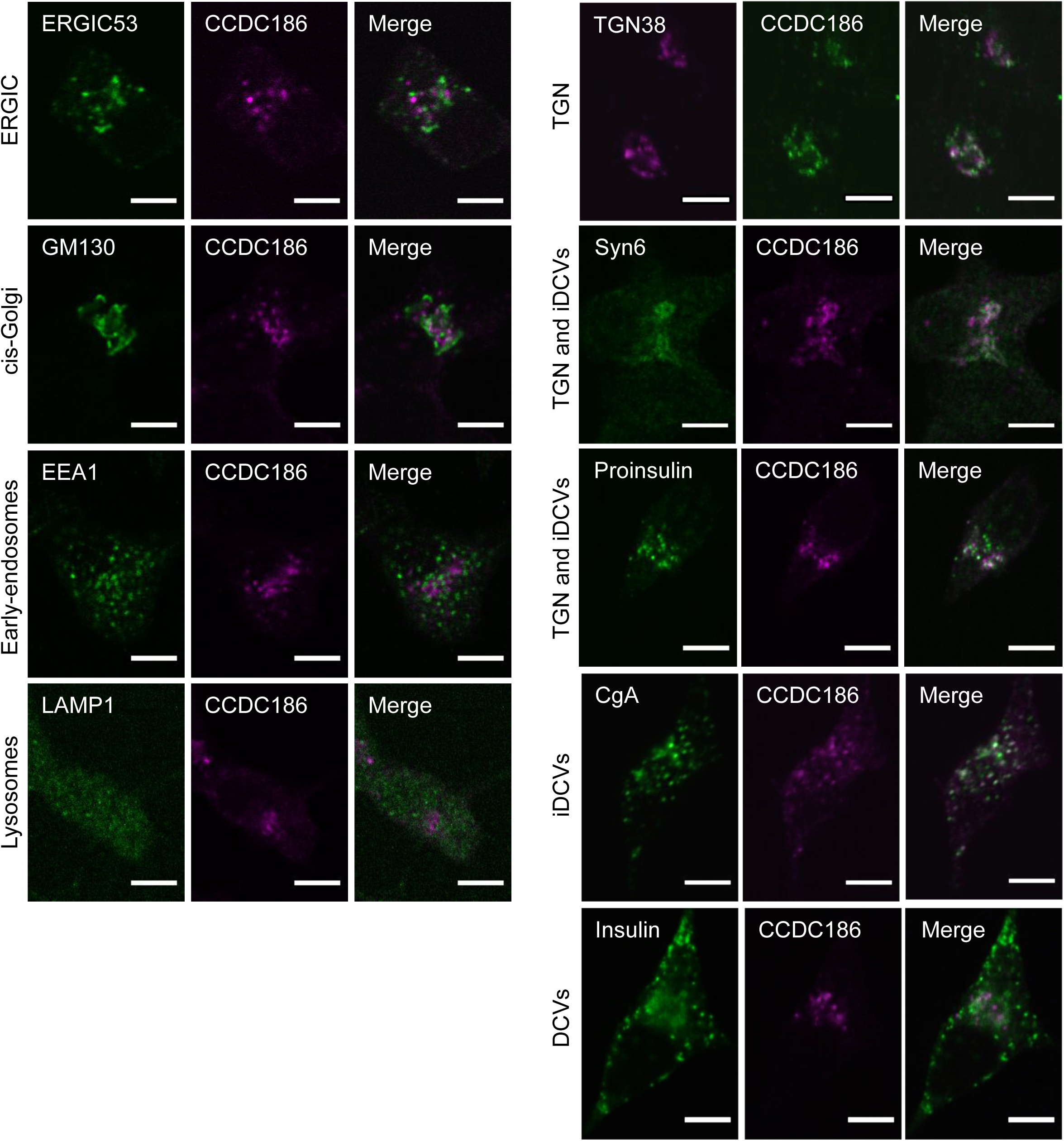
Endogenous CCDC186 localizes to a perinuclear area near the TGN and immature DCVs. Representative confocal images of 832/13 cells costained for endogenous CCDC186 and markers for different cell compartments: the ERGIC marker ERGIC53; the cis-Golgi marker GM130; the early-endosome marker EEA1; the lysosomal marker LAMP1; the TGN marker TGN38; the TGN/iDCV markers syntaxin 6 (Syn6), proinsulin, and chromogranin A (CgA); and the mature DCV marker insulin. Single slices. Scale bars: 5 µm.

**Figure S11.**
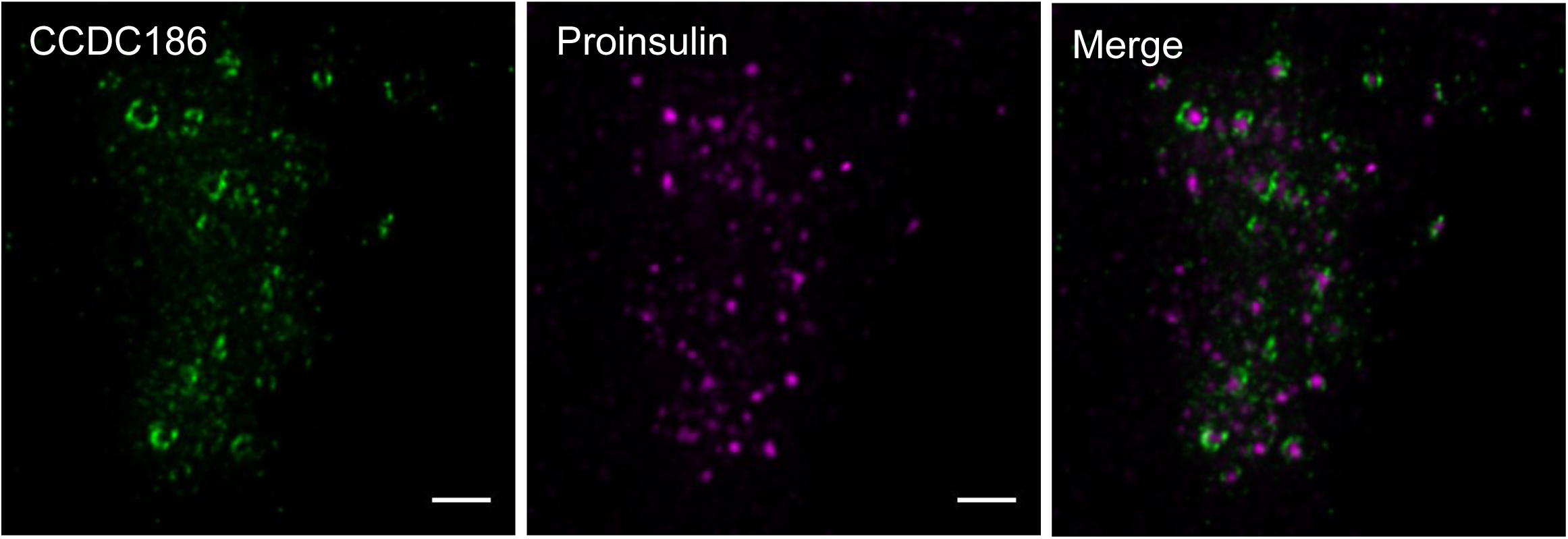
Endogenous CCDC186 localizes to ring-like structures around proinsulin. Representative stimulated emission depletion (STED) microscopy images of 832/13 cells costained for endogenous CCDC186 and endogenous proinsulin. Maximum-intensity projections. Scale bars: 2 µm.

**Figure S12.**
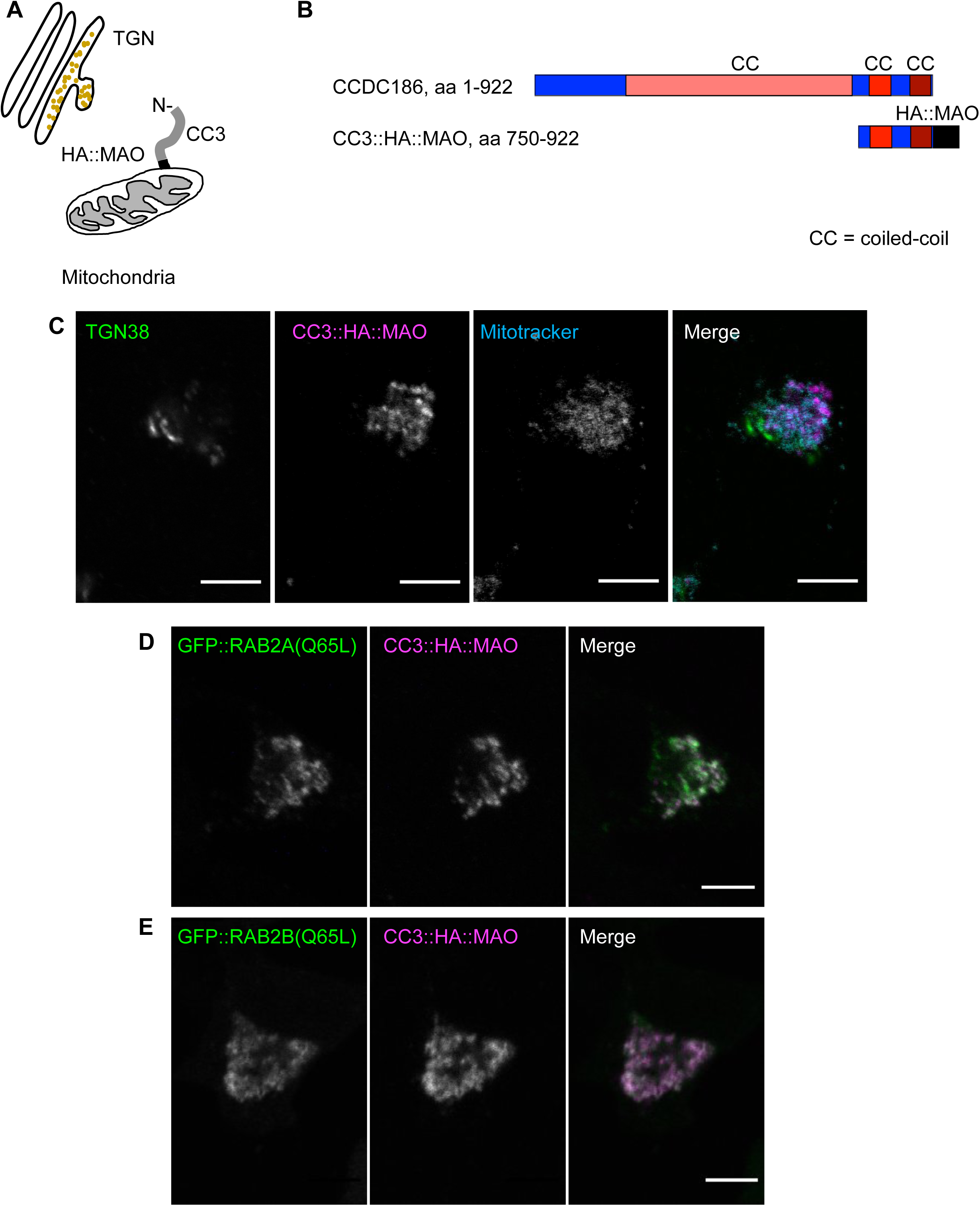
Mitochondria relocation of CCDC186 fragments. A. Schematic of the CCDC186 mitochondria relocation experiment. B. Domain structure of CCDC186 and CC3 fragment fused at their C-terminus to the transmembrane domain of monoamine oxidase (MAO, black). The domains marked with different shades of red are predicted coiled-coil domains (CC). C. The CC3 fragment fused to MAO is relocated to the mitochondria. Representative confocal images of 832/13 cells transiently transfected with CC3::HA::MAO, incubated with MitoTracker, and costained for HA and TGN38. The intensity and contrast of the image showing MitoTracker staining was adjusted to remove the high background. Of note, we observed that overexpression of CC3::HA::MAO caused clustering of mitochondria in a perinuclear region adjacent to the TGN marker TGN38. Single slices. Scale bars: 5 µm. The experiment was repeated twice. D,E. The CCDC186 fragment CC3 fused to MAO relocates the GTP-bound forms of RAB2A and RAB2B to mitochondria. Representative confocal images of 832/13 cells transiently cotransfected with CC3::HA::MAO and GFP::RAB2A(Q65L) or GFP::RAB2B(Q65L), and costained for HA and GFP. Single slices. Scale bars: 5 µm. The experiment was repeated twice.

**Figure S13.**
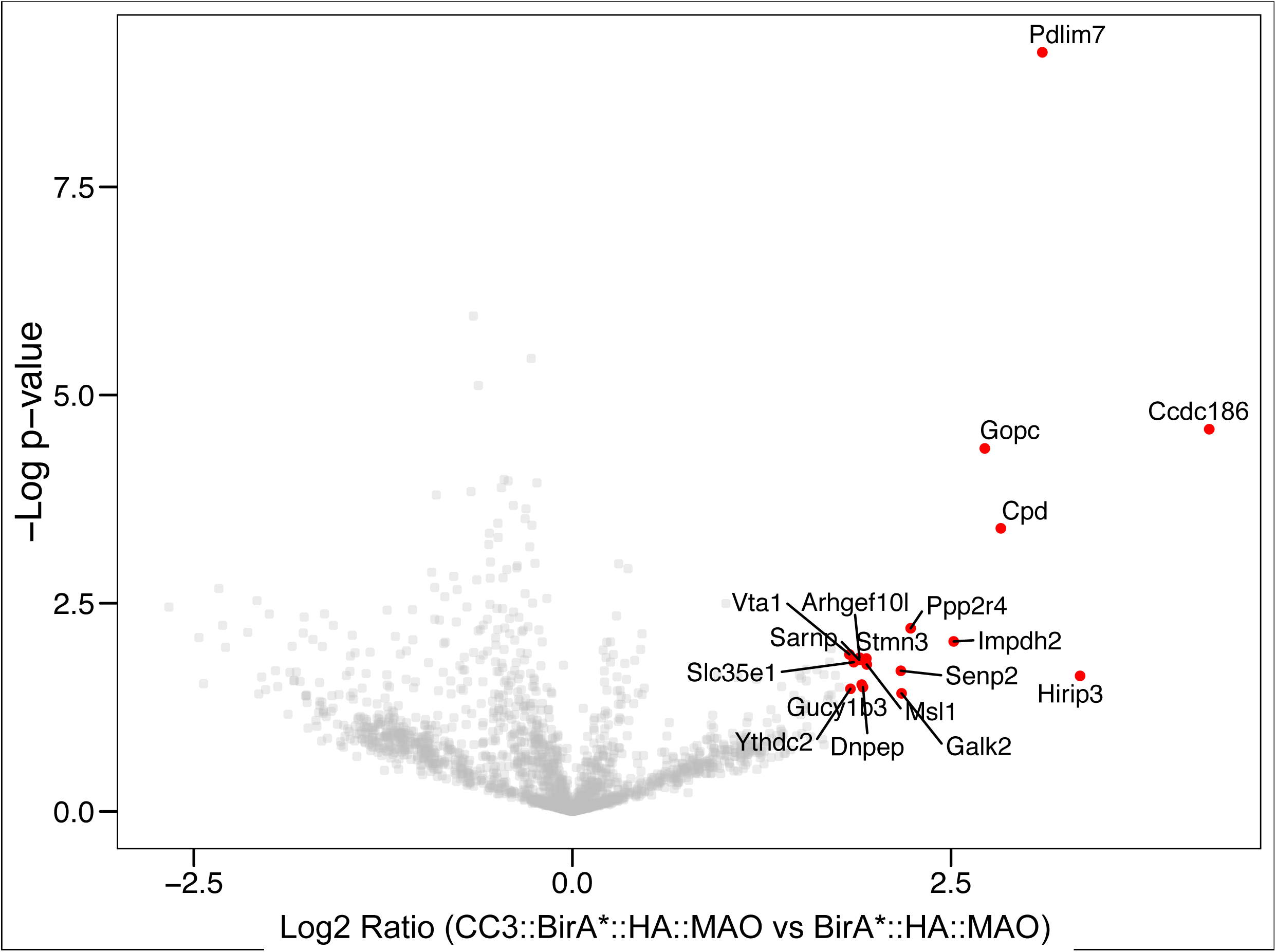
Volcano plot representation of the CC3 BioID experiment. Volcano plot showing the Student’s t-test p-value for each protein as a measure of that protein’s enrichment in CC3::BirA*::HA::MAO streptavidin pulldowns compared to the negative control BirA*::HA::MAO. Lysates from cells overexpressing CC3::BirA*::HA::MAO and BirA*::HA::MAO were subjected to affinity enrichment using streptavidin. Proteins with a Log2 ratio (CC3::BirA*::HA::MAO / BirA*::HA::MAO) larger than 1.8 are labeled.

**Figure S14.**
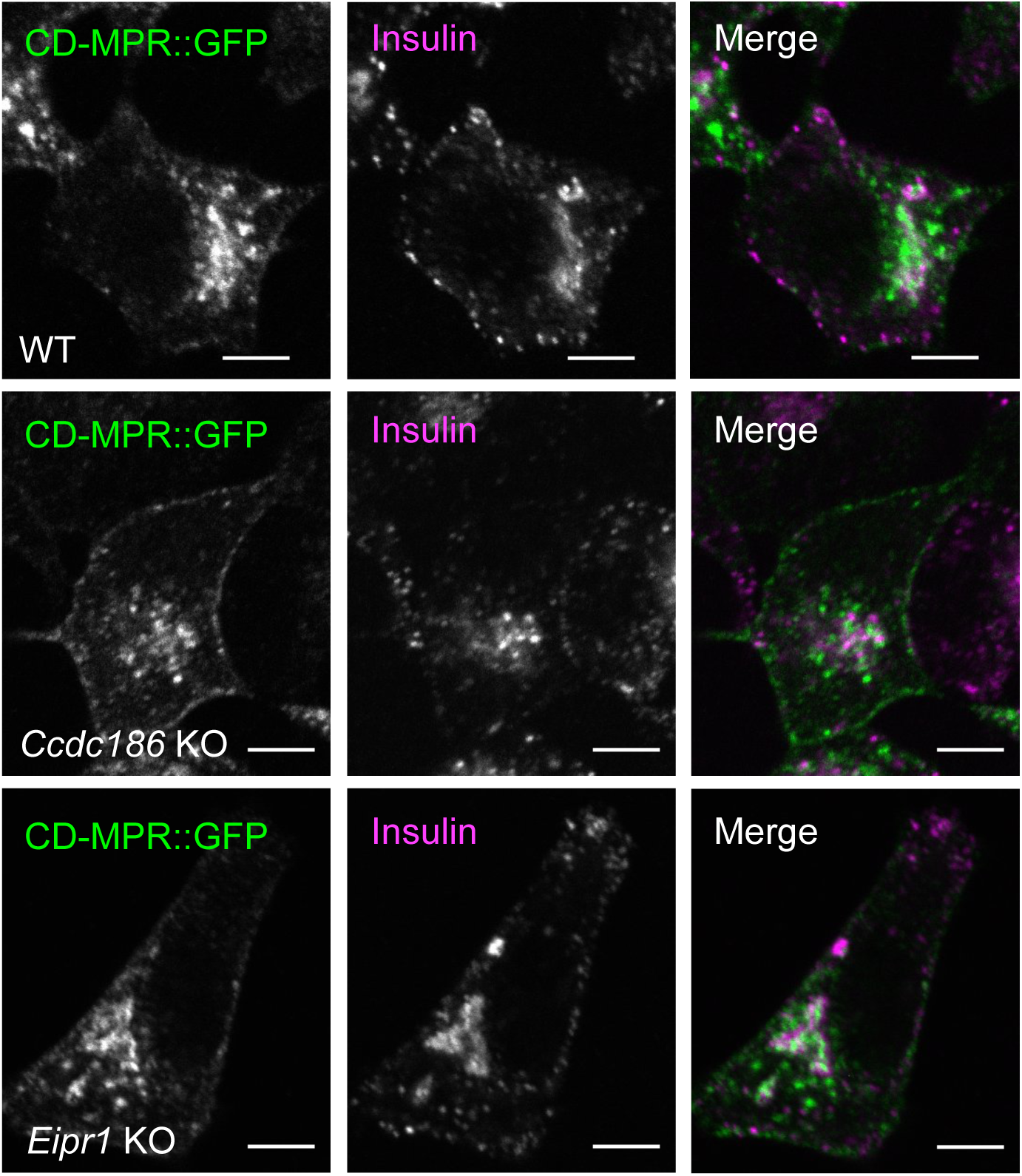
CCDC186 and EIPR1 are not required for CD-MPR trafficking. Representative confocal images of WT, *Ccdc186* KO, and *Eipr1* KO 832/13 cells transiently transfected with CD-MPR::GFP and costained for insulin and GFP. Single slices. Scale bars: 5 µm. The experiment was repeated twice.

**Table S1.**
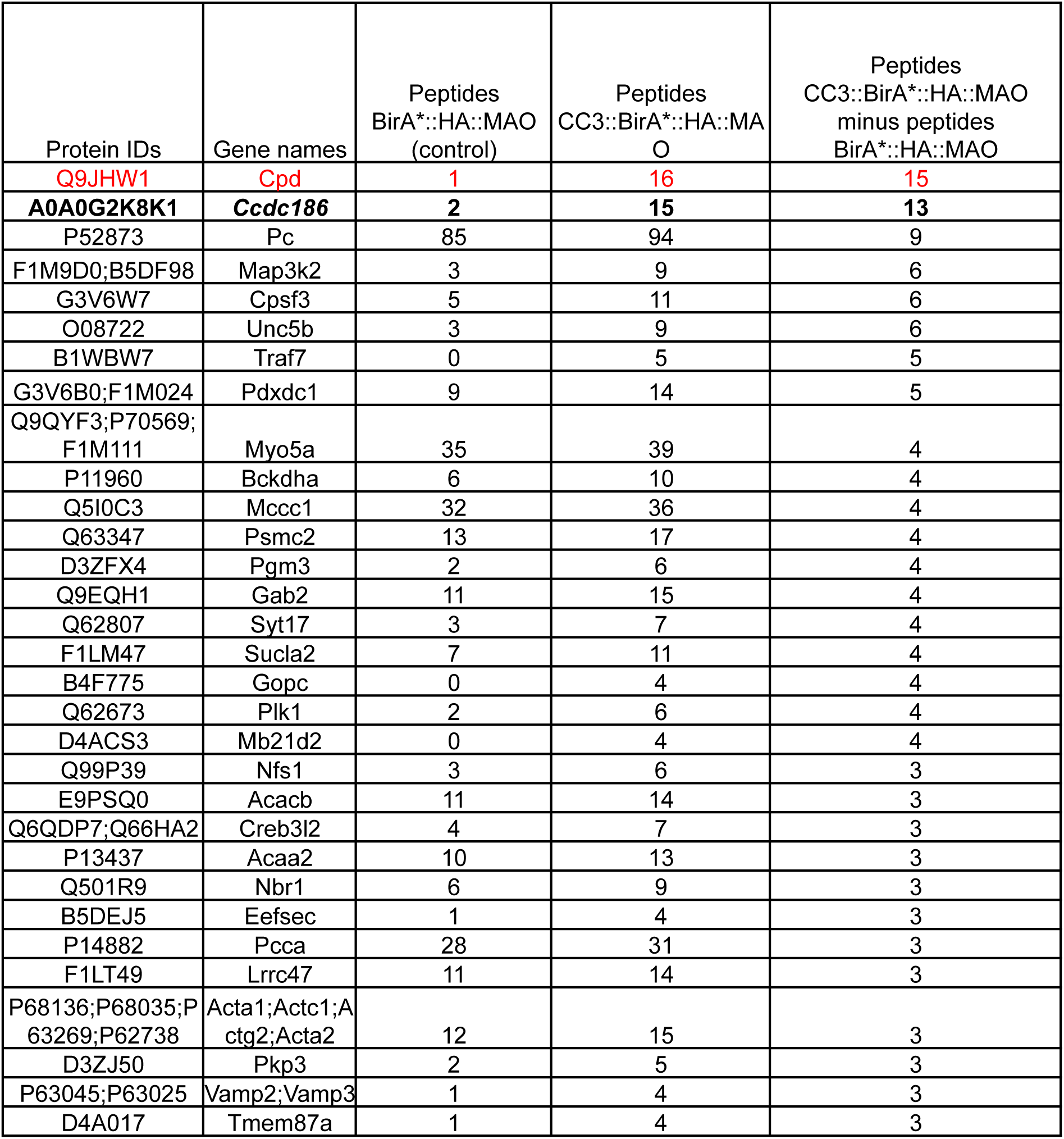
Top hits from CC3 BioID experiment. Unique peptide count organized in the descending order of the difference in number of unique peptides from CC3::BirA*::HA::MAO and the negative control BirA*::HA::MAO. The table shows the hits where the difference was 3 or greater. These data are from the BioID experiment that showed the largest amount of enrichment for CPD. For raw data, see Table S6.

**Table S2.**
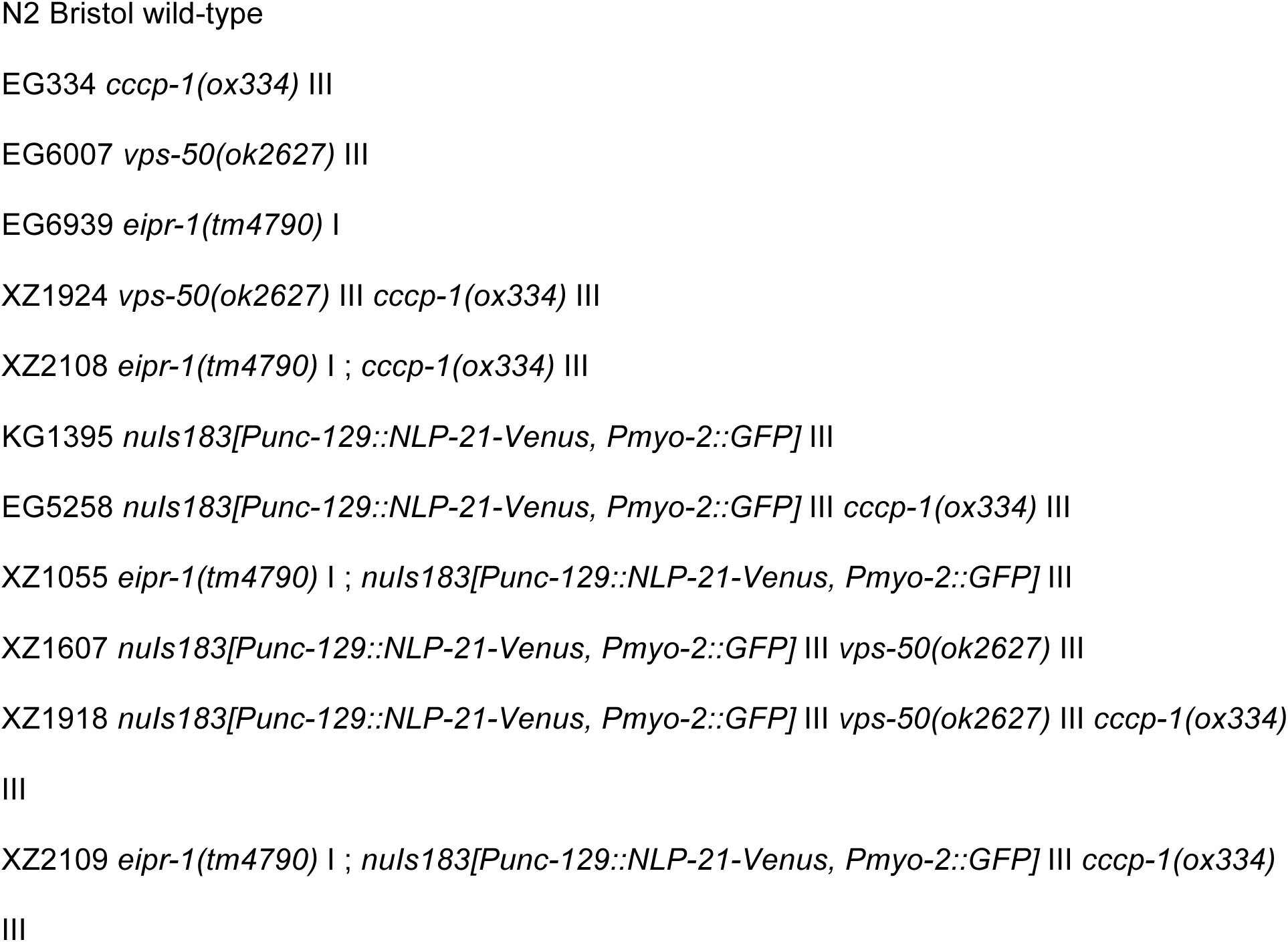
C. elegans strain list.

**Table S3.**
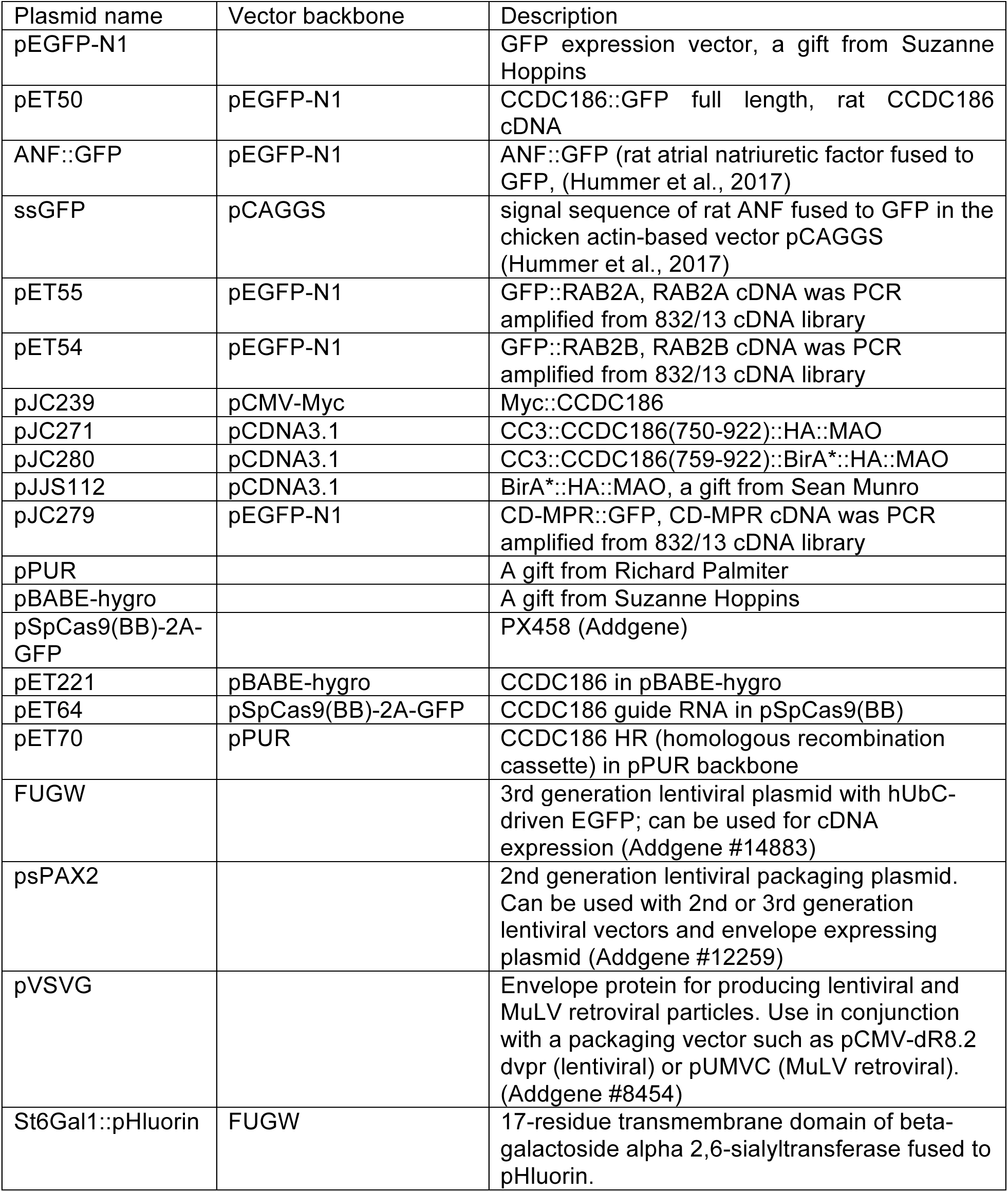
Plasmid list.

**Table S4.**
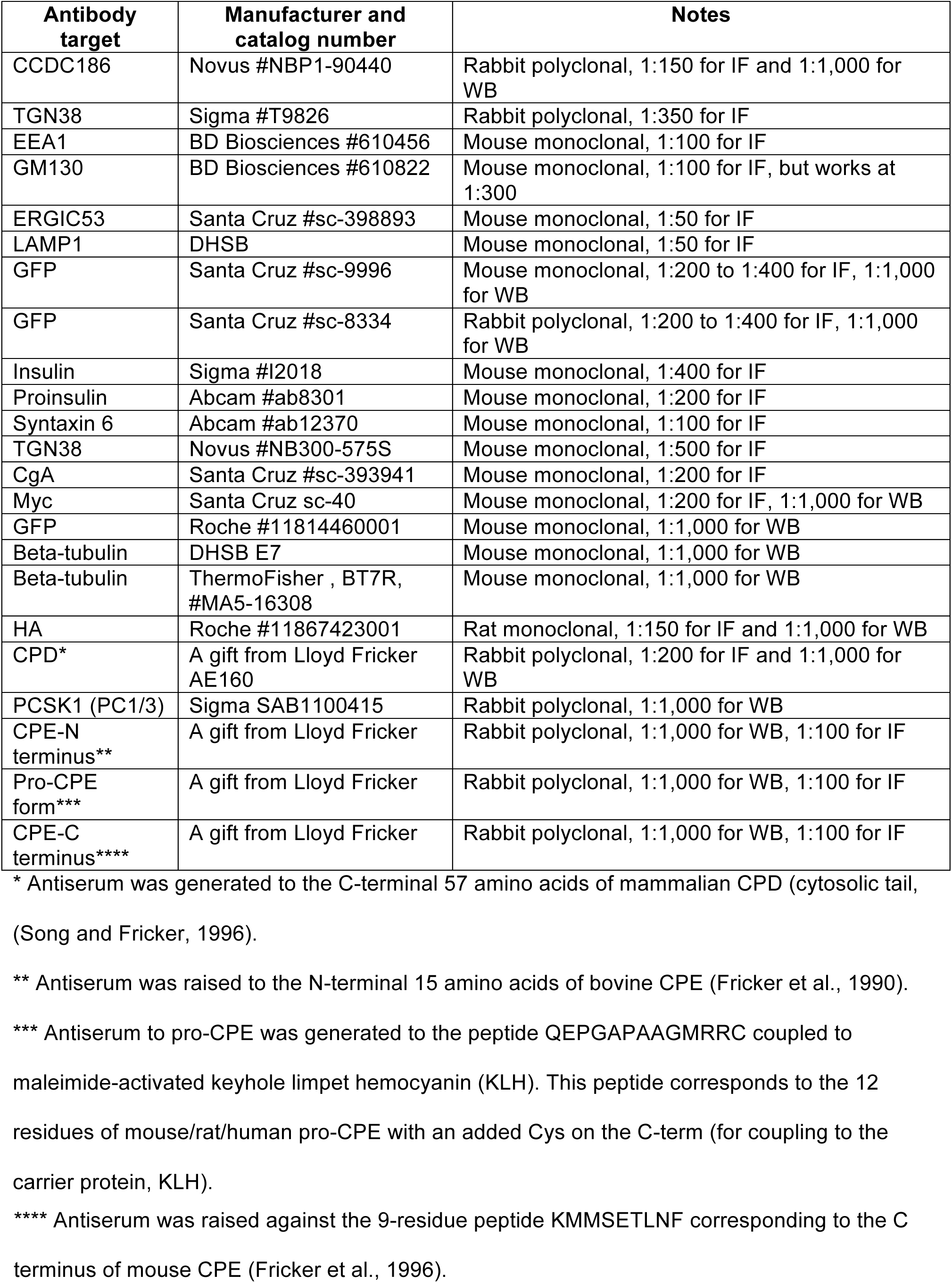
Primary antibody list.

**Table S5.**
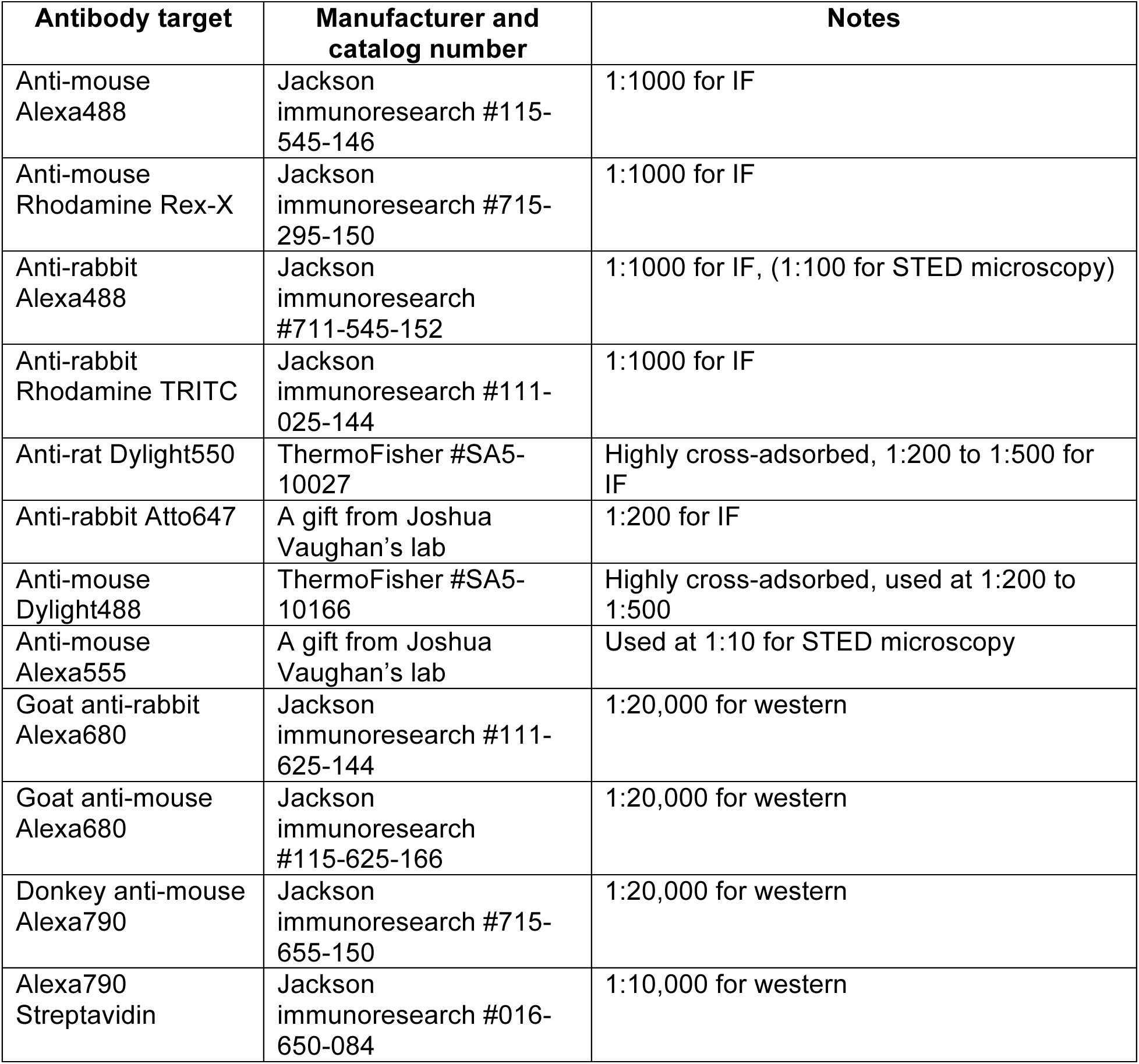
Secondary antibody list.

**Table S6. Raw mass-spec data for CC3 BioID experiment.**

The table contains all the data for the experiment shown in Table S1.

**Table S7. LFQ intensities for CC3 BioID experiment.**

The table contains Z-score normalized log2 LFQ intensities from MaxQuant output, Student’s t-test p-values and ratios (CC3::BirA*::HA::MAO vs BirA*::HA::MAO). For calculation of p-values and ratios, see Materials and Methods.

