## Supplementary figures and tables for "CCDC186 controls dense-core vesicle cargo sorting by exit"

### Figure S1

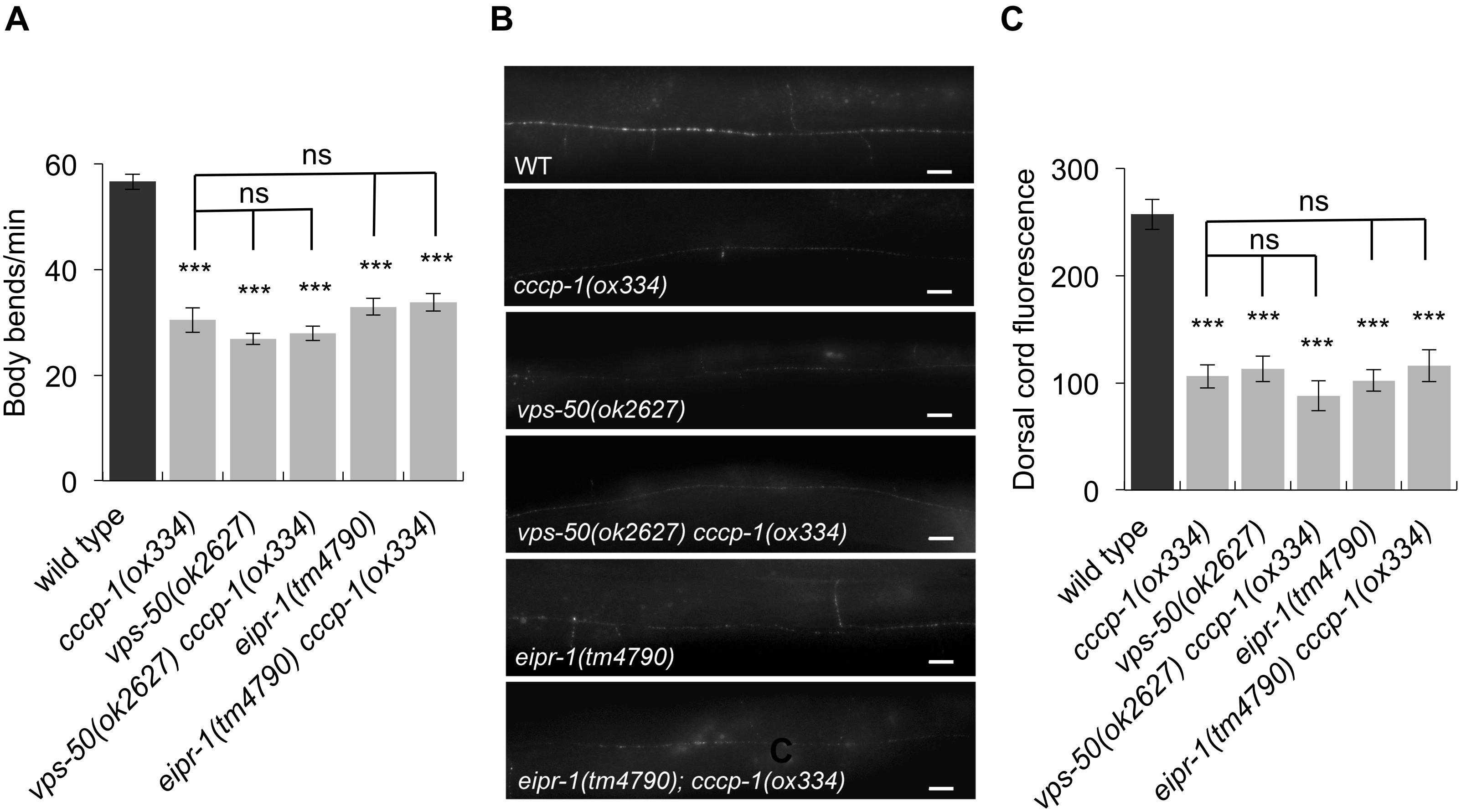
A
B
C
Body bends/min
60
40
20
0
ns
ns
***
wild type
cccp-1(ox334)
vps-50(ok2627)
vps-50(ok2627) cccp-1(ox334)
eipr-1(tm4790)
eipr-1(tm4790) cccp-1(ox334)
WT
cccp-1(ox334)
vps-50(ok2627)
vps-50(ok2627) cccp-1(ox334)
eipr-1(tm4790)
eipr-1(tm4790); cccp-1(ox334)
Dorsal cord fluorescence
300
200
100
0

### Figure S2

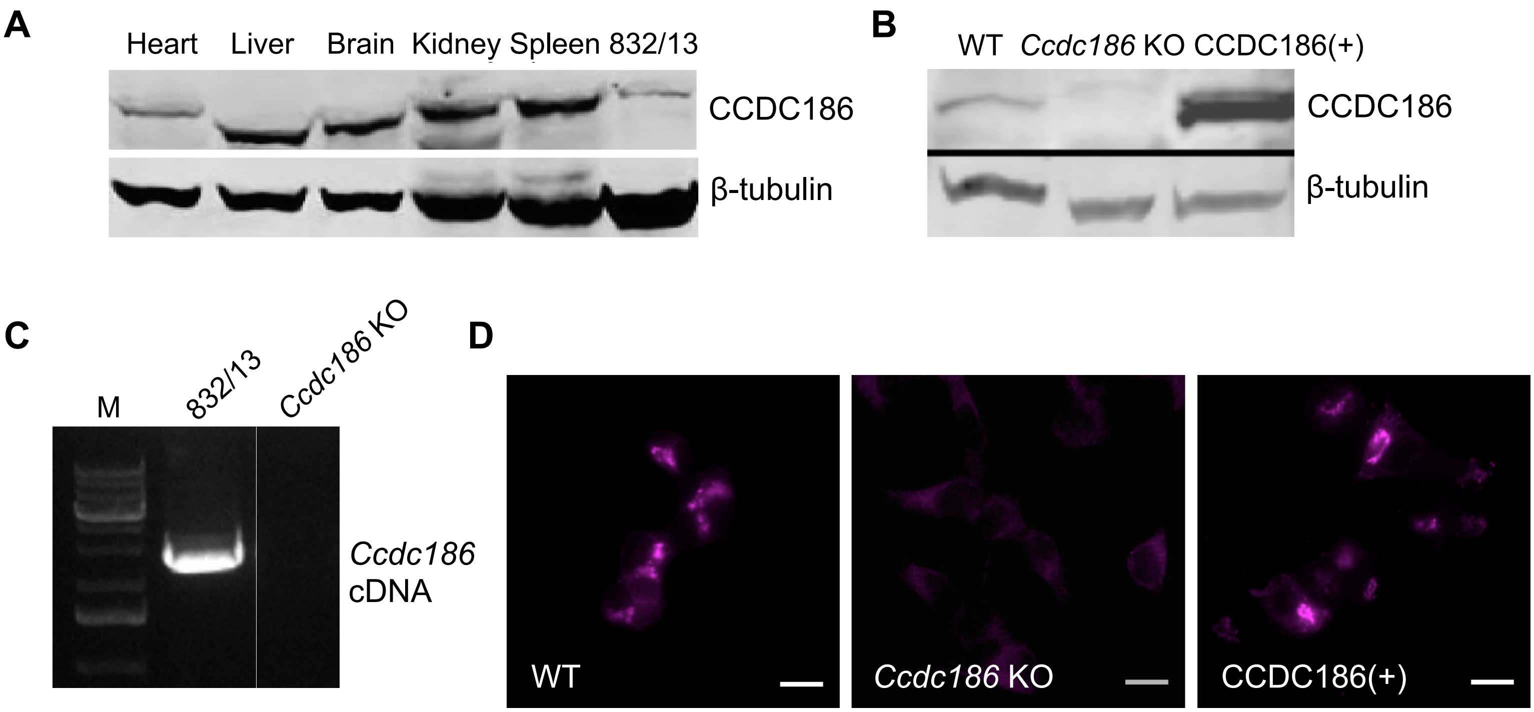
A
Heart
Liver
Brain
Kidney
Spleen
832/13
CCDC186
β-tubulin
B
WT
Ccdc186 KO
CCDC186(+)
CCDC186
β-tubulin
C
M
832/13
Ccdc186 KO
Ccdc186 cDNA
D
WT
Ccdc186 KO
CCDC186(+)

### Figure S3

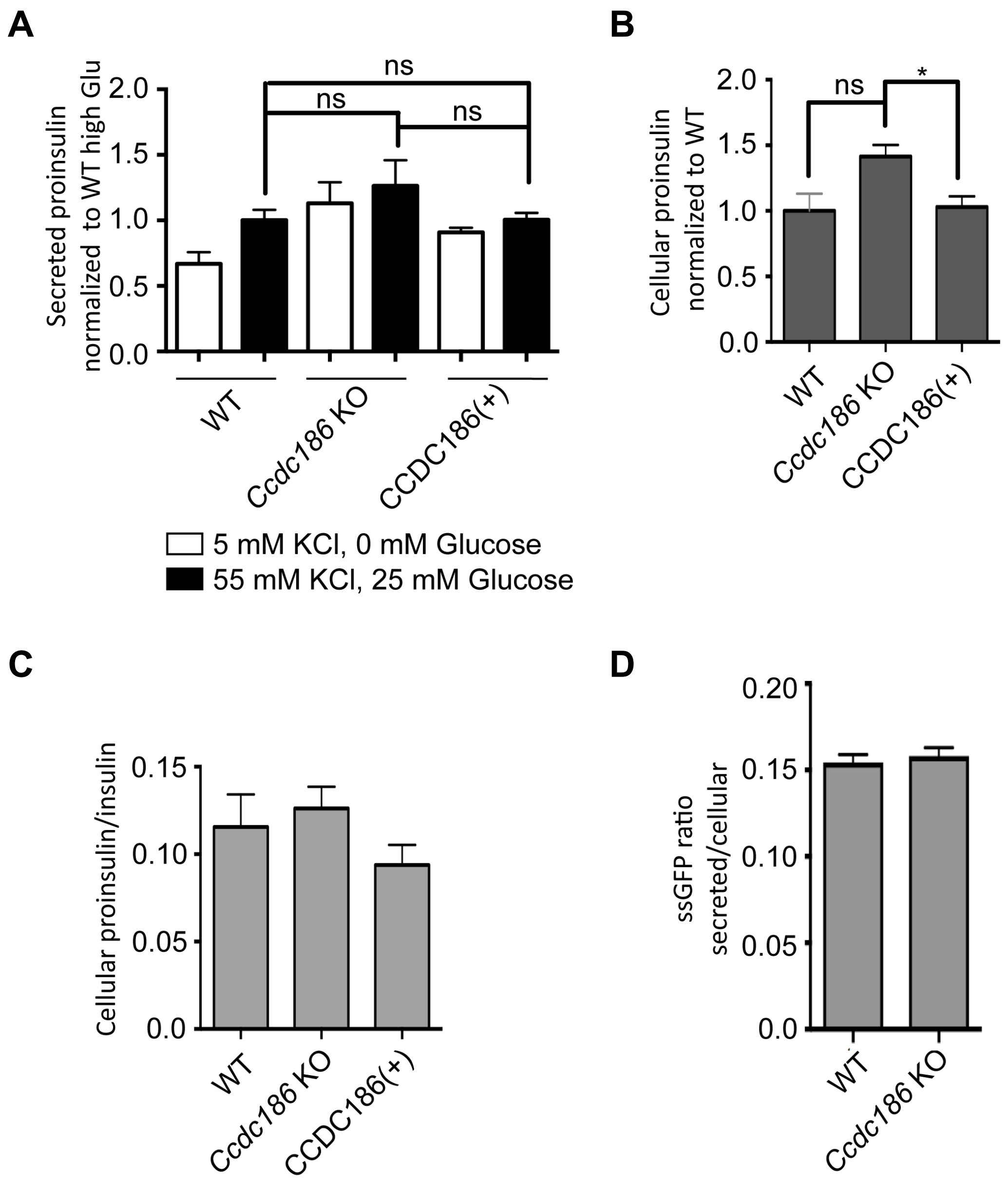
A
B
C
D
Secreted proinsulin normalized to WT high Glu
2.0
1.5
1.0
0.5
0.0
ns
ns
ns
WT
Ccdc186 KO
CCDC186(+)
5 mM KCl, 0 mM Glucose
55 mM KCl, 25 mM Glucose
Cellular proinsulin normalized to WT
ns
*
Cellular proinsulin/insulin
0.15
0.10
0.05
0.0
ssGFP ratio secreted/cellular
0.20

**Figure S4**

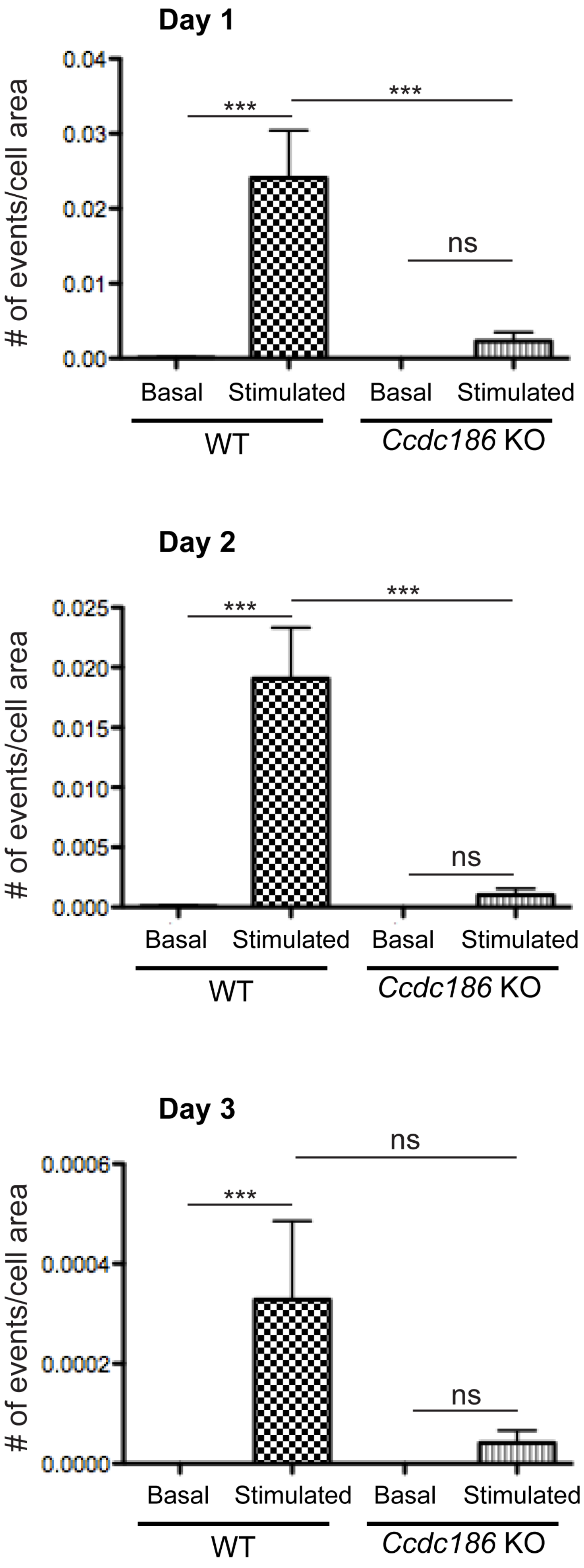

Day 1
### of events/cell area
0.04
0.03
0.02
0.01
0.00
***
***
ns
Basal
Stimulated
Basal
Stimulated
WT
Ccdc186 KO
Day 2
### of events/cell area
0.025
0.020
0.015
0.010
0.005
0.000
***
***
ns
Basal
Stimulated
Basal
Stimulated
WT
Ccdc186 KO
Day 3
### of events/cell area
0.0006
0.0004
0.0002
0.0000
ns
***
ns
Basal
Stimulated
Basal
Stimulated
WT
Ccdc186 KO

### Figure S5

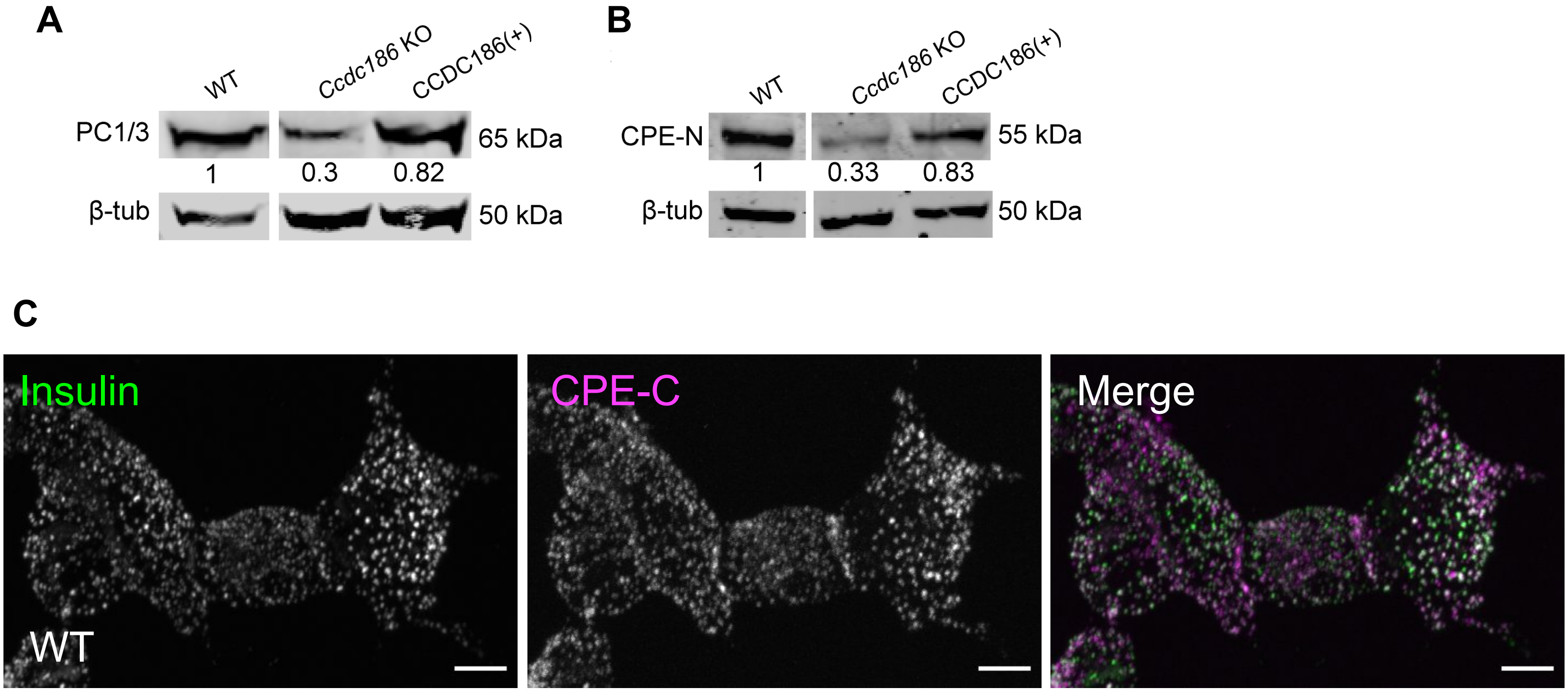
A
WT
Ccdc186 KO
CCDC186(+)
PC1/3
65 kDa
1
0.3
0.82
β-tub
50 kDa
B
WT
Ccdc186 KO
CCDC186(+)
CPE-N
55 kDa
1
0.33
0.83
β-tub
50 kDa
C
Insulin
CPE-C
Merge
WT

### Figure S6

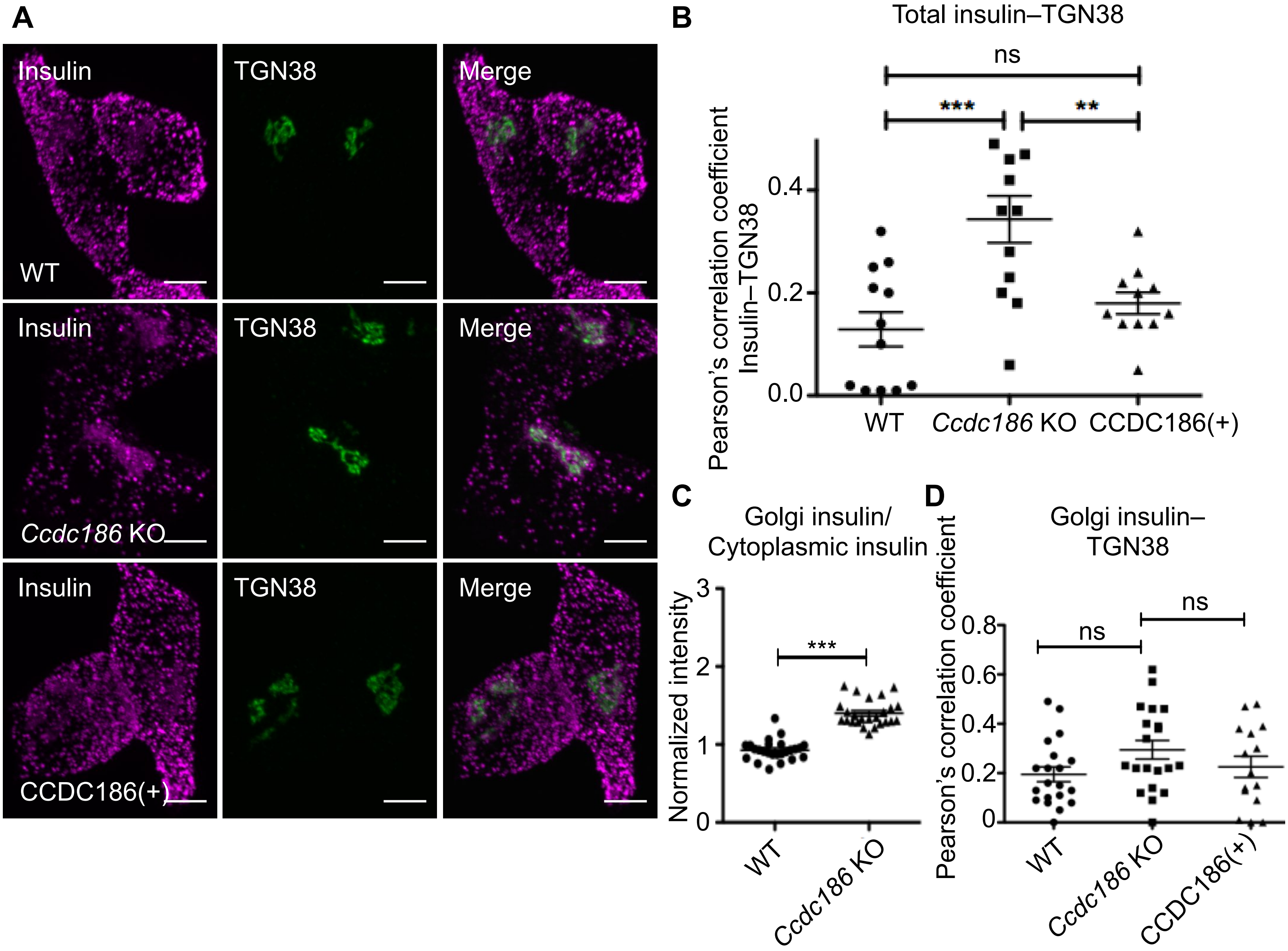
A
Insulin
TGN38
Merge
WT
Insulin
TGN38
Merge
Ccdc186 KO
Insulin
TGN38
Merge
CCDC186(+)
B
Total insulin–TGN38
ns
***
**
Pearson's correlation coefficient
Insulin–TGN38
0.4
0.2
0.0
WT
Ccdc186 KO
CCDC186(+)
C
Golgi insulin/
Cytoplasmic insulin
Normalized intensity
3
2
1
0
***
WT
Ccdc186 KO
D
Golgi insulin–
TGN38
Pearson's correlation coefficient
0.8
0.6
0.4
0.2
0
ns
ns
WT
Ccdc186 KO
CCDC186(+)

#### Figure S7

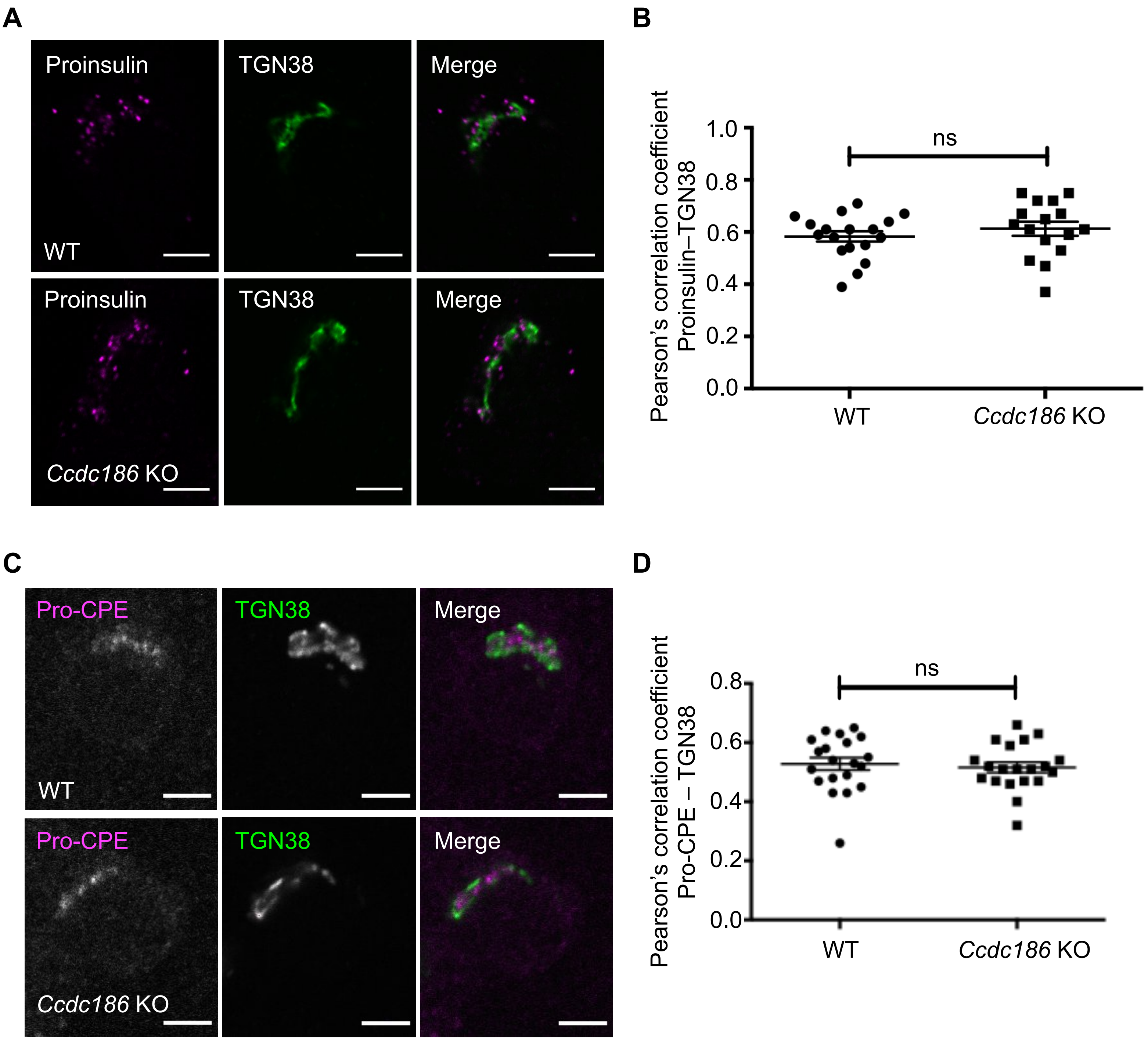

A
Proinsulin
TGN38
Merge
WT
Proinsulin
TGN38
Merge
Ccdc186 KO
B
Pearson's correlation coefficient
Proinsulin–TGN38
1.0
0.8
0.6
0.4
0.2
0.0
ns
WT
Ccdc186 KO
C
Pro-CPE
TGN38
Merge
WT
Pro-CPE
TGN38
Merge
Ccdc186 KO
D
Pearson's correlation coefficient
Pro-CPE – TGN38
0.8
0.6
0.4
0.2
0.0
ns
WT
Ccdc186 KO

**Figure S8**

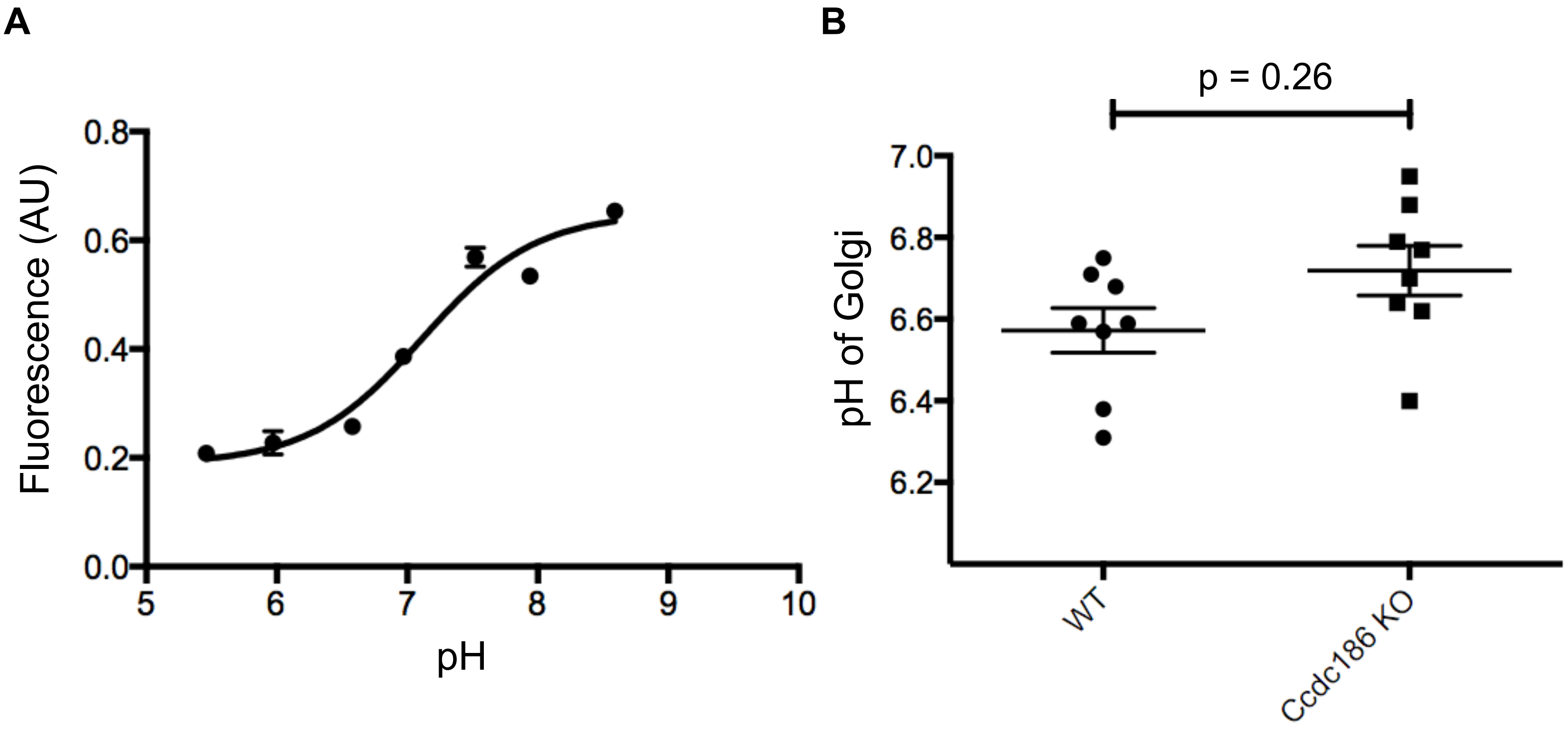

A
B
Fluorescence (AU)
0.0
0.2
0.4
0.6
0.8
5
6
7
8
9
10
pH
p = 0.26
pH of Golgi
6.2
6.4
6.6
6.8
7.0
WT
Ccdc186 KO

### Figure S9

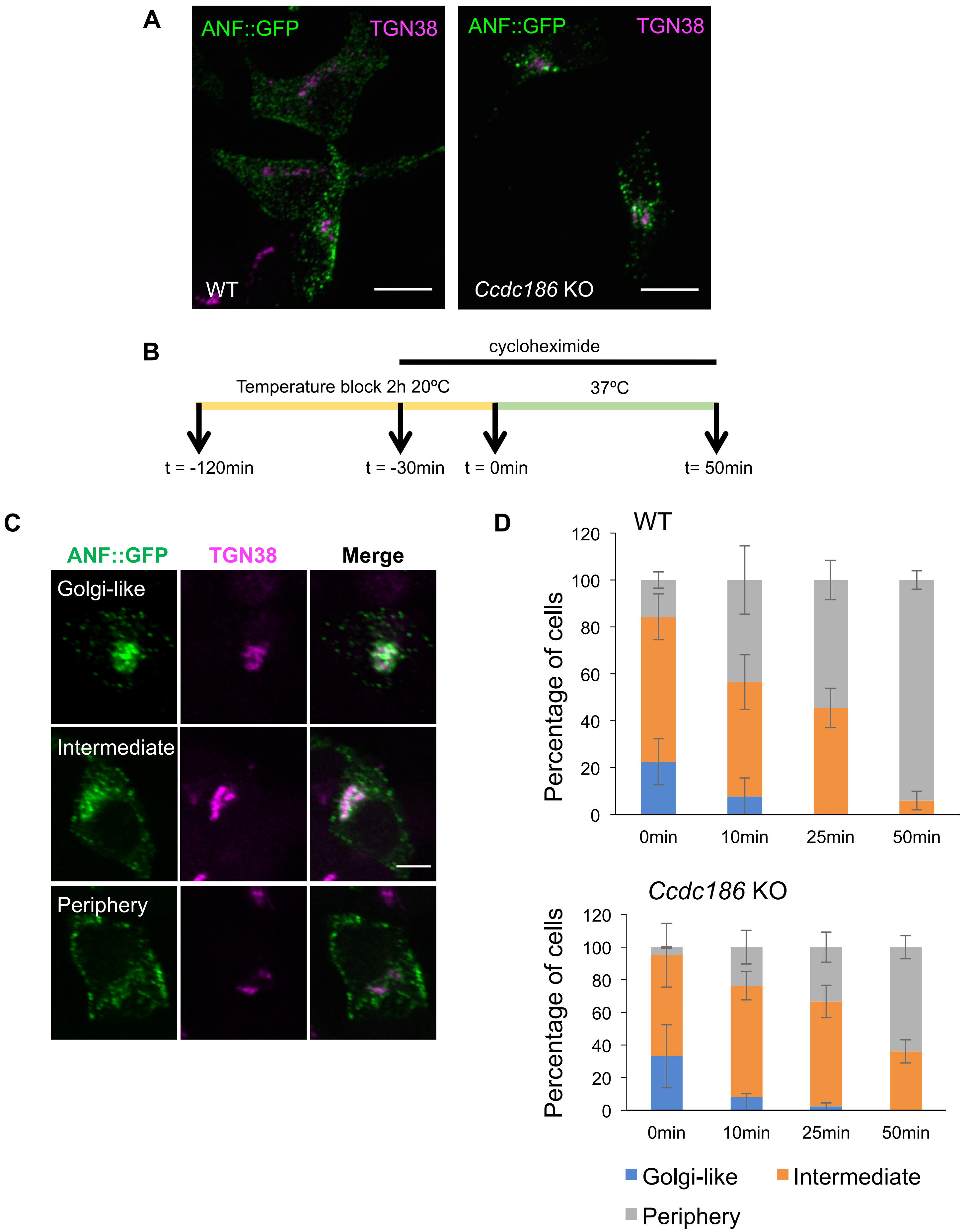
A
ANF::GFP
TGN38
WT
Ccdc186 KO
B
cycloheximide
Temperature block 2h 20ºC
37ºC
t = -120min
t = -30min
t = 0min
t= 50min
C
ANF::GFP
TGN38
Merge
Golgi-like
Intermediate
Periphery
D
WT
Percentage of cells
120
100
80
60
40
20
0
0min
10min
25min
50min
Ccdc186 KO
Golgi-like
Intermediate
Periphery

### Figure S10

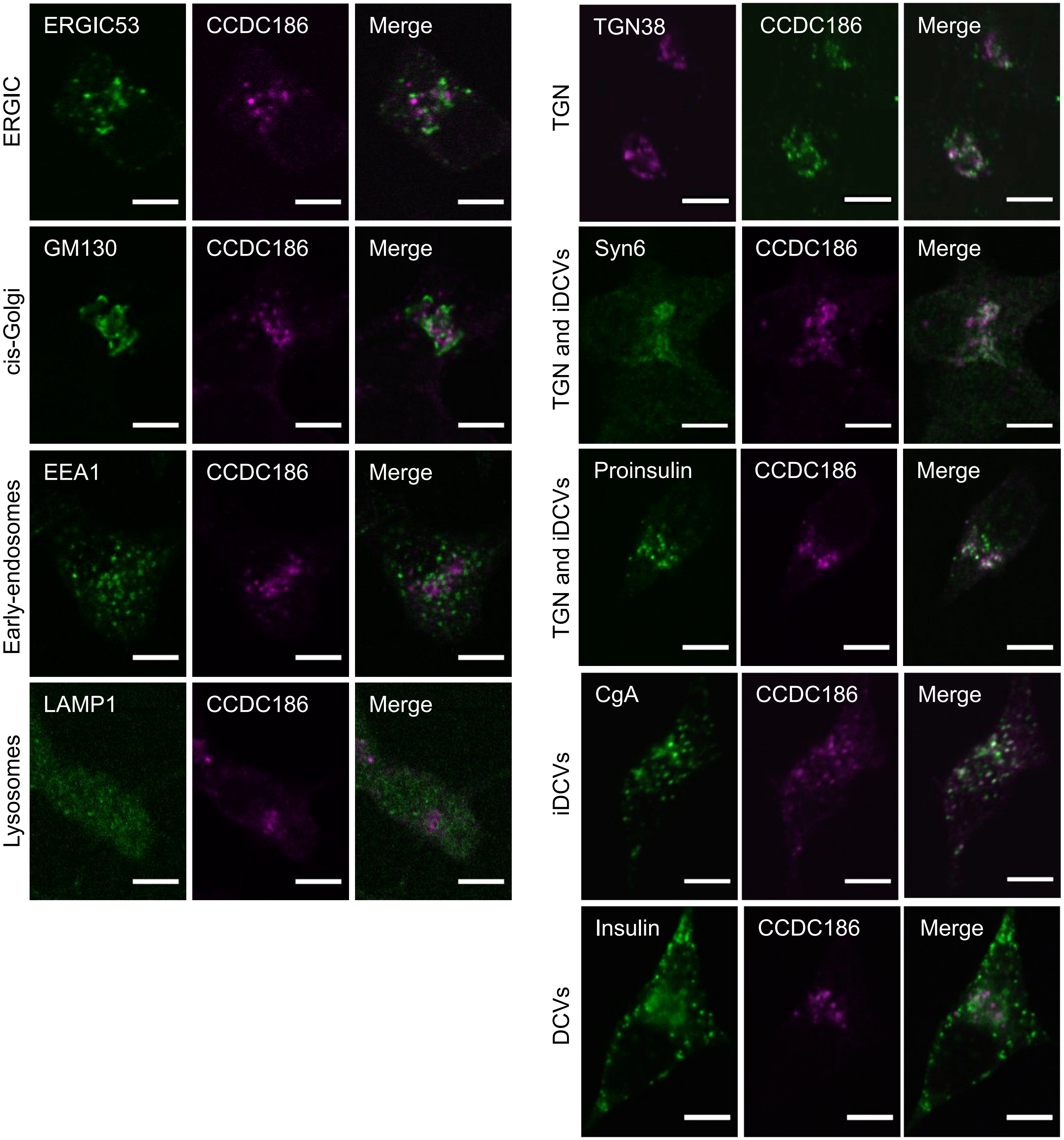

ERGIC
ERGIC53
CCDC186
Merge
cis-Golgi
GM130
CCDC186
Merge
Early-endosomes
EEA1
CCDC186
Merge
Lysosomes
LAMP1
CCDC186
Merge
TGN
TGN38
CCDC186
Merge
TGN and iDCVs
Syn6
CCDC186
Merge
TGN and iDCVs
Proinsulin
CCDC186
Merge
iDCVs
CgA
CCDC186
Merge
DCVs
Insulin
CCDC186
Merge

### Figure S11

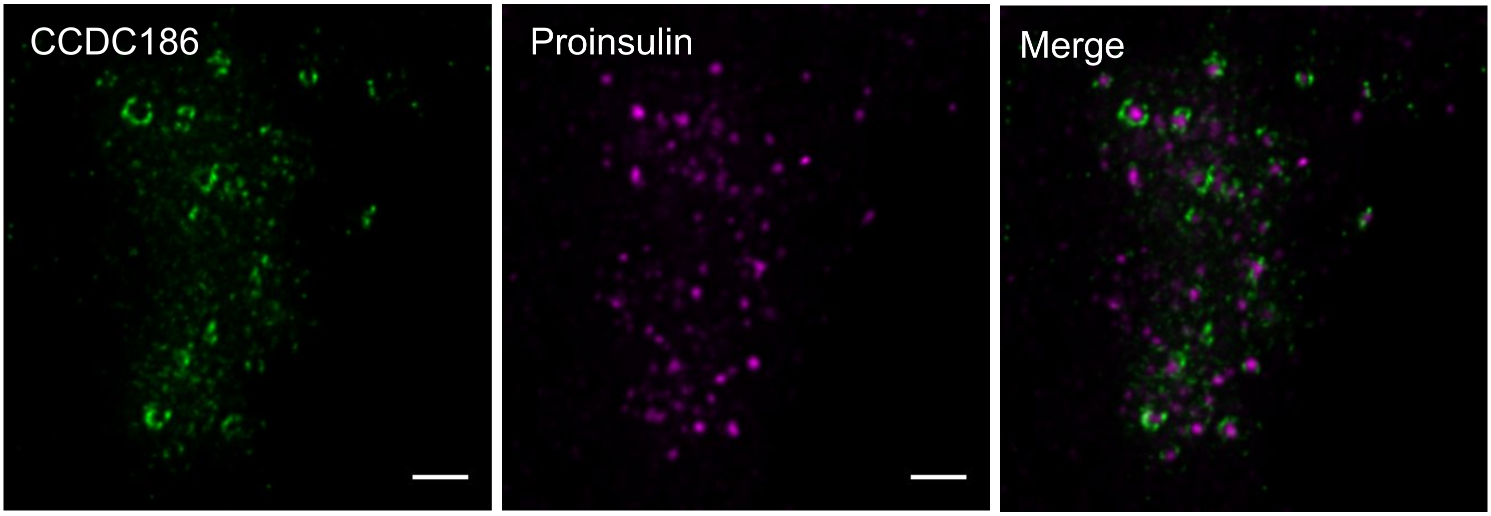

### Figure S12

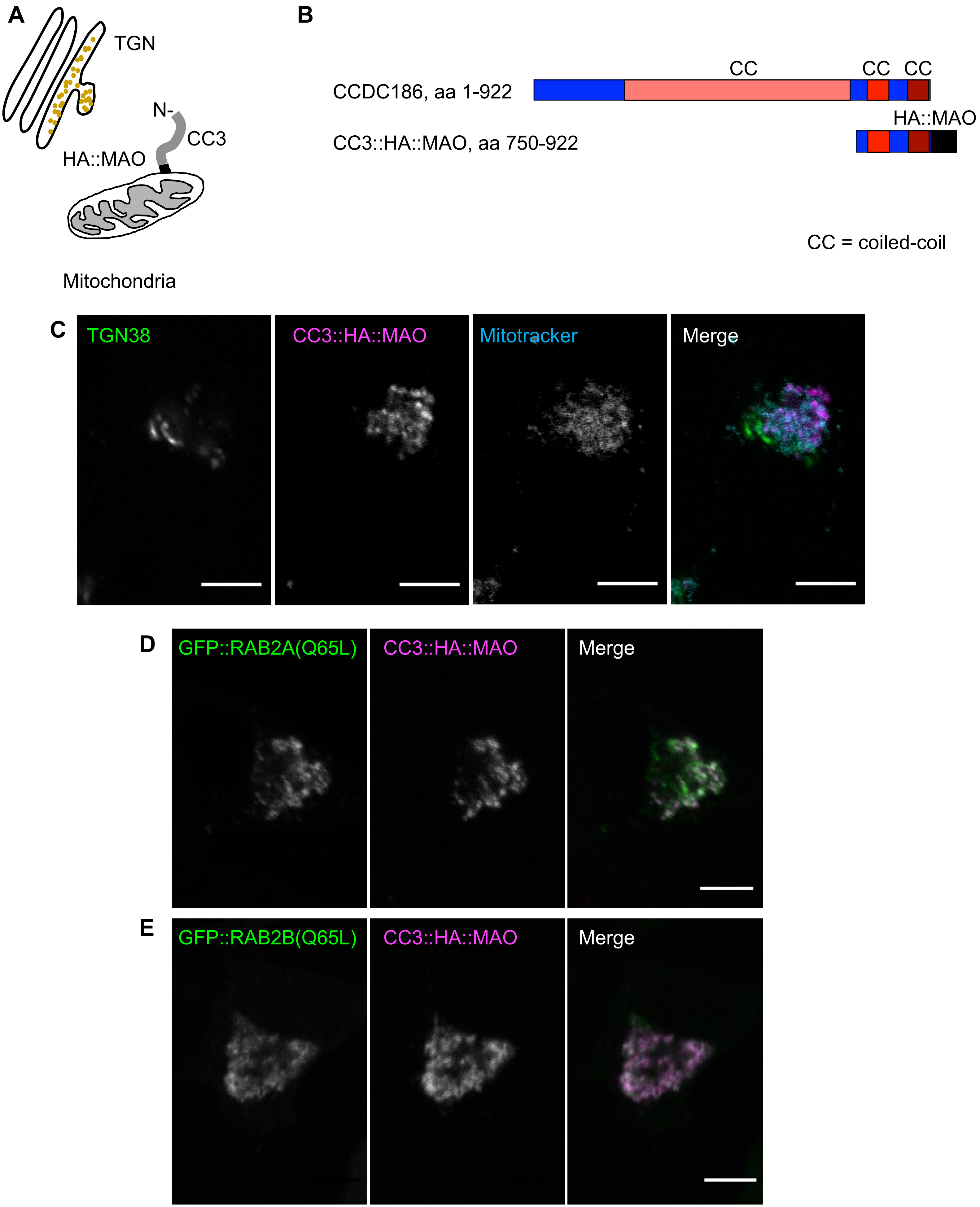

A
TGN
N-
CC3
HA::MAO
Mitochondria
B
CC
CC
CC
CCDC186, aa 1-922
HA::MAO
CC3::HA::MAO, aa 750-922
CC = coiled-coil
C
TGN38
CC3::HA::MAO
Mitotracker
Merge
D
GFP::RAB2A(Q65L)
CC3::HA::MAO
Merge
E
GFP::RAB2B(Q65L)
CC3::HA::MAO
Merge

**Figure S13**

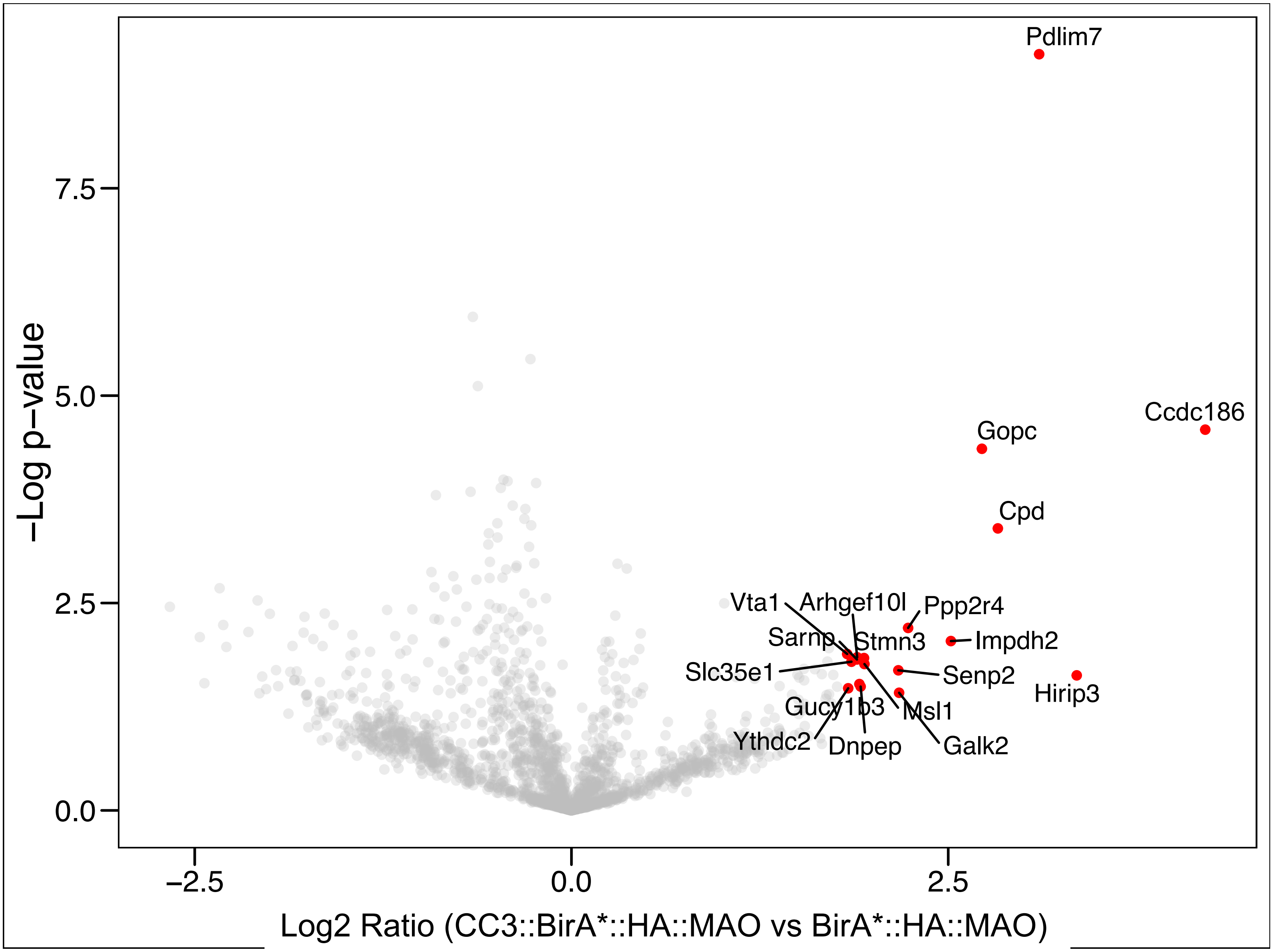
Pdlim7
Ccdc186
Gopc
Cpd
Vta1
Arhgef10l
Ppp2r4
Sarnp
Stmn3
Impdh2
Slc35e1
Senp2
Hirip3
Gucy1b3
Msl1
Ythdc2
Dnpep
Galk2
−Log p−value
7.5
5.0
2.5
0.0
−2.5
0.0
2.5
Log2 Ratio (CC3::BirA*::HA::MAO vs BirA*::HA::MAO)

### Figure S14

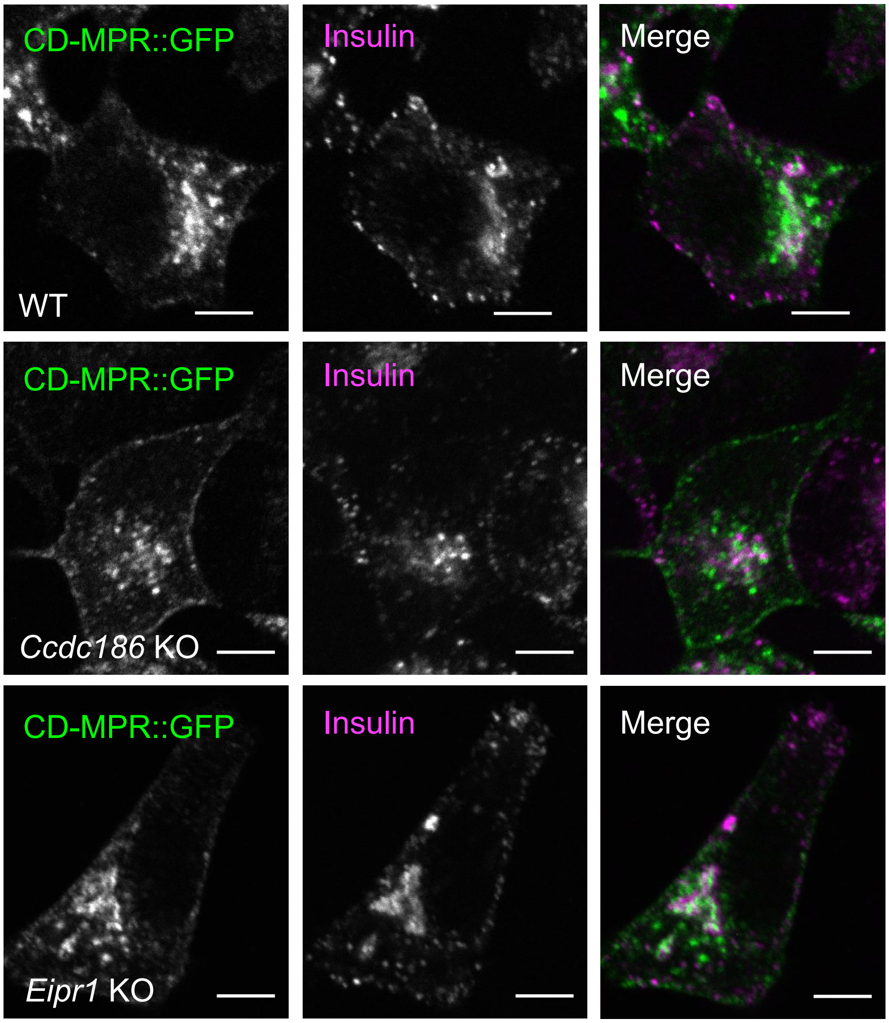
CD-MPR::GFP
Insulin
Merge
WT
CD-MPR::GFP
Insulin
Merge
*Ccdc186* KO
CD-MPR::GFP
Insulin
Merge
*Eipr1* KO

#### TABLES

**Table S1. Top hits from CC3 BioID experiment.**

Unique peptide count organized in the descending order of the difference in number of unique peptides from CC3::BirA*::HA::MAO and the negative control BirA*::HA::MAO. The table shows the hits where the difference was 3 or greater. These data are from the BioID experiment that showed the largest amount of enrichment for CPD. For raw data, see Table S6.

### Table S1

| Protein IDs | Gene names | Peptides BirA*::HA::MAO (control) | Peptides CC3::BirA*::HA::MAO | Peptides CC3::BirA*::HA::MAO minus peptides BirA*::HA::MAO |
|---|---|---|---|---|
| Q9JHW1 | Cpd | 1 | 16 | 15 |
| **A0A0G2K8K1** | ***Ccdc186*** | **2** | **15** | **13** |
| P52873 | Pc | 85 | 94 | 9 |
| F1M9D0;B5DF98 | Map3k2 | 3 | 9 | 6 |
| G3V6W7 | Cpsf3 | 5 | 11 | 6 |
| O08722 | Unc5b | 3 | 9 | 6 |
| B1WBW7 | Traf7 | 0 | 5 | 5 |
| G3V6B0;F1M024 | Pdxdc1 | 9 | 14 | 5 |
| Q9QYF3;P70569;F1M111 | Myo5a | 35 | 39 | 4 |
| P11960 | Bckdha | 6 | 10 | 4 |
| Q5I0C3 | Mccc1 | 32 | 36 | 4 |
| Q63347 | Psmc2 | 13 | 17 | 4 |
| D3ZFX4 | Pgm3 | 2 | 6 | 4 |
| Q9EQH1 | Gab2 | 11 | 15 | 4 |
| Q62807 | Syt17 | 3 | 7 | 4 |
| F1LM47 | Sucla2 | 7 | 11 | 4 |
| B4F775 | Gopc | 0 | 4 | 4 |
| Q62673 | Plk1 | 2 | 6 | 4 |
| D4ACS3 | Mb21d2 | 0 | 4 | 4 |
| Q99P39 | Nfs1 | 3 | 6 | 3 |
| E9PSQ0 | Acacb | 11 | 14 | 3 |
| Q6QDP7;Q66HA2 | Creb3l2 | 4 | 7 | 3 |
| P13437 | Acaa2 | 10 | 13 | 3 |
| Q501R9 | Nbr1 | 6 | 9 | 3 |
| B5DEJ5 | Eefsec | 1 | 4 | 3 |
| P14882 | Pcca | 28 | 31 | 3 |
| F1LT49 | Lrrc47 | 11 | 14 | 3 |
| P68136;P68035;P63269;P62738 | Acta1;Actc1;Actg2;Acta2 | 12 | 15 | 3 |
| D3ZJ50 | Pkp3 | 2 | 5 | 3 |
| P63045;P63025 | Vamp2;Vamp3 | 1 | 4 | 3 |
| D4A017 | Tmem87a | 1 | 4 | 3 |

**Table S2. *C. elegans* strain list**

N2 Bristol wild-type

EG334 *cccp-1(ox334)* III

EG6007 *vps-50(ok2627)* III

EG6939 *eipr-1(tm4790)* I

XZ1924 *vps-50(ok2627)* III *cccp-1(ox334)* III

XZ2108 *eipr-1(tm4790)* I ; *cccp-1(ox334)* III

KG1395 *nuIs183[Punc-129::NLP-21-Venus, Pmyo-2::GFP]* III

EG5258 *nuIs183[Punc-129::NLP-21-Venus, Pmyo-2::GFP]* III *cccp-1(ox334)* III

XZ1055 *eipr-1(tm4790)* I ; *nuIs183[Punc-129::NLP-21-Venus, Pmyo-2::GFP]* III

XZ1607 *nuIs183[Punc-129::NLP-21-Venus, Pmyo-2::GFP]* III *vps-50(ok2627)* III

XZ1918 *nuIs183[Punc-129::NLP-21-Venus, Pmyo-2::GFP]* III *vps-50(ok2627)* III *cccp-1(ox334)* III

XZ2109 *eipr-1(tm4790)* I ; *nuIs183[Punc-129::NLP-21-Venus, Pmyo-2::GFP]* III *cccp-1(ox334)* III

**Table S3. Plasmid list**

| Plasmid name | Vector backbone | Description |
|---|---|---|
| pEGFP-N1 | | GFP expression vector, a gift from Suzanne Hoppins |
| pET50 | pEGFP-N1 | CCDC186::GFP full length, rat CCDC186 cDNA |
| ANF::GFP | pEGFP-N1 | ANF::GFP (rat atrial natriuretic factor fused to GFP, (Hummer et al., 2017) |
| ssGFP | pCAGGS | signal sequence of rat ANF fused to GFP in the chicken actin-based vector pCAGGS (Hummer et al., 2017) |
| pET55 | pEGFP-N1 | GFP::RAB2A, RAB2A cDNA was PCR amplified from 832/13 cDNA library |
| pET54 | pEGFP-N1 | GFP::RAB2B, RAB2B cDNA was PCR amplified from 832/13 cDNA library |
| pJC239 | pCMV-Myc | Myc::CCDC186 |
| pJC271 | pCDNA3.1 | CC3::CCDC186(750-922)::HA::MAO |
| pJC280 | pCDNA3.1 | CC3::CCDC186(759-922)::BirA*::HA::MAO |
| pJJS112 | pCDNA3.1 | BirA*::HA::MAO, a gift from Sean Munro |
| pJC279 | pEGFP-N1 | CD-MPR::GFP, CD-MPR cDNA was PCR amplified from 832/13 cDNA library |
| pPUR | | A gift from Richard Palmiter |
| pBABE-hygro | | A gift from Suzanne Hoppins |
| pSpCas9(BB)-2A-GFP | | PX458 (Addgene) |
| pET221 | pBABE-hygro | CCDC186 in pBABE-hygro |
| pET64 | pSpCas9(BB)-2A-GFP | CCDC186 guide RNA in pSpCas9(BB) |
| pET70 | pPUR | CCDC186 HR (homologous recombination cassette) in pPUR backbone |
| FUGW | | 3rd generation lentiviral plasmid with hUbC-driven EGFP; can be used for cDNA expression (Addgene #14883) |
| psPAX2 | | 2nd generation lentiviral packaging plasmid. Can be used with 2nd or 3rd generation lentiviral vectors and envelope expressing plasmid (Addgene #12259) |
| pVSVG | | Envelope protein for producing lentiviral and MuLV retroviral particles. Use in conjunction with a packaging vector such as pCMV-dR8.2 dvpr (lentiviral) or pUMVC (MuLV retroviral). (Addgene #8454) |
| St6Gal1::pHluorin | FUGW | 17-residue transmembrane domain of beta-galactoside alpha 2,6-sialyltransferase fused to pHluorin. |

**Table S4. Primary antibody list**

| Antibody target | Manufacturer and catalog number | Notes |
|---|---|---|
| CCDC186 | Novus #NBP1-90440 | Rabbit polyclonal, 1:150 for IF and 1:1,000 for WB |
| TGN38 | Sigma #T9826 | Rabbit polyclonal, 1:350 for IF |
| EEA1 | BD Biosciences #610456 | Mouse monoclonal, 1:100 for IF |
| GM130 | BD Biosciences #610822 | Mouse monoclonal, 1:100 for IF, but works at 1:300 |
| ERGIC53 | Santa Cruz #sc-398893 | Mouse monoclonal, 1:50 for IF |
| LAMP1 | DHSB | Mouse monoclonal, 1:50 for IF |
| GFP | Santa Cruz #sc-9996 | Mouse monoclonal, 1:200 to 1:400 for IF, 1:1,000 for WB |
| GFP | Santa Cruz #sc-8334 | Rabbit polyclonal, 1:200 to 1:400 for IF, 1:1,000 for WB |
| Insulin | Sigma #I2018 | Mouse monoclonal, 1:400 for IF |
| Proinsulin | Abcam #ab8301 | Mouse monoclonal, 1:200 for IF |
| Syntaxin 6 | Abcam #ab12370 | Mouse monoclonal, 1:100 for IF |
| TGN38 | Novus #NB300-575S | Mouse monoclonal, 1:500 for IF |
| CgA | Santa Cruz #sc-393941 | Mouse monoclonal, 1:200 for IF |
| Myc | Santa Cruz sc-40 | Mouse monoclonal, 1:200 for IF, 1:1,000 for WB |
| GFP | Roche #11814460001 | Mouse monoclonal, 1:1,000 for WB |
| Beta-tubulin | DHSB E7 | Mouse monoclonal, 1:1,000 for WB |
| Beta-tubulin | ThermoFisher , BT7R, #MA5-16308 | Mouse monoclonal, 1:1,000 for WB |
| HA | Roche #11867423001 | Rat monoclonal, 1:150 for IF and 1:1,000 for WB |
| CPD* | A gift from Lloyd Fricker AE160 | Rabbit polyclonal, 1:200 for IF and 1:1,000 for WB |
| PCSK1 (PC1/3) | Sigma SAB1100415 | Rabbit polyclonal, 1:1,000 for WB |
| CPE-N terminus** | A gift from Lloyd Fricker | Rabbit polyclonal, 1:1,000 for WB, 1:100 for IF |
| Pro-CPE form*** | A gift from Lloyd Fricker | Rabbit polyclonal, 1:1,000 for WB, 1:100 for IF |
| CPE-C terminus**** | A gift from Lloyd Fricker | Rabbit polyclonal, 1:1,000 for WB, 1:100 for IF |

* Antiserum was generated to the C-terminal 57 amino acids of mammalian CPD (cytosolic tail, (Song and Fricker, 1996).

** Antiserum was raised to the N-terminal 15 amino acids of bovine CPE (Fricker et al., 1990).

*** Antiserum to pro-CPE was generated to the peptide QEPGAPAAGMRRC coupled to maleimide-activated keyhole limpet hemocyanin (KLH). This peptide corresponds to the 12 residues of mouse/rat/human pro-CPE with an added Cys on the C-term (for coupling to the carrier protein, KLH).

**** Antiserum was raised against the 9-residue peptide KMMSETLNF corresponding to the C terminus of mouse CPE (Fricker et al., 1996).

**Table S5. Secondary antibody list**

| Antibody target | Manufacturer and catalog number | Notes |
|---|---|---|
| Anti-mouse Alexa488 | Jackson immunoresearch #115-545-146 | 1:1000 for IF |
| Anti-mouse Rhodamine Rex-X | Jackson immunoresearch #715-295-150 | 1:1000 for IF |
| Anti-rabbit Alexa488 | Jackson immunoresearch #711-545-152 | 1:1000 for IF, (1:100 for STED microscopy) |
| Anti-rabbit Rhodamine TRITC | Jackson immunoresearch #111-025-144 | 1:1000 for IF |
| Anti-rat Dylight550 | ThermoFisher #SA5-10027 | Highly cross-adsorbed, 1:200 to 1:500 for IF |
| Anti-rabbit Atto647 | A gift from Joshua Vaughan's lab | 1:200 for IF |
| Anti-mouse Dylight488 | ThermoFisher #SA5-10166 | Highly cross-adsorbed, used at 1:200 to 1:500 |
| Anti-mouse Alexa555 | A gift from Joshua Vaughan's lab | Used at 1:10 for STED microscopy |
| Goat anti-rabbit Alexa680 | Jackson immunoresearch #111-625-144 | 1:20,000 for western |
| Goat anti-mouse Alexa680 | Jackson immunoresearch #115-625-166 | 1:20,000 for western |
| Donkey anti-mouse Alexa790 | Jackson immunoresearch #715-655-150 | 1:20,000 for western |
| Alexa790 Streptavidin | Jackson immunoresearch #016-650-084 | 1:10,000 for western |

**Table S6. Raw mass-spec data for CC3 BioID experiment.**

The table contains all the data for the experiment shown in Table S1.

**Table S7. LFQ intensities for CC3 BioID experiment.**

The table contains Z-score normalized log2 LFQ intensities from MaxQuant output, Student's t-test p-values and ratios (CC3::BirA*::HA::MAO vs BirA*::HA::MAO). For calculation of p-values and ratios, see Materials and Methods.
